# Triplet tumbling microscopy enables in situ quantification of protein complex assembly and dynamics

**DOI:** 10.64898/2026.05.07.723557

**Authors:** Julia R. Lazzari-Dean, Alfred Millett-Sikking, Prashant Rao, Zena D. Jensvold, Sidharth Kashyap, Hannah Baddock, Maria Ingaramo, Aaron H. Nile, Andrew G. York, Magdalena Preciado López

## Abstract

Protein-protein interactions (PPIs) mediate diverse cellular processes but are typically characterized in vitro with recombinant proteins. Current cell-compatible approaches for measuring PPIs have limited scope and often require prior knowledge of the interacting partners. Here, we develop triplet tumbling microscopy (TTM) to reveal PPIs in cells in real time. TTM reports protein complex size from rotational diffusion (“tumbling”) by leveraging infrared-triggerable emission from triplet states. These long-lived triplets overcome the size limitations of traditional fluorescence anisotropy, enabling TTM to measure species from small protein complexes to organelle-scale beads. We apply TTM to detect PPIs, quantify bound fractions, and distinguish protein complexes by size. We measure diverse interactions, including rapamycin-induced dimerization, p53 homo-oligomerization, and binding of the E3-ligase E6AP to the human papilloma virus 16 E6 protein. TTM hardware is compatible with most fluorescence microscopes, providing a versatile way to access molecular insights within cells.

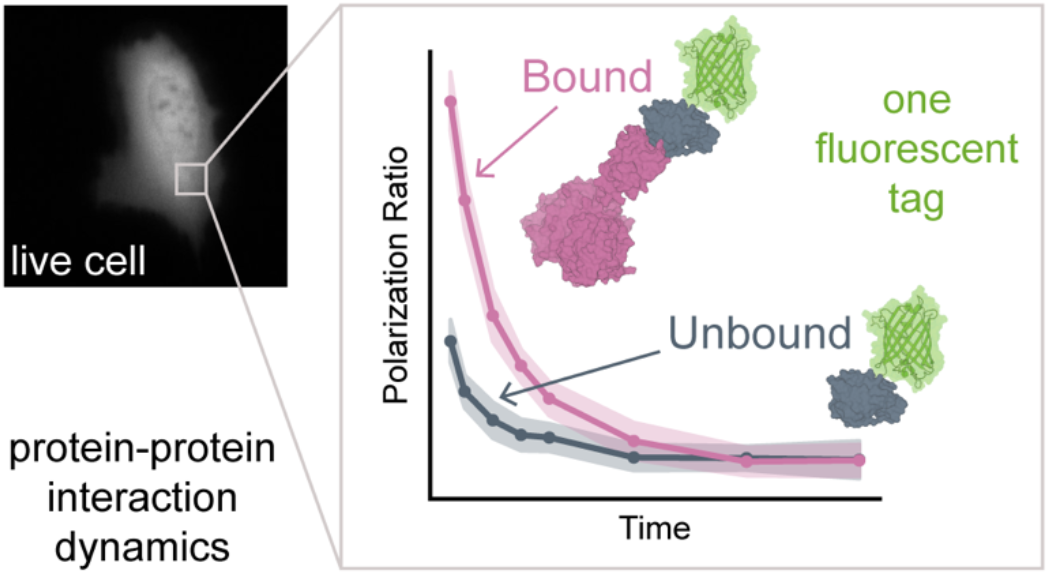

## Introduction

Protein function is often determined by interactions: association with regulatory domains, oligomerization into activated complexes, or transient connections as part of a signal transduction cascade. Gold standard methods for protein-protein interaction (PPI) characterization are typically biochemical and biophysical, performed on pure protein or cell lysate (e.g. surface plasmon resonance, co-immunoprecipitation, crosslinking, structure determination).^1^ Although these approaches can detect PPIs, they cannot capture the spatiotemporal patterning of interactions that underlies cellular function. Techniques that can monitor and quantify PPIs in situ are essential for linking intermolecular interactions to the cellular phenotypes they produce.

Fluorescence microscopy is routinely used to map the spatiotemporal distribution of proteins in living cells, tissues, and animals, but it does not reveal protein-protein interactions (PPIs). To address this limitation, several complementary techniques are commonly used; they fall into three main categories. First, Förster resonance energy transfer (FRET) can report if two fluorophores are within 10 nm of each other, but it does not easily extend to multi-component interactions, and it requires two appropriately positioned fluorescent tags.^2^ Second, translational diffusion-based measurements, such as fluorescence correlation spectroscopy^3^ or single particle tracking,^4^ report particle size, with size changes used as a proxy for PPIs. However, translational diffusion is weakly dependent on size (scaling with the cube root of volume),^5^ and these methods can suffer from photobleaching artifacts and impose strict requirements on label density. Third, rotational diffusion (“tumbling”) measurements, such as fluorescence anisotropy, report particle size with high sensitivity (scaling linearly with volume),^6^ but they are traditionally restricted to small targets (<50 kDa).

Because fluorescence anisotropy is a sensitive, scalable, and versatile technique, we sought to address its primary limitation: the restricted size range it can detect. The size limitation of fluorescence anisotropy originates from the short fluorescence lifetime of typical fluorophores (<5 ns). In anisotropy measurements, polarized light first photoselects an aligned subpopulation of fluorophores from the sample. If these fluorophores are immobilized or very large, they tumble slowly, and the emission remains polarized. If the fluorophores are instead very small, tumbling is fast, randomizing the fluorophores’ orientations and resulting in depolarized emission. Intermediate rates of tumbling produce intermediate levels of polarization, from which particle size can be inferred (**Fig. 1a**). Unfortunately, for most protein complexes (>50 kDa), tumbling is negligible over the nanosecond-scale fluorescence lifetime, so the emission remains polarized (i.e. unchanged) and is not sensitive to binding.

**Figure 1.**
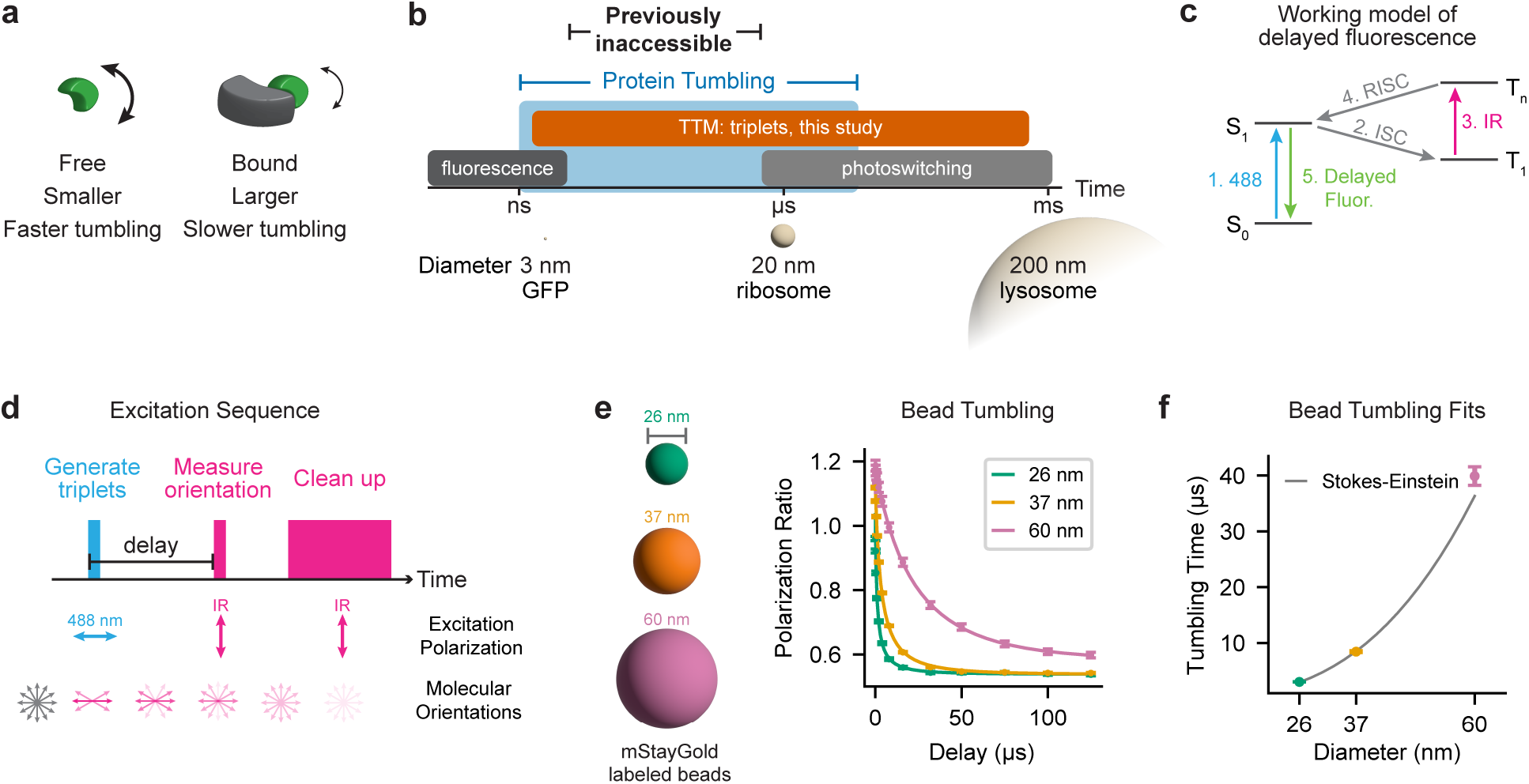
Triplet tumbling microscopy (TTM) approach and validation with fluorescent beads. (**a**) Tumbling rate depends on particle size. Binding increases the hydrodynamic radius of the labeled species and slows tumbling. (**b**) Protein tumbling occurs on the ∼50 ns to the hundreds of μs timescale. This timescale is not fully captured within the lifetime of fluorescence or reversible switchable fluorescent proteins^7^ (photoswitching), but these timescales are well matched with triplet lifetimes. Corresponding particle diameters and example biomolecules for each timescale are included below. (**c**) Working model of delayed fluorescence, as described previously.^11^ Initial absorption of 488 nm light excites ground state singlets (S0) to the excited singlet state (S1). Triplets (T1) spontaneously form via intersystem crossing (ISC) from S1. Subsequent infrared (IR) illumination generates excited triplets (Tn), which exhibit reverse intersystem crossing (RISC) back to S1. These re-generated excited singlets then emit delayed fluorescence (fluor.). (**d**) TTM excitation sequence. A 488 nm light pulse (blue, parallel) generates triplet states, followed by an infrared pulse (magenta, perpendicular) that measures orientation by triggering delayed fluorescence. A second infrared pulse cleans up triplets that were not triggered by the first pulse. The detector is only active during the first infrared pulse and is otherwise off, rejecting fluorescence generated by the 488 nm or cleanup pulses. Delays are measured from the start of the triplet generation pulse to the start of the first infrared pulse. (**e**) Tumbling curves for mStayGold-labeled latex beads of known diameter. Points represent mean ± s.d. of n=3 measurements, and solid line indicates the exponential fit. (**f**) Tumbling times from the fits in (**e**), shown with predicted values from the Stokes-Einstein-Debye equation^17^ (gray line), assuming viscosity 30% higher than water.

Tumbling of cellular protein complexes occurs on the time scale of hundreds of nanoseconds or longer^7^ (**Fig. 1b**). Successful use of tumbling to measure PPIs requires a long-lived state that can report particle orientation over these longer timescales. To address this, previous work explored dye phosphorescence to extend the time range,^8–10^ but phosphorescence is typically dim, and dyes with high phosphorescence yields can be photosensitizers. In an alternative approach, reversible switchable fluorescent proteins were recently used to track microsecond-scale tumbling, but they exhibit low signal for sub-microsecond measurements.^7^ Building on this work, we and others have shown that delayed, infrared-triggerable fluorescence of fluorescent proteins^11^ can be used for microsecond-scale tumbling measurements. This “optically activated delayed fluorescence” (OADF) is believed to result from long-lived triplet states (**Fig. 1c**).

Triplets form spontaneously as a byproduct of fluorescence excitation; in fluorescent proteins, triplets can persist for milliseconds.^12^ Infrared light (∼900 nm for GFP) triggers triplets to return to an excited singlet state, which emits delayed fluorescence.^11,13^ Although the triplet lifetime matches protein tumbling timescales well, previous demonstrations of triplet tumbling were limited to large particles such as viruses or beads and, to the best of our knowledge, have not been applied to protein complexes in cells.^14–16^

Here, we develop triplet tumbling microscopy (TTM) to monitor PPIs in living cells. We describe a custom widefield fluorescence TTM instrument and rigid fluorescent protein tags with improved dynamic range. Using a rapamycin-induced dimerization system, we demonstrate that TTM quantifies protein complex size, bound fraction, and interaction dynamics on the timescale of seconds. We then apply TTM to measure homo-oligomerization of the tumor suppressor protein p53, thereby extending beyond simple heterodimerization interactions. Finally, we assess TTM sensitivity by resolving the small (∼10%) change in size associated with 16E6 binding the E3-ligase E6AP. Together, these results establish TTM as a broadly applicable approach for quantitatively assessing PPIs in living cells.

## Results

### Instrumentation for triplet tumbling microscopy

To enable triplet-based tumbling measurements of protein complexes in cells, we developed a pulse sequence to record the delayed fluorescence with nanosecond time resolution. To concentrate the limited signal at the most informative time points, we used triplet generation and detection pulses (**Fig. 1d**) rather than continuous illumination as previously demonstrated.^14^ The sequence is as follows: we first excite the sample with a pulse of linearly polarized 488 nm light, generating both prompt (conventional) fluorescence and triplets. For each absorption event, the probability of generating a triplet is low (approximately 0.3% for YFP^11^); the triplet fraction can be increased over multiple absorption events during the 30-100 ns excitation pulse. These triplet state fluorophores are initially aligned with the 488 laser’s polarization but tumble over time, scrambling their orientations. Following a variable delay (75 ns to hundreds of microseconds), we then gate the camera on and apply an infrared trigger pulse (785, 830, 915, and/or 940 nm) to measure the orientation distribution (i.e. tumbling). Lastly, to avoid carryover of triplets between cycles, we include a longer infrared “cleanup” pulse after each probe pulse to trigger any remaining triplets. Because the triggered emission signal is relatively weak, we integrate thousands of cycles per measurement (total acquisition time of a few hundred milliseconds, **Supplementary Table 1**), which greatly improves the signal-to-noise ratio.

Because the excitation and detection scheme described above is not typically supported by commercial microscope software, we built a custom widefield microscope for TTM (**Supplementary Fig. 1**). In our system, we generated 30-100 ns excitation pulses by digitally modulating the diode lasers. To maximize the available intensity, we focused the laser output to an 8 μm (1/e^2^) diameter spot for tumbling measurements. Critically, we used the fast-gating capability of an intensified camera to reject the prompt fluorescence generated by the 488 nm pulse and selectively detect delayed photons from the IR trigger. We confirmed that both the 488 and IR pulses were required for generation of delayed fluorescence, ruling out a contribution from spontaneous delayed emission (**Supplementary Fig. 2**). We split the emission with a polarizing beam splitter and then recorded the parallel (aligned with 488 beam) and perpendicularly (aligned with IR beam) polarized emission images side-by-side on the intensified camera. Cell finding and focusing were performed with a separate, LED-illuminated beam path (**Supplementary Fig. 1**).

To analyze TTM data, we calculate the polarization ratio by dividing the parallel signal by the perpendicular signal. When we measure the polarization ratio at different delays, we obtain a tumbling curve. We fit this tumbling curve to an exponential decay model to obtain its characteristic tumbling time, which should be proportional to the size of the fluorescent particles. Our analysis approach is described further in **Supplementary Note 1**.

To validate that the tumbling time correlates with particle size, we measured tumbling curves for fluorescent latex beads across a range of diameters (**Fig. 1e,f**). Beads were made fluorescent by coating their surface with purified mStayGold^18^ proteins. Using delays ranging from 150 ns to 125 μs, we observe tumbling times proportional to bead size, in good agreement with predictions from the Stokes-Einstein-Debye equation^17^ (**Supplementary Table 2**).

### Triplet tumbling reports PPIs in a purified protein system

To identify fluorescent protein tags suitable for TTM, we first surveyed the triplet signal across a variety of purified green fluorescent proteins (**Supplementary Fig. 3, Supplementary Fig. 4**). For reference, 60 nm diameter beads (**Fig. 1e**), which are the size of small lysosomes, fully scramble their orientations over a 30 μs timescale; therefore, a lifetime > 50 μs is sufficient to resolve very large protein assemblies. Most proteins tested had an IR triggerable triplet signal, with triplet lifetimes ranging from 100 μs to 5 ms. Interestingly, mStayGold and mBaoJin,^19^ both recently developed for their photostability, had two of the shortest triplet lifetimes. From the proteins tested, we selected mVenus because it provided the highest triggerable triplet signal relative to the prompt fluorescence; we also included mStayGold for its higher total photon budget.

We sought to demonstrate that TTM can monitor PPIs in a reconstituted recombinant system. To this end, we created a size ladder from the rapamycin-inducible dimerization of FK506-binding protein (FKBP) and the rapamycin binding domain of mTOR (FRB).^20^ We fused FKBP to a fluorescent protein and FRB to binding partners of varying molecular weight (**Fig. 2a**, Penn State protein ladder:^21^ FRB, SMT3-FRB, MBP-FRB). We affinity purified 6xHis-mVenus-FKBP or 6xHis-mStayGold-FKBP and nonfluorescent FRB fusions of varying sizes (**Supplementary Fig. 4**) with 6xHis and Strep tags. A critical design element of a tumbling tag is its rigidity, i.e. the rotation of the tag must track the rotation of the target. Flexible regions between the tag and the protein of interest reduce the dynamic range.^7^ To avoid the flexible linker sequences commonly used at fluorescent protein termini, we tested how progressive truncations of the mVenus C-terminus affect the tumbling signal (**Supplementary Fig. 5**). As previously reported for actin fusions,^22^ we found that removing the flexible residues up to the ß-barrel was well tolerated and increased the dynamic range. We truncated mStayGold after the ß-barrel to the same number of residues that was optimal for mVenus. For the mStayGold-FKBP and MBP-FRB interacting pair, we used surface plasmon resonance (SPR) to verify that the addition of bulky tags did not disrupt the interaction. Using a two-state reaction model, we measured an apparent Kd of 13 nM for MBP-FRB binding to pre-formed mStayGold-FKBP:rapamycin complexes (**Supplementary Fig. 4**), consistent with literature reports for FRB and FKBP:rapamycin alone (Kd of 12 nM^20^).

**Figure 2.**
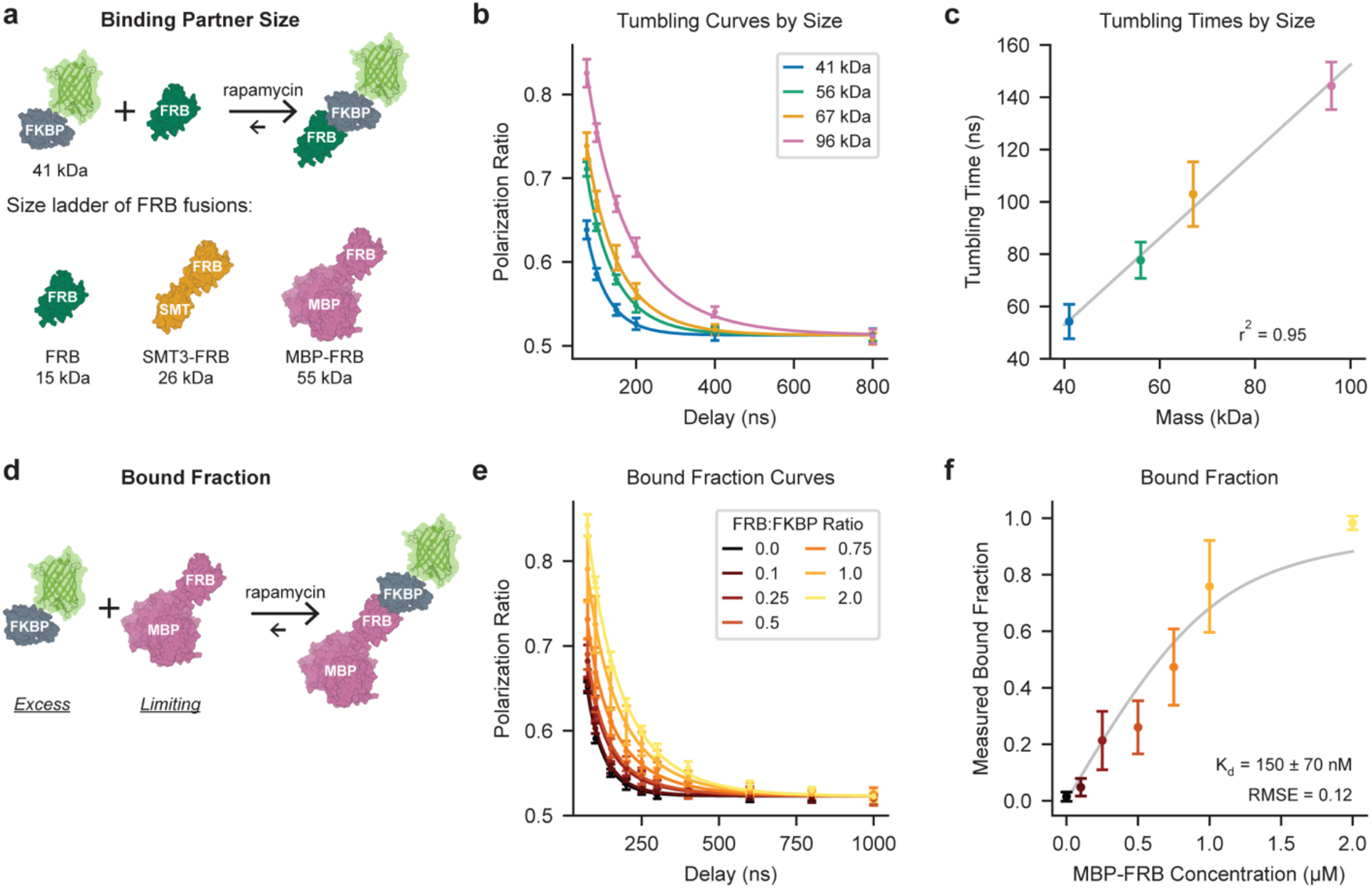
Triplet tumbling microscopy reveals complex size and bound fraction in purified protein mixtures. (**a**) Schematic of the rapamycin-inducible FKBP:FRB dimerization system. Fluorescent protein-tagged FKBP binds nonfluorescent FRB only in the presence of rapamycin. Fusing FRB to partners of increasing mass yields a “size ladder” upon rapamycin addition. Protein cartoons are based on PDB IDs 8ER6, 1AUE, 1EUV, 1MPD, and 8BXT. (**b**) Tumbling curves measured from samples in (a), either before rapamycin addition (mStayGold-FKBP alone, 41 kDa, blue) or after rapamycin-induced dimerization with an FRB binding partner. Solid line indicates the fit. (**c**) Tumbling times from fitting the data in (b) are linearly proportional to the predicted complex mass. A line of best fit is shown in light gray. (**d**) Schematic for generating mixtures with excess labeled component (mStayGold-FKBP) relative to unlabeled binding partner (MBP-FRB). (**e**) Tumbling curves for incompletely complexed mStayGold-FKBP (total concentration 1 µM) after addition of 2 µM rapamycin in the presence of limiting MBP-FRB. (**f**) Bound fraction extracted from curves in (e) by double exponential fitting. Gray line indicates fit to the Morrison equation^23^ to determine dissociation constant (± 95% c.i.). Error bars show the mean ± s.d. across n=9-12 (b, c) or n=6 (e, f) replicates with rapamycin; recordings without rapamycin were pooled across all samples. All measurements were done in solutions containing 20% glycerol to approximate cellular viscosity.^24,25^

We recorded tumbling curves from mixtures of recombinant mStayGold-FKBP and an FRB binding partner before and after rapamycin addition. Rapamycin addition slowed tumbling, and the magnitude of the tumbling change increased with larger FRB binding partners (**Fig. 2b**).

These tumbling curves were well described by a single exponential decay. As expected for approximately spherical particles,^17^ the relationship between the tumbling time and the expected molecular weight of the complex was linear (**Fig. 2c**). We also observed a linear relationship between mass and tumbling time with fluorescently tagged FRB and an unlabeled FKBP (i.e. swapped tags, **Supplementary Fig. 6**), highlighting the robustness of our approach.

By varying the stoichiometry of the labeled and unlabeled components, we can generate mixtures containing free and MBP-FRB-bound mStayGold-FKBP. As expected, sub-stoichiometric MBP-FRB leads to tumbling curves that are intermediate between the fully unbound and fully bound states (**Fig. 2e**). We modeled this rotational diffusion as a sum of two exponentials and calculated the bound fraction from the ratio of the two exponentials’ amplitudes. We used this to optically estimate an effective Kd of 150 ± 70 nM (95% c.i.) (**Fig. 2f**). Our optical endpoint measurement is analogous to a steady-state analysis of SPR data, from which we obtain an effective Kd of 89 nM (**Supplementary Fig. 4**), in good agreement with the TTM estimate.^20^

### Extending triplet tumbling microscopy to living cells

Encouraged by our in vitro results, we transferred the rapamycin-inducible system to live U2OS cells (**Fig. 3a**). To generate an excess of unlabeled partner relative to mStayGold-FKBP, we placed the unlabeled component upstream of an internal ribosome entry site (IRES) and mStayGold-FKBP downstream. As in the purified protein system, we generated a size ladder of the unlabeled component by fusing FRB to SMT3, MBP, and MBP-pepN (**Fig. 3b**).

**Figure 3.**
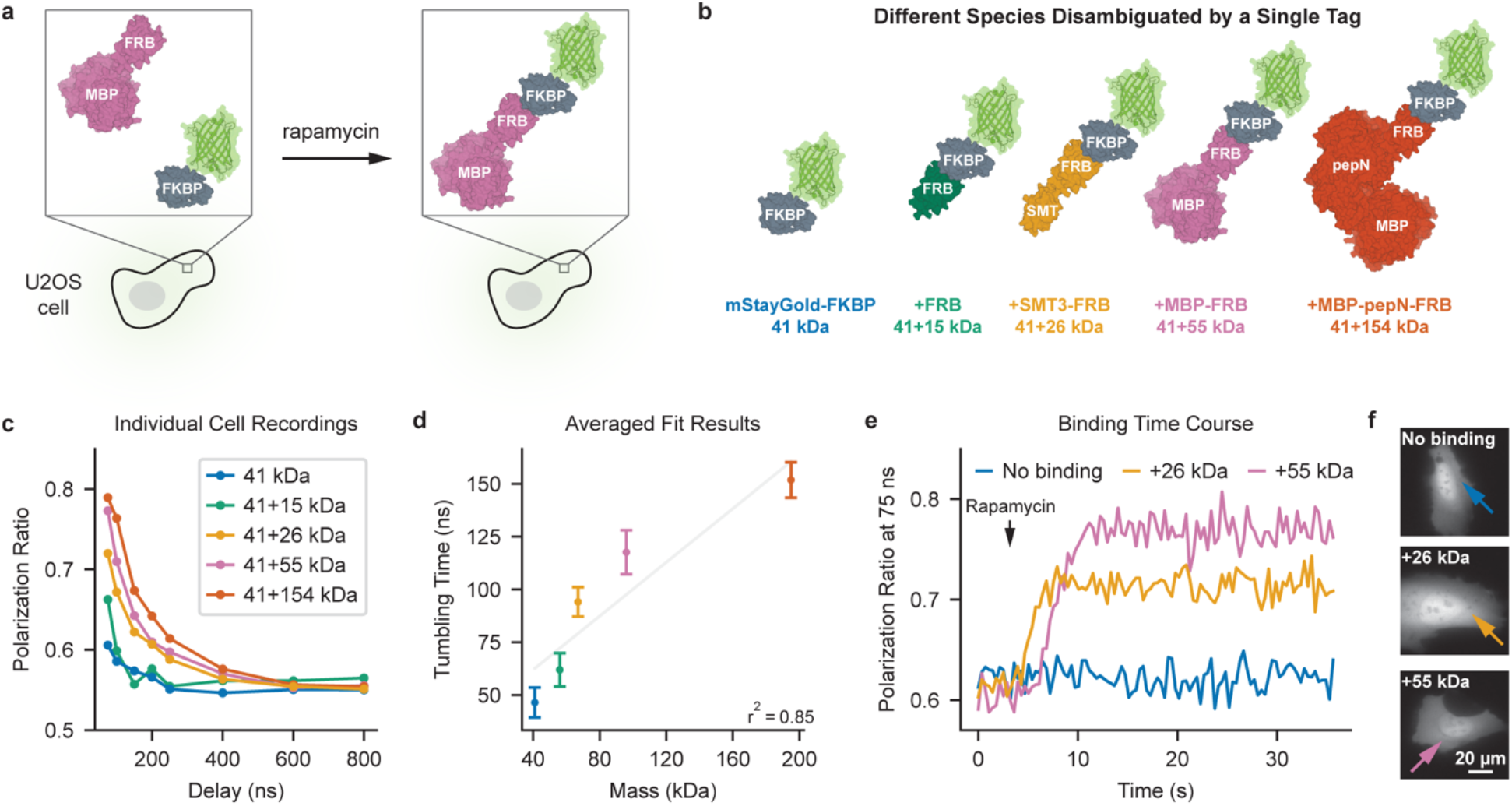
Triplet tumbling microscopy reveals protein complex formation and dynamics in cells. (**a**) Schematic of the rapamycin-induced dimerization system expressed in U2OS cells. (**b**) Four constructs were generated, each expressing mStayGold-FKBP downstream of an unlabeled FRB binding partner via an IRES. FRB binding partners span molecular weights from 15 to 154 kDa. Cells were transfected with a plasmid encoding one of the four binding partners. PDB IDs for protein cartoons: 1AUE, 1EUV, 1MPD, 2HPT, 8BXT, 8ER6. (**c**) Representative tumbling curves measured from individual cells treated with DMSO control (blue, 41 kDa) or 10 µM rapamycin (other colors) for 1 hour. (**d**) Tumbling times derived from fits to individual recordings. Light gray line indicates the best linear fit across masses. Points represent mean ± s.d. of n=11-15 cells; untreated cells were pooled across constructs (n=54). (**e**) Time course of the polarization ratio at 75 ns while DMSO (no binding, blue) or 10 µM rapamycin (yellow, pink) was added to a cell expressing the rapamycin-induced dimerization system. Black arrowhead indicates time of DMSO or rapamycin addition. (**f**) Widefield fluorescence images of the cells in (e). Colored arrows indicate the center of the ∼8 µm (1/e^2^) recording location.

All FRB fusions tested exhibited similar localization and tumbling prior to rapamycin treatment. Each recording was made from an 8 µm (1/e^2^) diameter spot in the cytosol of an individual cell (**Fig. 3c**). After rapamycin treatment, tumbling slowed in a size-dependent manner, as revealed by the tumbling times obtained from single exponential fitting (**Fig. 3d**). The tumbling times measured in cells exhibited a similar trend to those measured in vitro (**Fig. 2c**). Tumbling in cells was slightly faster than with recombinant proteins, likely reflecting viscosity differences between the cytoplasm and the 20% glycerol solutions used in vitro. We observe a linear relationship between molecular weight and tumbling time, with some nonlinearity for the largest fusions tested, perhaps attributable to segmental motion. Beyond population averages, single-recording tumbling curves in cells were sufficient to distinguish complex sizes (**Fig. 3c**). FRB alone constructs showed overall ∼10-fold lower expression based on mStayGold intensity, yet we were still able to infer binding state. We observed similar patterns in mVenus-tagged versions of these constructs (**Supplementary Fig. 7**), suggesting that these results are generalizable across fluorescent protein tags.

Having demonstrated that TTM can resolve PPIs in living cells, we next evaluated whether TTM can track binding dynamics. To explore this, we added rapamycin to cells while acquiring TTM images in real time and observed a marked increase in polarization ratio upon dimerization (**Fig. 3e,f**, **Supplementary Fig. 8**), consistent with larger complexes. For endpoint measurements, we would typically acquire a full tumbling curve at each experimental timepoint. However, the fast dynamics of rapamycin-induced dimerization^20^ necessitated faster imaging speeds than were possible with multi-delay acquisitions. To achieve an imaging speed of ∼3 Hz, we took advantage of the flexibility of our pulsed excitation scheme, recording one delay (75 ns) at 100 experimental timepoints. Because the 41+26 kDa and 41+55 kDa complexes have different polarization ratios at 75 ns, this single delay approach allowed us to differentiate between the two final protein complex sizes formed. Although we observed some photobleaching during the recording (**Supplementary Fig. 9**), the polarization ratio remained stable throughout.

### TTM reveals p53 homo-oligomerization in cells

Having demonstrated the power of TTM in a model system, we next asked whether TTM could report a physiologically relevant PPI of a different type: homo-oligomerization of the tumor suppressor protein p53. In cells, p53 is reported to exist as a mixture of monomers, homodimers, and homo-tetramers (dimer of dimers), depending on its activation state^26,27^ (**Fig. 4a**). Tetramerization is required for DNA binding and activation of downstream tumor suppressor pathways;^26^ mutations in the tetramerization domain contribute to oncogenesis.

**Figure 4.**
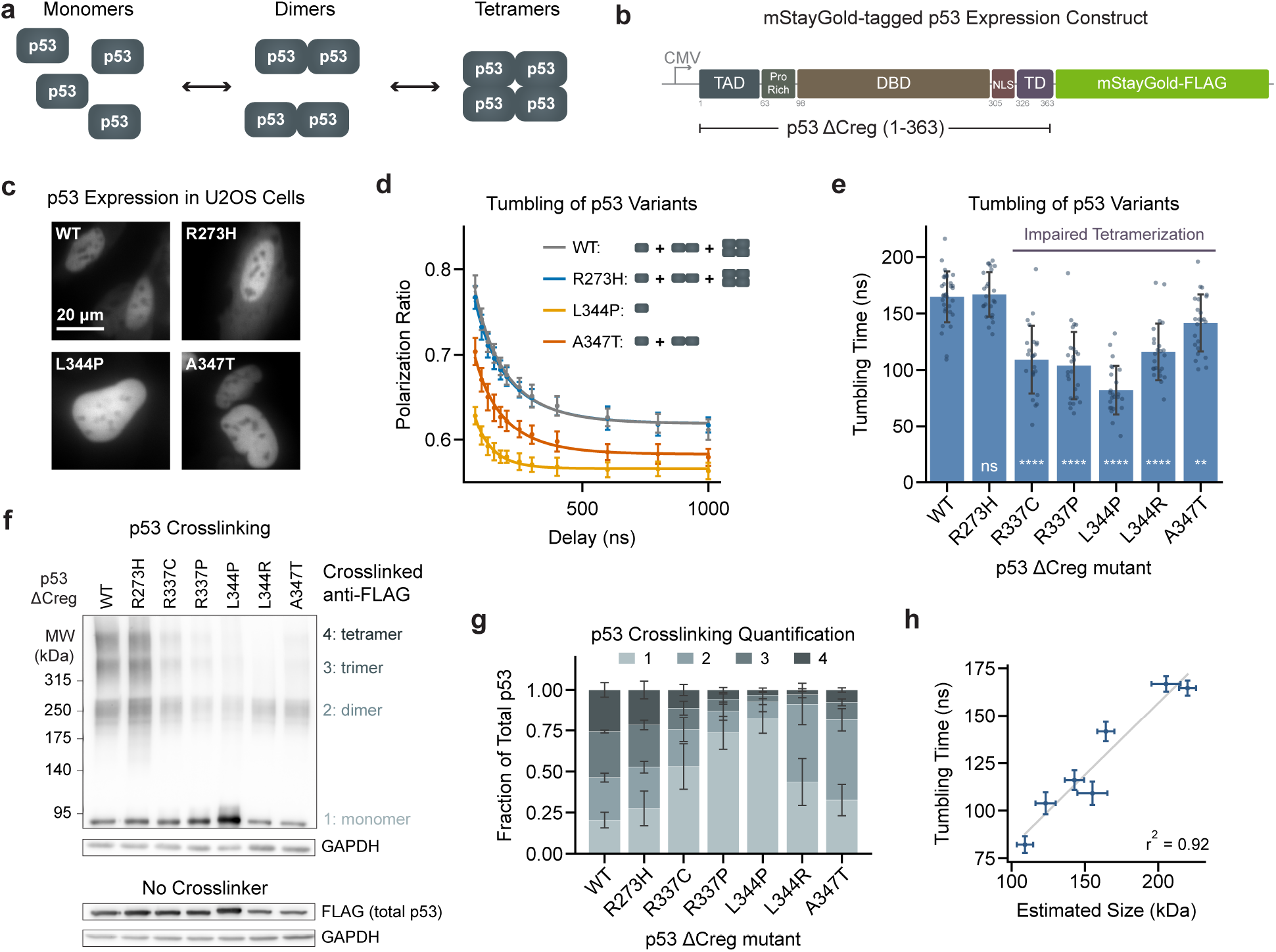
TTM reveals homo-oligomerization of wild type and mutant p53 in cells. (**a**) Schematic of p53 monomeric, dimeric, and tetrameric states. (**b**) Domain architecture of p53 expression constructs lacking the C-terminal regulatory domain. TAD, transactivation domain; DBD, DNA binding domain; NLS, nuclear localization sequence; TD, tetramerization domain. (**c**) Representative widefield fluorescence images of U2OS cells transiently expressing p53 variants. (**d**) Averaged tumbling curves for p53 expressed in U2OS cells. Solid line indicates fit to a single exponential rotational diffusion model. Error bars are mean ± s.d. (n=24-32 cells, ∼8 µm 1/e^2^ diameter recording spot in the nucleus of each cell). (**e**) Tumbling times fit on a per-cell basis for the data in (d). Asterisks indicate p value for Dunnett’s post hoc test comparing each mutant with the wild type, following one-way ANOVA (F(6, 171) = 48, p<0.0001). Points indicate individual cell recordings; error bars are mean ± s.d. (**f**) Representative Western blot of p53-expressing U2OS lysates, with (top) and without (bottom) DSG crosslinker. Tentative band assignments are next to the blot; apparent trimers likely result from incomplete crosslinking. (**g**) Quantification of relative band intensity on the crosslinked Western blots (**Supplementary Fig. 11**). Error bars are mean ± s.d. of n=4 blots. (**h**) Correlation between tumbling time in (e) and weighted average size in (g), with error bars as mean ± s.e.m.

Cellular p53 interactions are multifaceted: in addition to homo-oligomerization, p53 binds numerous regulatory proteins and DNA, and it is the target of various post-translational modifications. To simplify the system, we used a truncated p53 lacking the C-terminal regulatory region (Creg, final 30 amino acids, **Fig. 4b**). Similar truncations have shown reduced DNA binding and reduced interaction with regulatory proteins.^28^ We fused mStayGold to the C-terminus of p53, as C-terminal modifications have been shown to preserve p53 dynamics.^29^ To test how p53 oligomerization state affects tumbling, we generated nine individual point mutations: five in the tetramerization domain (TD: R337C, R337P, L344P, L344R, A347T) and four in the DNA binding domain (DBD: V143A, R175H, Y220C, R273H) (**Fig. 4b, Supplementary Fig. 10**).

We transiently overexpressed the above constructs in U2OS cells, observing mStayGold fluorescence for all constructs. Wild type p53 and most mutants primarily localized to the nucleus, although three DBD N-terminal mutations (V143A, R175H, Y220C) resulted in cytosolic localization (**Fig. 4c, Supplementary Fig. 10**), which has been previously observed for mutant p53.^30^ All constructs migrated at the expected molecular weight by Western blot (**Supplementary Fig. 11**). Expression levels varied between constructs and between cells; for tumbling recordings, we selected cells with similar, intermediate fluorescence levels.

With this system in hand, we applied TTM to quantify how p53 complex size varied across our mutant series in live cells. Based on the corrected Akaike information criteria,^31^ the single cell tumbling curves were best described by a single exponential decay model (**Supplementary Note 1**, **Fig. 4d,e**). As expected, we observed longer tumbling times for wild type p53 and for mutations outside of the tetramerization domain, consistent with a larger complex size.

Interestingly, for non-tetramerization domain mutants, tumbling time did not depend strongly on whether fluorescence was localized to the cytoplasm or the nucleus (**Supplementary Fig. 10**). Mutations in the tetramerization domain resulted in shorter tumbling times, consistent with smaller complex sizes. Consistent with previous reports on p53 interactions,^32^ L344P promotes a monomeric distribution and has a short tumbling time, whereas A347T favors a dimeric state and has an intermediate tumbling time.

To validate the TTM results, we applied chemical crosslinking to preserve p53 interactions for analysis by Western blot (WB). In crosslinked lysates, p53 constructs with wild type tetramerization domains were predominantly higher molecular weight species; we tentatively assigned bands as monomers, dimers, trimers, and tetramers. In agreement with TTM results, TD mutants had stronger monomer bands and reduced higher molecular weight banding (**Fig. 4f,g, Supplementary Fig. 11**). Fluorescence-detected size exclusion chromatography (FSEC)^33^ on uncrosslinked lysates showed monomer, dimer, and tetramer populations (**Supplementary Fig. 12**), suggesting that the trimer band is the product of incomplete crosslinking. Because crosslinking captures interactions without lysis or dilution of sample (unlike FSEC), we used the crosslinked WB size results as a benchmark for TTM. From the relative band intensities on the crosslinked WB, we estimated a weighted average p53 complex size for each mutant, and we observed good agreement between crosslinked size and tumbling time (**Fig. 4h**).

We observed different polarization ratios at long times for different p53 mutants (**Supplementary Fig. 10**), which we did not expect from tumbling alone. We posited that this offset was attributable to homo-FRET between mStayGold tags in oligomerized p53. It is likely that, in dimers and tetramers, homo-FRET partially scrambles polarization before significant tumbling occurs, resulting in a lower dynamic range and shifted polarization ratios. In monomeric constructs, this effect is removed (**Supplementary Fig. 10**). To test this interpretation, we measured homo-FRET with conventional fluorescence anisotropy, observing moderate fluorescence depolarization over the first few nanoseconds in tetramerization-competent constructs (**Supplementary Fig. 13**). Nevertheless, because homo-FRET occurs on much faster timescales than tumbling, meaningful tumbling times can still be extracted in samples with mild or moderate homo-FRET (**Fig. 4h**).

To confirm that our results were not restricted to truncated p53 constructs, we compared TTM results from the full length p53 (**Supplementary Fig. 14**) wild type and the L344P tetramerization domain mutant. In full length p53, as with the truncation constructs (**Fig. 4**), we observed shorter tumbling times in the L344P mutant. In the crosslinking assay, the L344P mutant had a strong monomer band and minimal higher molecular weight banding (**Supplementary Fig. 11**), corroborating the TTM result.

### TTM can reveal small size changes in cells

To demonstrate TTM’s sensitivity to small size changes, we next examined binding of the E3-ligase E6AP to the HPV16 E6 protein (16E6). This high-affinity interaction^34^ between a host E3-ligase and viral protein leads to degradation of the tumor suppressor protein p53 and is a primary driver of HPV-induced oncogenesis^35^ (**Fig. 5a**). Obtaining a robust cellular readout for the 16E6:E6AP interaction would therefore be valuable for the development of therapeutics that disrupt this complex.

**Figure 5.**
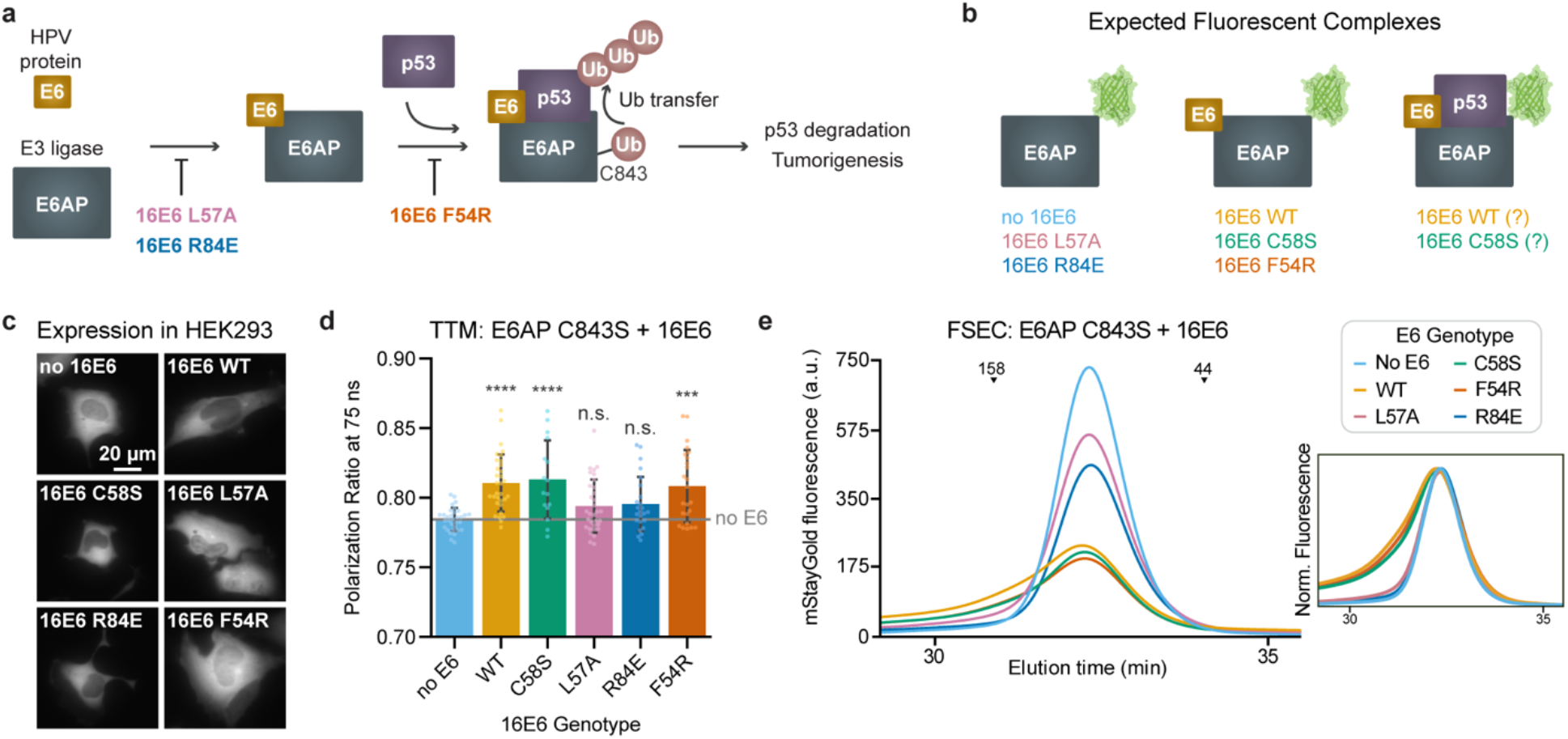
TTM detects small mass changes associated with 16E6:E6AP interactions in cells. (**a**) Schematic of the HPV 16E6:E6AP interaction and its role in p53 degradation and oncogenesis. 16E6 binding to E6AP promotes ubiquitination of p53 by E6AP. As a HECT E3-ligase, E6AP first loads ubiquitin onto its active cysteine (C843, isoform 2 numbering) and then transfers it to the p53 substrate.^42^ (**b**) Expected fluorescent complexes upon co-expression of 16E6 and mStayGold-tagged E6AP for 16E6 variants with different binding affinities. (**c**) Representative widefield fluorescence images of cells expressing E6AP C843S (catalytically dead) and 16E6 variants. 16E6 and E6AP were each under the control of a distinct promoter and polyadenylation sequence. (**d**) Polarization ratios from a cytosolic ROI (∼8 µm 1/e^2^ diameter) in HEK293 cells expressing C843S E6AP-mStayGold with the indicated 16E6 variant. To improve the signal-to-noise ratio, the full photon budget was concentrated into a single 75 ns delay with 10 replicate measurements. Bars represent the mean ± s.d. (n=16-32 cells). Asterisks indicate results of Dunnett post-hoc tests relative to the no 16E6 condition, following one-way ANOVA (F(5,154)=8.6, p<0.0001). (**e**) FSEC traces of HEK293 lysates expressing E6AP C843S-mStayGold and the indicated 16E6 variant. Inset shows a normalized view of the main peak profiles. Black arrows indicate elution time of molecular weight standards (kDa). a.u., arbitrary units.

To maximize sensitivity, polarization assays are typically designed such that the smaller binding partner (16E6, ∼15 kDa) is fluorescent, and the larger partner (E6AP, 100 kDa) is unlabeled.

However, because 16E6 is known to interact with numerous cellular proteins;^36^ we reasoned that tagging E6AP could provide a more specific readout. Therefore, we expressed mStayGold-tagged E6AP with V5-tagged 16E6 protein in HEK293 cells (**Fig. 5b**), with the goal of resolving a modest (10%) increase in E6AP molecular weight upon 16E6 binding.

Because the 16E6:E6AP interaction promotes complex turnover via self-ubiquitination,^37^ we included constructs with a catalytically dead E6AP mutant (C843S), which we expected to have more consistent protein levels. Indeed, WB analysis revealed reduced wild type E6AP levels when co-expressed with binding-competent 16E6 (**Supplementary Fig. 15**) and no such reduction in the context of E6AP C843S. Fluorescence intensity in cells tracked with the WB band intensities (**Fig. 5c, Supplementary Fig. 15, Supplementary Fig. 16**).

With both wild type and catalytically dead E6AP, co-expression with wild type 16E6 in HEK293 cells increased the polarization ratio at 75 ns relative to E6AP alone (**Fig. 5d, Supplementary Fig. 16**), consistent with 16E6 binding to E6AP. This signal depended on the affinity of 16E6 for E6AP, as tumbling of E6AP C843S co-expressed with the low binding affinity 16E6 L57A and R84E mutants^34,38^ was statistically indistinguishable from the condition without 16E6. The conservative mutation 16E6 C58S^39^ had a similar polarization ratio to wild type 16E6, as did the 16E6 F54R mutant designed to disrupt the p53 binding interface.^40^ The F54R result suggests that the polarization ratio change originates from the 16E6:E6AP interaction and not from a larger ternary complex. Wild type E6AP showed a similar trend, with additional cell-to-cell variability (**Supplementary Fig. 16**). In conditions with higher expression levels (in HEK293T cells, expression estimated from total fluorescence), we observe some 16E6:E6AP binding even for the lower affinity mutants (**Supplementary Fig. 16**). In HEK293 cells, we observed similar tumbling curves for E6AP isoforms 1 and 2, as well as with AZUL domain modifications reported to affect proteasome interactions^41^ (**Supplementary Fig. 17**). To orthogonally validate the TTM results, we used fluorescence-detected size exclusion chromatography (FSEC) to measure 16E6:E6AP interactions. We observed small shifts in elution time (**Fig. 5e, Supplementary Fig. 16**) when binding-competent 16E6 was co-expressed with E6AP, consistent with the observed shifts in polarization ratio.

Unexpectedly, catalytically dead E6AP had a lower polarization ratio at 75 ns than wild-type E6AP, suggesting a smaller fluorescent species. The difference was ubiquitination-dependent, as inhibition of the ubiquitin-activating enzyme UBA1 with TAK-243 decreased the polarization ratio in E6AP WT to match the catalytically dead constructs (**Supplementary Fig. 16**). FSEC (**Fig. 5e, Supplementary Fig. 16**) and reducing WB analyses (**Supplementary Fig. 15**) showed very similar molecular weight profiles for wild type E6AP and C843S, suggesting lysine-linked ubiquitination of the complex is not responsible for the polarization ratio change. To investigate whether differences in binding partners might explain the shift, we compared co-immunoprecipitation profiles of wild-type and C843S mutant E6AP (**Supplementary Fig. 18**), but we did not observe clear differences. However, nonreducing WB analysis revealed an approximately 10 kDa increase in molecular weight for wild type E6AP relative to C843S (**Supplementary Fig. 19**), consistent with ubiquitin loading on the active cysteine.

## Discussion

In this study, we demonstrated that triplet tumbling microscopy (TTM) provides rich information about protein interactions in living cells, expanding the size range of rotational diffusion assays to span the full proteome. Using a rapamycin-inducible FRB:FKBP dimerization system, we first showed that TTM can discriminate between protein complexes of different sizes on the timescale of seconds. Next, we investigated the homo-oligomerization of p53, observing a range of tumbling phenotypes that correlated with orthogonal biochemical measures of protein tetramerization. Finally, pushing the resolution limits of TTM, we detected the small (10% of molecular weight) increase associated with the binding of viral HPV 16E6 protein to the E3-ligase E6AP. Because TTM only requires a single tag to report diverse PPIs, it also enabled an unexpected observation during these experiments: wild type and catalytically dead E6AP tumble differently, likely due to ubiquitin loading on WT but not catalytically dead E6AP. Taken together, our data show that TTM can capture a variety of protein-protein interactions, including unexpected phenotypes that merit deeper follow-up investigations.

Although any sufficiently long-lived state could be used for tumbling measurements, triplets offer key advantages. Unlike reversibly switchable fluorescent proteins,^7^ infrared triggering of fluorescent protein triplets supports both short (100 ns) and long (microsecond) tumbling measurements, making this approach compatible with both small and large protein complexes. Relative to spontaneous processes (e.g. fluorescence, phosphorescence), triggerable triplet states provide “photons on demand,” concentrating the signal within the most informative time window. By selecting delays (**Fig. 1d**) with large polarization ratio differences for the particle sizes of interest and by accumulating thousands of pulse pairs within one camera exposure, we improved TTM’s signal-to-noise ratio relative to previous work with triplets,^14^ thus enabling single-trial detection in living cells.

Another advantage of TTM is its use of camera-based detection, which we did not fully realize here. In principle, TTM can record the whole field of view in one shot, dramatically reducing image acquisition time over point-based measures of translational diffusion like fluorescence correlation spectroscopy (FCS). In our prototype, available infrared power limited the accessible imaging area, but more powerful lasers spread over a larger area could image an entire cell simultaneously. Furthermore, the intensified camera we used here is a major source of photon loss, as its S20 photocathode has only 10% quantum efficiency at GFP emission wavelengths.

Although photocathodes with higher quantum efficiencies are commercially available, our early testing showed they did not adequately gate out prompt fluorescence from triggered triplets at short delays. Incorporation of a nanosecond-scale gating system with better detection efficiency, such as a SPAD camera,^43^ may improve these shortcomings. Finally, triplet tumbling measurements are not confined to microscopy and could be implemented in other instrument architectures. For example, a flow cytometer could be used for improved throughput,^15^ especially for pooled genetic screening applications.

Triplet tumbling measurements impose two unique demands on fluorescent labels: maximal triggered triplet signal and rigid coupling between the fluorophore and the protein of interest. Typical triplet yields from existing fluorescent proteins are less than 1%, limiting signal. Here, we worked around this limitation by using cellular overexpression systems and accumulating many excitation cycles per image. For protein-protein interaction measurements, it would be preferable to work at endogenous expression levels, which would be signal-limited for low abundance proteins. Looking ahead, because small molecule fluorophores with high triplet yields exist,^44^ hybrid chemical-genetic approaches or directed evolution campaigns could produce tags with more infrared triggered signal. A second requirement is that tumbling of the tag is rigidly coupled to the tumbling of the overall complex. Here, we increased tag rigidity by shortening the flexible “tails” of green fluorescent proteins, as previously reported for F-actin polarization labels.^22^ Even with a rigid fluorophore attachment, a long, flexible terminus on the protein of interest could decouple tag motion from the total molecular weight of the complex. Alternative labeling strategies such as internal insertion or elongated rigid linkers^7^ could potentially improve tag coupling. With any tagging, it is important to validate that protein function is largely unperturbed. In our study, we confirmed that the Kd of MBP-FRB and StayGold-FKBP matched reported values for untagged FRB and FKBP (**Supplementary Fig. 4**). Furthermore, when we tested published point mutants in p53 and E6, we observed the expected trends and localization, suggesting that the interactions were minimally perturbed.

For a rigid sphere in a homogenous solution of known viscosity and temperature, tumbling time is a direct readout of particle size. Quantitative interpretation for cellular proteins is more complex: proteins are nonspherical and flexible, cells contain heterogeneous mixtures of different PPI states, and the intracellular milieu is an active, crowded fluid of varying viscosity. Despite this, we observed approximately linear relationships between mass and tumbling time in both the rapamycin hetero-dimerization and the p53 homo-oligomerization systems, demonstrating that TTM can quantitatively report on complex size. For the recombinant rapamycin-induced heterodimerization system, we also estimated Kd, obtaining good agreement with steady-state SPR analysis (**Fig. 2f**, **Supplementary Fig. 4**). Estimation of Kd in cells requires knowledge of the intracellular protein concentrations, which is possible in principle but challenging in practice.

We observed different linear relationships between size and tumbling time for differently tagged proteins, highlighting the need for characterization and validation of each new tumbling tag. Characterization of recombinant tagged proteins by TTM before moving into cells, as we did with the rapamycin-inducible heterodimerization system, provides a solid foundation for connecting tumbling times to defined molecular species. However, purification and characterization of recombinant proteins is not possible in all circumstances. Without in vitro measurements, a combination of positive controls and orthogonal validation experiments can aid interpretation of tumbling curves. In the p53 studies, we used the well-characterized monomerizing mutant L344P, alongside FSEC and crosslinking, to benchmark the TTM data (**Fig. 4h**). In the HPV16E6 and E6AP studies, we used previously characterized 16E6 mutants, together with FSEC data, to connect tumbling times to the underlying molecular species. Even if no orthogonal validation is performed, tumbling times are still internally comparable, i.e. whether a protein’s effective size is different between untreated and drug treated conditions.

In this work, we analyzed mixtures in two contexts: rapamycin-driven hetero-dimerization and p53 homo-oligomerization. In the rapamycin dimerization system, we calculated the bound fraction via characterization of the recombinant fully bound and fully unbound states. In contrast, p53 exists as a mixture of monomers, dimers, and tetramers,^26,27^ making it difficult to isolate pure reference populations for calibration measurements. Furthermore, with the number of delay times collected, the p53 tumbling data were best approximated by a single exponential rotational diffusion term (**Supplementary Note 1**). We therefore opted to describe these data with a single tumbling time, which empirically tracks changes in the distribution of monomers, dimers, and tetramers. This analysis highlights a limitation of TTM: precise reporting of bound fractions in mixtures depends on robust fitting of amplitudes of a sum of exponential decays. Therefore, small changes (either small changes in mass or small changes in bound fraction) may be difficult to quantify; future improvements to the TTM signal-to-noise ratio as described above would improve this detection limit.

Because TTM only requires one interaction partner to be fluorescently tagged, it can reveal interactions without a priori knowledge of a binding partner. For example, during our analysis of the E6AP:16E6 system, we observed that wild type E6AP tumbled more slowly than catalytically dead E6AP. We initially considered a ubiquitin-dependent binding partner, but we did not observe a clear candidate via FSEC or immunoprecipitation (IP) (**Fig. 5e**, **Supplementary Fig. 16, Supplementary Fig. 18**). Instead, we speculate that the polarization ratio change may arise from ubiquitin loading onto the E6AP active site, consistent with the mechanism of HECT E3-ligases such as E6AP.^42^ The small mass increase visible on our non-reducing WB (**Supplementary Fig. 19**) is consistent with this interpretation. The active site of E6AP is near the C-terminus and our tag is C-terminal, so local steric effects may also change the polarization ratio. However, we cannot rule out an interaction that does not survive the cell lysis process required for FSEC or IP. For future efforts to identify interacting partners, we envision a pairing of TTM with modern proximity labeling technologies^45^ to identify candidate interactors in situ.

A key advance of our study is enabling the detection of strong intracellular PPIs from only one tagged binding partner. As such, TTM is not limited to PPIs; this approach could be expanded to other macromolecular interactions, e.g. those involving proteins bound to nucleic acids or carbohydrates. By circumventing the lifetime-limited size constraints of traditional fluorescence anisotropy, TTM provides a robust approach to quantify the assembly and dynamics of the proteome within the complex environment of the living cell.

## Data and Code Availability

Raw data, python processing scripts, and instrument control code used for each acquisition is available on Zenodo here: https://doi.org/10.5281/zenodo.20041587. Code to run the instrument is also available on GitHub at: https://github.com/calico/triplet_tumbling_microscopy.

## Acknowledgements

We thank Ian Foe and Autoosa Salari for helpful conversations about E6AP. We thank Clint Remarcik and Benjamin Tucker for advice on improving expression of E6AP constructs. We thank Thomas Bradbury, Jenny Clark, James Manton, Alice Walker, Andrea Volpato, and Ilaria Testa for helpful conversations about triplet photophysics. We thank Benjamin Adler for feedback on the manuscript. We thank Calico Life Sciences LLC for supporting this work.

## Author Contributions

M.I., A.G.Y., and J.R.L.D. conceptualized the project. J.R.L.D., A.M.S., and A.G.Y. designed the optical system. J.R.L.D. built the triplet tumbling instrument, performed the triplet tumbling microscopy experiments, analyzed the data, and wrote the original manuscript draft. J.R.L.D. and M.I. designed expression constructs and purified protein. Z.D.J., P.R., and S.K. performed biochemical cross-validation. J.R.L.D., M.I., M.P.L., H.B., Z.D.J., and A.H.N. contributed to the design and development of biological assays. A.M.S., A.H.N., and M.P.L. provided critical revisions and edited the manuscript. M.P.L. and A.G.Y. supervised the research. All authors reviewed and approved the final version of the manuscript.

## References

1. Phizicky, E. M. & Fields, S. Protein-protein interactions: methods for detection and analysis. Microbiol. Rev. 59, 94–123 (1995).

2. Piston, D. W. & Kremers, G.-J. Fluorescent protein FRET: the good, the bad and the ugly. Trends Biochem. Sci. 32, 407–414 (2007).

3. Magde, D., Elson, E. & Webb, W. W. Thermodynamic Fluctuations in a Reacting System— Measurement by Fluorescence Correlation Spectroscopy. Phys. Rev. Lett. 29, 705–708 (1972).

4. Ghosh, R. N. & Webb, W. W. Automated detection and tracking of individual and clustered cell surface low density lipoprotein receptor molecules. Biophys. J. 66, 1301–1318 (1994).

5. Einstein, A. Über die von der molekularkinetischen Theorie der Wärme geforderte Bewegung von in ruhenden Flüssigkeiten suspendierten Teilchen. Ann. Phys. 322, 549–560 (1905).

6. Weber, G. Polarization of the fluorescence of macromolecules. 1. Theory and experimental method. Biochem. J. 51, 145–155 (1952).

7. Volpato, A. et al. Extending fluorescence anisotropy to large complexes using reversibly switchable proteins. Nat. Biotechnol. 41, 552–559 (2023).

8. Austin, R. H., Chan, S. S. & Jovin, T. M. Rotational diffusion of cell surface components by time-resolved phosphorescence anisotropy. Proc National Acad Sci 76, 5650–5654 (1979).

9. Hogan, M., Wang, J., Austin, R. H., Monitto, C. L. & Hershkowitz, S. Molecular motion of DNA as measured by triplet anisotropy decay. Proc Natl Acad Sci U S A 79, 3518–22 (1982).

10. Moore, C., Boxer, D. & Garland, P. Phosphorescence depolarization and the measurement of rotational motion of proteins in membranes. FEBS Lett. 108, 161–166 (1979).

11. Peng, B. et al. Optically Modulated and Optically Activated Delayed Fluorescent Proteins through Dark State Engineering. J Phys Chem B 125, 5200–5209 (2021).

12. Byrdin, M., Duan, C., Bourgeois, D. & Brettel, K. A Long-Lived Triplet State Is the Entrance Gateway to Oxidative Photochemistry in Green Fluorescent Proteins. J. Am. Chem. Soc. 140, 2897–2905 (2018).

13. Ludvikova, L. et al. Near-infrared co-illumination of fluorescent proteins reduces photobleaching and phototoxicity. Nat. Biotechnol. 42, 872–876 (2024).

14. Lu, Y.-H. et al. Sequential Two-Photon Delayed Fluorescence Anisotropy for Macromolecular Size Determination. J. Phys. Chem. B 127, 3861–3869 (2023).

15. Lazzari-Dean, J. R., et al. From cameras to confocal to cytometry: measuring tumbling rates is a general way to reveal protein binding. Zenodo 10.5281/zenodo.10028432 (2023).

16. York, A. G., Ingaramo, M. del M. & Lazzari-Dean, J. Systems for triggering emission from triplet states and methods of use thereof. US Patent Number US20260194463A1. Filed 2023-11-10. Published 2026-07-09.

17. Landau, L. D. & Lifshitz, E. M. Fluid Mechanics. (1987).

18. Ando, R. et al. StayGold variants for molecular fusion and membrane-targeting applications. Nat. Methods 21, 648–656 (2024).

19. Zhang, H. et al. Bright and stable monomeric green fluorescent protein derived from StayGold. Nat. Methods 21, 657–665 (2024).

20. Banaszynski, L. A., Liu, C. W. & Wandless, T. J. Characterization of the FKBP·Rapamycin·FRB Ternary Complex. J. Am. Chem. Soc. 127, 4715–4721 (2005).

21. Santilli, R. T. et al. The Penn State Protein Ladder system for inexpensive protein molecular weight markers. Sci. Rep. 11, 16703 (2021).

22. Martins, C. S. et al. Genetically encoded reporters of actin filament organization in living cells and tissues. Cell 188, 2540–2559.e27 (2025).

23. Williams, J. W. & Morrison, J. F. The kinetics of reversible tight-binding inhibition. Methods Enzymol 63, 437–67 (1979).

24. Swaminathan, R., Hoang, C. P. & Verkman, A. S. Photobleaching recovery and anisotropy decay of green fluorescent protein GFP-S65T in solution and cells: cytoplasmic viscosity probed by green fluorescent protein translational and rotational diffusion. Biophys. J. 72, 1900–1907 (1997).

25. Volk, A. & Kähler, C. J. Density model for aqueous glycerol solutions. Exp Fluids 59, 75 (2018).

26. Gencel-Augusto, J. & Lozano, G. p53 tetramerization: at the center of the dominant-negative effect of mutant p53. Genes Dev. 34, 1128–1146 (2020).

27. Gaglia, G., Guan, Y., Shah, J. V. & Lahav, G. Activation and control of p53 tetramerization in individual living cells. Proc. Natl. Acad. Sci. 110, 15497–15501 (2013).

28. Laptenko, O., Tong, D. R., Manfredi, J. & Prives, C. The Tail That Wags the Dog: How the Disordered C-Terminal Domain Controls the Transcriptional Activities of the p53 Tumor-Suppressor Protein. Trends Biochem. Sci. 41, 1022–1034 (2016).

29. Stewart-Ornstein, J. & Lahav, G. p53 dynamics in response to DNA damage vary across cell lines and are shaped by efficiency of DNA repair and activity of the kinase ATM. Sci. Signal. 10, (2017).

30. Sun, X.-F. et al. Prognostic significance of cytoplasmic p53 oncoprotein in colorectal adenocarcinoma. The Lancet 340, 1369–1373 (1992).

31. Hurvich, C. M. & Tsai, C.-L. Regression and time series model selection in small samples. Biometrika 76, 297–307 (1989).

32. Kamada, R., Nomura, T., Anderson, C. W. & Sakaguchi, K. Cancer-associated p53 Tetramerization Domain Mutants. Quantitative analysis reveals a low threshold for tumor suppressor inactivation. J. Biol. Chem. 286, 252–258 (2011).

33. Kawate, T. & Gouaux, E. Fluorescence-Detection Size-Exclusion Chromatography for Precrystallization Screening of Integral Membrane Proteins. Structure 14, 673–681 (2006).

34. Wang, J. C. K. et al. Structure of the p53 degradation complex from HPV16. Nat. Commun. 15, 1842 (2024).

35. Münger, K. et al. Mechanisms of Human Papillomavirus-Induced Oncogenesis. J. Virol. 78, 11451–11460 (2004).

36. White, E. A. et al. Comprehensive Analysis of Host Cellular Interactions with Human Papillomavirus E6 Proteins Identifies New E6 Binding Partners and Reflects Viral Diversity. J. Virol. 86, 13174–13186 (2012).

37. Kao, W. H., Beaudenon, S. L., Talis, A. L., Huibregtse, J. M. & Howley, P. M. Human Papillomavirus Type 16 E6 Induces Self-Ubiquitination of the E6AP Ubiquitin-Protein Ligase. J. Virol. 74, 6408–6417 (2000).

38. Zanier, K. et al. Structural Basis for Hijacking of Cellular LxxLL Motifs by Papillomavirus E6 Oncoproteins. Science 339, 694–698 (2013).

39. Ye, X. et al. Discovery of reactive peptide inhibitors of human papillomavirus oncoprotein E6. Chem. Sci. 14, 12484–12497 (2023).

40. Zanier, K. et al. Solution Structure Analysis of the HPV16 E6 Oncoprotein Reveals a Self-Association Mechanism Required for E6-Mediated Degradation of p53. Structure 20, 604–617 (2012).

41. Kühnle, S. et al. Angelman syndrome–associated point mutations in the Zn2+-binding N-terminal (AZUL) domain of UBE3A ubiquitin ligase inhibit binding to the proteasome. J. Biol. Chem. 293, 18387–18399 (2018).

42. Scheffner, M., Nuber, U. & Huibregtse, J. M. Protein ubiquitination involving an E1–E2–E3 enzyme ubiquitin thioester cascade. Nature 373, 81–83 (1995).

43. Morimoto, K. et al. Megapixel time-gated SPAD image sensor for 2D and 3D imaging applications. Optica 7, 346–354 (2020).

44. Demissie, A. A. & Dickson, R. M. Triplet Shelving in Fluorescein and Its Derivatives Provides Delayed, Background-Free Fluorescence Detection. J Phys Chem 124, 1437–1443 (2020).

45. Qin, W., Cho, K. F., Cavanagh, P. E. & Ting, A. Y. Deciphering molecular interactions by proximity labeling. Nat. Methods 18, 133–143 (2021).

