## Supplementary Material for "Triplet tumbling microscopy enables in situ quantification of protein complex assembly and dynamics"

Julia R. Lazzari-Dean*, Alfred Millett-Sikking, Prashant Rao, Zena D. Jensvold, Sidharth Kashyap, Hannah Baddock, Maria Ingaramo, Aaron H. Nile, Andrew G. York, Magdalena Preciado López
Calico Life Sciences LLC, South San Francisco, CA, 94080, USA

### Contents

Supplementary Note 1
Materials and Methods
Supplementary Figures 1-19
Supplementary Tables 1-5
Amino Acid Sequences for all Constructs Used
Supplementary References

**Supplementary Note 1: Analyzing triplet tumbling microscopy (TTM) data**

Conventional fluorescence anisotropy analysis assumes emission is cylindrically symmetrical around the axis of the linearly polarized excitation beam, which allows definition of the total time-dependent intensity I(t) as in equation 1:

$$I(t) = I_x(t) + I_y(t) + I_z(t) = I_{\parallel}(t) + 2I_{\perp}(t) \quad (1)$$

This cylindrical symmetry assumption underlies the traditional calculation of anisotropy r(t), which for spherical fluorophores in solution is defined as a sum of exponentials[1] (equation 2). Each fluorophore size corresponds to a rotational correlation time $\theta_j$ in the exponential decay, with weights $m_j$ proportional to abundance:

$$r(t) = \frac{I_{\parallel}(t) - I_{\perp}(t)}{I(t)} = \frac{I_{\parallel}(t) - I_{\perp}(t)}{I_{\parallel}(t) + 2I_{\perp}(t)} = r_0 \sum_j m_j e^{-t/\theta_j} \quad (2)$$

In our TTM experiments with crossed linear polarizations of the 488 nm and infrared lasers, this assumption fails. We do not have cylindrical symmetry, and I(t) is a function of particle size, as the fraction of triplet-state fluorophores that have rotated enough to be triggered by the infrared laser depends on the rotational correlation time. Additionally, there may be imperfect alignment between the 488 nm absorption dipole ($S_0$->$S_1$), the IR absorption dipole ($T_1$->$T_n$) and the emission dipole ($S_1$->$S_0$). Because of this complexity, it is no longer straightforward to define a general I(t) normalization factor from which to calculate a traditional anisotropy r(t).

We can avoid symmetry assumptions by directly analyzing the "polarization ratio" of intensities parallel to the 488 nm beam (defined here as "parallel") and parallel to the infrared beams (defined here as "perpendicular"). To develop some intuition for how the polarization ratio should behave over time, we can first consider the simple case of a population of randomly oriented GFPs excited with linearly polarized 488 nm light (no IR trigger yet). If they are perfectly immobilized and excitation and emission dipoles are parallel, there will be 3x greater parallel intensity than perpendicular.[1] If the orientations scramble completely during the fluorescence lifetime, the parallel and perpendicular intensities will be equal, giving a polarization ratio of 1.

Moving to the TTM case, at long timescales, a fully tumbled population of triplets should have 3x more intensity in the perpendicular IR channel when triggered, i.e. a polarization ratio of 1/3, since the emission will be preferentially aligned with the IR beam. This is a direct analogy to the 488-only case described above, just the inverse of the ratio. Our data never quite reaches 1/3; factors such as non-parallel excitation dipoles (see above) and saturation in the IR depletion beam contribute to this. At shorter times, when the triplets have not fully scrambled their orientations, there will be more intensity along the parallel (488 nm polarization) axis, giving a higher polarization ratio.

In the most general case, the fluorescence intensity in either channel is proportional to the total number of molecules (N(t)) available, multiplied by a time-dependent orientation factor E(t) with constants defining an amplitude b and offset a (equations 3 and 4).

$$I_{\parallel}(t) \propto N(t) * \left(a_{\parallel} + b_{\parallel}E(t)\right) \tag{3}$$

$$I_{\perp}(t) \propto N(t) * \left(a_{\perp} + b_{\perp}E(t)\right) \tag{4}$$

For a population of uniform spheres, E(t) is a single exponential in the rotational correlation time θ (equation 5). In our system, proteins are not uniform spheres and there is often flexibility in the linkage between the fluorescent protein and the target protein, so the time constant is not a pure rotational correlation time. Throughout the text, we choose to refer to any fitted exponential time constant as "tumbling time" rather than "rotational correlation time." We typically see a clear linear relationship between "tumbling time" and mass (size), so we think that tumbling time is often a good proxy for rotational correlation time.

$$E(t) = e^{-t/\theta} \tag{5}$$

The constants $a_{\parallel}$, $b_{\parallel}$, $a_{\perp}$, and $b_{\perp}$ depend on the relative angles of the absorption and emission dipoles, as well as any saturation of the excitation beams, mixing effects from high NA collection, and depolarization during the fluorescence lifetime (e.g. homoFRET or fast rotation). Since at least some of these factors are unknown or variable, we chose to take them as fit parameters for our data. To cancel out the dependence on total number of molecules, we calculate the ratio of $I_{\parallel}$ and $I_{\perp}$, given as:

$$\frac{I_{\parallel}(t)}{I_{\perp}(t)} = \frac{a_{\parallel} + b_{\parallel}E(t)}{a_{\perp} + b_{\perp}E(t)} \tag{6}$$

For convenience of fitting, we can rearrange and simplify the constants in equation 6 into fewer, more interpretable parameters. At long times, the exponential terms go to zero, so we define $\rho_{\infty} \equiv a_{\parallel}/a_{\perp}$. At t=0, the exponential terms are equal to 1, so we similarly define $\rho_0 \equiv (a_{\parallel} + b_{\parallel})/(a_{\perp} + b_{\perp})$. If we substitute these into the equation above, it is also convenient to define $\kappa \equiv b_{\perp}/a_{\perp}$, which is the fractional change in the $I_{\perp}$ channel that is caused by rotation.

Putting this together, we obtain equation 7 as a general expression for the polarization ratio ρ:

$$\rho(t) = \frac{I_{\parallel}(t)}{I_{\perp}(t)} = \frac{\rho_{\infty} + [\rho_0(1+\kappa) - \rho_{\infty}]E(t)}{1 + \kappa E(t)} \tag{7}$$

For samples that are not well described as a population of spheres with a single rotational correlation time, we can generalize this equation with different forms of E(t). For mixtures of two possible states, such as bound and unbound components in the rapamycin dimerization dataset (**Figure 2**), we describe the rotational component as a sum of two exponentials, with $\theta_1$ from the unbound population and $\theta_2$ from the bound population (equation 8). The bound fraction $f_b$ determines the relative amplitude of these exponentials.

$$E(t) = (1 - f_b)e^{-t/\theta_1} + f_b e^{-t/\theta_2} \tag{8}$$

For samples that do not have single discrete populations but instead a spread of sizes from the manufacturing process, such as the latex beads used in **Figure 1**, we instead fit the data to a log-normal distribution of tumbling times τ with a characteristic median θ and log width σ.

$$E(t) = \int_0^\infty e^{-t/\tau} \frac{1}{\tau\sigma\sqrt{2\pi}} exp\left[-\frac{(\ln\tau - \ln\theta)^2}{2\sigma^2}\right] d\tau \tag{9}$$

To implement these models with our data, we first calculated a background-subtracted and g-factor-corrected version of the intensity ratio (equation 10). A circular region of interest encapsulating the entirety of either the parallel or perpendicular spot was averaged to determine the raw intensities. The background measured on the camera sensor with no light was 96 counts, and the g factor ($\frac{I_{\parallel,488}}{I_{\parallel,IR}}$) was 1.16, measured from a 10 μM solution of fluorescein in 0.1 N NaOH.

$$\rho = \frac{I_{\parallel,488} - background}{\left(I_{\parallel,IR} - background\right) * g} \tag{10}$$

We then fit the time-dependent changes in ρ to equation 7 using a nonlinear least squares algorithm (scipy least_squares).[2] The form of the time-dependent rotation term for each dataset is tabulated below, alongside a list of which parameters were fit globally for the dataset and which parameters were allowed to float for each time trace. To justify sharing parameters across all curves or constraining them to a global value for the dataset, we calculated the corrected Akaike information criteria (AICc)[3] and the Bayesian information criteria (BIC)[4] and chose the more complex model if ΔAICc was greater than 2. Because κ should depend on the fluorophore and any instrument saturation, but not on mass, we first determined it from a global fit on the purified protein data (**Figure 2**) and used that fixed value for all subsequent analysis. We found similar κ values for the mStayGold data (κ = -0.50 ± 0.08) and the mVenus data (κ = -0.44 ± 0.11), so we elected to fix κ to -0.5 for all analyses.

For the rapamycin bound fraction measurements, we fitted $\theta_1$ and $\theta_2$ as single exponentials to the limiting case recordings (fully unbound or fully bound) and then used those fixed values in the model to determine bound fraction in all traces.

In the p53 samples, we expected a mixture of particle sizes, which we expect to produce a multiexponential decay with three or more exponential terms. When the traces were averaged across multiple recordings for each mutant, both the AICc and BIC strongly favored the double exponential over a single exponential model (Δ > 10 in all conditions except for L344P, which had ΔAICc=6.8). However, for individual cell p53 recordings, the AICc favored the single exponential in most recordings (median ΔAICc = −10.8), whereas the BIC was near zero and slightly favored the single exponential (median ΔBIC = -0.7). Together, this indicates a second component is present but cannot be reliably resolved from a single cell recording. Because we wanted a fitting protocol that would be robust on the single cell level, we opted to process all p53 data with single exponential fitting.

| Dataset | Rotational Model | Globally fit parameters | Parameters fit per trace |
|---|---|---|---|
| mStayGold on beads | Distribution (eqn. 9) | | $\theta$, $\sigma$, $\rho_0$, $\rho_\infty$ |
| Rapamycin size ladder | Single exponential (eqn. 5) | $\rho_0$, $\rho_\infty$ | $\theta$ |
| Rapamycin bound fraction | Sum of 2 exponentials (eqn. 8) | $\theta_1$, $\theta_2$, $\rho_0$, $\rho_\infty$ | $f_b$ |
| p53 oligomerization | Single exponential (eqn. 5) | | $\theta$, $\rho_0$, $\rho_\infty$ |

To more directly justify our choice of ratio-based fits with our data, we compared our ratio-based exponential model to exponential fits on either anisotropy (equation 2, which incorrectly assumes cylindrical symmetry) or polarization (equation 11, as calculated for the related OADFA technique[5]).

$$P(t) = \frac{I_{\parallel}(t) - I_{\perp}(t)}{I_{\parallel}(t) + I_{\perp}(t)} \tag{11}$$

The use of polarization was justified in the original reference using a derivation that assumed emission was restricted to the planes of the laser triggering, i.e. $I_{tot}=I_x+I_y$, despite the 3D nature of diffusion.

Although neither polarization nor anisotropy is physically justified for our data, we wanted to benchmark their performance against the ratio model. We chose to compare fits using the in vitro rapamycin dataset (**Fig. 2B**). As with the ratio calculations, we designated the polarization of the 488 nm beam as parallel and the polarization of the infrared beam as perpendicular, which yields negative P and r values at long timescales. The polarization and anisotropy values were fit to a single exponential decay (equation 12), and the single exponential form of E(t) (equation 5) was used for the ratio fits.

$$P(t) = ae^{-t/\theta} + c \tag{12}$$

All three fits recovered a clear dependence of θ on mass, with modest shifts in the calculated θ values (see Scheme 1 below). The ratio model performed the best by ΔAICc and ΔBIC, although fitting polarization performed similarly. Because the ratio model is the most general and doesn't require assumptions about symmetries in the angular distributions, we represent our data here in terms of ratios.

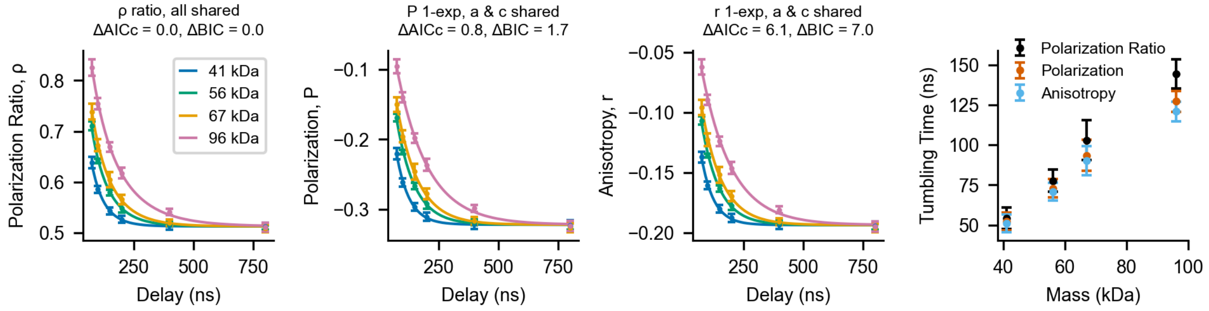


**Scheme 1: Comparison of ratio, polarization, and anisotropy-based fits.**

Comparison of ratio, polarization, and anisotropy-based fits for the rapamycin size ladder data in Figure 2. All error bars indicate mean ± s.d., and solid lines indicate fit curves. ΔAICc and ΔBIC are calculated relative to the ratio fits (leftmost panel).

### Materials and Methods

*Materials*

Suppliers and catalog numbers for reagents are available in **Supplementary Table 3**.

*Tumbling instrument design*

The essential components of a TTM measurement are control over excitation and emission polarization, nanosecond-scale control over illumination sequence, and a fast shutter to separate the prompt fluorescence from the triggered triplet signal. We chose to implement these components on a custom-built widefield microscope, but this concept could be applied in a variety of optical configurations, as we previously simulated.[6] **Supplementary Fig. 1** is a detailed diagram of the system, and **Supplementary Table 4** enumerates all optical components.

To control illumination with nanosecond precision, we used the fast digital modulation of the diode lasers (Cobolt 06-01 MLD: 200 mW 488 nm, 250 mW 785 nm, 250 mW 830 nm, 250 mW 915 nm, 250 mW 940 nm; Hubner Photonics) to generate variable duration pulses (nanosecond to microsecond). We opted for four separate infrared lasers, combined with dichroic beamsplitters, to access higher total IR powers at reduced cost. For the data in this study, the four infrared lasers were used simultaneously. A digital delay generator (BNC 575, Berkeley Nucleonics Corporation) provided the digital modulation input to the lasers. The output of each laser was already linearly polarized; we used half-wave plates to set the 488 at crossed polarization with the infrared lines and to correct off small offsets in output polarization among the infrared lasers.

To produce an 8 μm diameter ($1/e^2$) illumination spot in the sample, we expanded the 488 nm laser output to a ~2 mm beam and relayed that to the back focal plane of the objective (Nikon 25x 1.1NA Apo water immersion LWD). We similarly expanded the infrared laser outputs, combined them with dichroic beamsplitters, and used another dichroic beamsplitter to relay them to the back focal plane of the objective. The combination of dichroic beamsplitters in the emission path transmit wavelengths from 488 nm to 750 nm. We then split this emission light with a polarizing beamsplitter, focused it with tube lenses, and steered the horizontal and vertically polarized outputs next to each other on the intensified camera (pco.dicam C1 with 18 mm S20 intensifier and P46 phosphor, Excelitas Technologies) with a knife edge mirror. We characterized spot diameter and verified overlap of the different colors with a camera at the sample and with a mixture of fluorescein and upconverting nanoparticles (808 nm nominal absorption, Millipore Sigma) on the widefield path (described below).

Because the triggered triplet signal is identical in emission spectrum and fluorescence lifetime to the prompt emission,[6] the primary way to separate the two signals is in time. We achieved this fast shuttering with an intensified camera and gating signals from the digital delay generator. The image intensifier we used had an S20 photocathode, which has a 10% quantum efficiency (QE) at 500 nm but a shutter ratio 100x higher than the GenIII GaAsP photocathode (50% QE at 500 nm). We saw some bleedthrough of the singlet signal with the GenIII intensifier (data not shown) and therefore opted to use the S20 intensifier despite its lower QE. To protect the intensified camera from accidental light exposure, we fitted it with a shutter (SmartShutter, Sutter Instruments).

To facilitate sample finding, we incorporated a widefield bypass with an LED and an sCMOS camera (pco.panda 4.2 bi, Excelitas Technologies). An automated flipper mirror switched the visible (488 nm) path between the LED illumination and the laser illumination, and a second automated flipper mirror switched the emission path between the sCMOS and the intensified camera. An OKO UNO stagetop incubator (OkoLabs) provided temperature control at the sample. A custom XYZ stage assembly (Physik Instrumente) allowed for positioning of the sample at the objective.

Communication between the PC and devices was done with python device adapters, which were coordinated by a central python module. A user interface interacting with the central python module allowed for real-time visualization of samples. An analog out card (National Instruments PCIe-6738) provided the voltage signals to allow microsecond synchronization between devices, in addition to slower serial port commands for settings changes. The full codebase is available on GitHub at https://github.com/calico/triplet_tumbling_microscopy.

*Settings for tumbling data acquisition*

The pulse sequence depicted in **Figure 1d** was used unless otherwise indicated. All infrared lasers were used simultaneously to maximize triggering efficiency. **Supplementary Table 5** lists nominal laser output and measured power at the sample for each wavelength, and **Supplementary Table 1** indicates additional relevant acquisition parameters. Power was measured with a slide power meter head (Thorlabs S170C on PM200 power meter). Losses in power at the sample for increasingly red infrared lines is likely attributable to the transmission profile of the objective lens.

*Molecular cloning*

All plasmids were generated by Gibson assembly using NEB HiFi assembly master mix (New England Biolabs) or were synthesized by an external vendor (VectorBuilder or Genscript). Final constructs were confirmed by whole plasmid sequencing (Plasmidsaurus); amino acid sequences are available in the supplementary material and DNA sequences are available with the source data on Zenodo (https://doi.org/10.5281/zenodo.20041587). Plasmid DNA for mammalian transfection was miniprepped from *E. coli* NEB 5-alpha or outsourced for midiprep production (Azenta). Constructs used for protein expression were propagated in *E. coli* NEB 5-alpha or *E. coli* BL21(DE3). All *E. coli* strains were obtained from New England Biolabs.

*Protein purification*

$His_6$-tagged proteins were expressed in *Escherichia coli* BL21(DE3) and purified as described below. Purity was assessed by SDS-PAGE and Coomassie stain (GelCode Blue, Thermo Fisher) (**Supplementary Fig. 4**). For constructs that did not yield sufficient purity, a secondary affinity purification step was performed using a Strep tag located at the opposite terminus.

To express recombinant proteins, chemically competent BL21(DE3) were transformed with pRSET B vectors and plated onto LB agar plates supplemented with 100 μg/mL carbenicillin. Plates were incubated overnight at 35°C and then maintained at room temperature for an additional 24-36 hours. *E. coli* were scraped from the surface of the plate and resuspended in lysis buffer (300 mM NaCl, 50 mM sodium phosphate, pH 8) supplemented with BugBuster Protein Extraction Reagent (non-denaturing detergent-based, Millipore Sigma). Cells were lysed at room temperature for 30 minutes with gentle rocking, followed by centrifugation at 18,000 x g for 20 minutes at 4°C. Supernatants were collected and transferred to a fresh tube containing 50-100 μL of washed Ni-NTA resin (NEBExpress, New England Biolabs). Lysates were incubated with resin for 1 hour at room temperature, followed by three washes in lysis buffer. Protein was eluted with lysis buffer containing 500 mM imidazole. If an additional purification step on Strep resin was performed, eluates were transferred to washed StrepTactin SuperFlow Plus resin (Qiagen) and allowed to bind for 1 hour at 4°C. Resin was washed three times in PBS, and proteins were eluted in PBS with 2.5 mM desthiobiotin. After the final elution, proteins were buffer exchanged to PBS and spin concentrated with 10 kDa MWCO PES spin concentrator (Pierce, Thermo Fisher Scientific). Proteins in PBS were stored at 4°C for short term use.

Purity was assessed by SDS-PAGE. Protein samples were mixed with 2x Tris-glycine SDS sample buffer containing 5% beta-mercaptoethanol and heated to 85°C for 5 minutes. Samples were resolved on a Tris-glycine 4-20% gradient gel (Thermo Fisher) run at 225 V for 40-50 minutes. Gels were fixed for 20-30 minutes in a solution of 50% methanol, 10% acetic acid, and 40% water, rinsed with deionized water three times, and stained overnight with colloidal Coomassie G-250 (GelCode Blue Stain, Thermo Fisher). Gels were destained in deionized water and imaged (ChemiDoc, Bio-Rad).

Protein concentration was assessed with UV-Vis absorbance (Nanodrop 2000C Spectrophotometer, Thermo Fisher). For fluorescent proteins, published absorption coefficients were used to determine concentration based on the peak fluorescence absorption (mVenus[7] 515 nm, 104,000 $M^{-1}$ $cm^{-1}$; mStayGold[8] 499 nm, 164,000 $M^{-1}$ $cm^{-1}$; mEGFP[9] 488 nm, 56,000 $M^{-1}$ $cm^{-1}$; mBaoJin[10] 500 nm, 128,000 $M^{-1}$ $cm^{-1}$; mNeonGreen[11] 506 nm, 116,000 $M^{-1}$ $cm^{-1}$; Clover[12] 505 nm, 111,000 $M^{-1}$ $cm^{-1}$). For nonfluorescent proteins, absorbance at 280 nm with an extinction coefficient predicted by the Expasy ProtParam tool was used.[13]

Some of the fluorescent protein samples used here were purified by contract research organizations (Biortus for mEGFP, Sino Biologicals for mStayGold); for those samples, purity was verified by SDS-PAGE.

*Cell culture and transfection*

U2OS, HEK293, or HEK293T cells were maintained in a 37°C humidified incubator at 5% $CO_2$ in Dulbecco's modified Eagle Medium (DMEM with 4.5 g/L glucose without phenol red) supplemented with 10% FBS, 2 mM GlutaMAX and 1 mM sodium pyruvate. Confluent cells were dissociated with TrypLE Express and passaged into fresh media every 2-4 days when ~90% confluent. Cultures were tested for mycoplasma contamination monthly. All media components were purchased from Gibco (Thermo Fisher Scientific)

For imaging experiments, 8-well chambered #1.5 coverglass (CellVis) was coated with bovine fibronectin (~1 mg/mL, Millipore Sigma, diluted 100x in PBS) for at least 1 hour at 37°C, followed by three washes with PBS. Cells were seeded onto dishes at 25,000 cells per well and allowed to adhere at room temperature for 20-40 minutes before being transferred to the incubator.

For transient transfection, cells were seeded approximately 24 hours before lipofection with TransIT-LT1 (Mirus Bio) complexes in Opti-MEM (Gibco) according to the manufacturer's instructions (250-650 ng DNA/well, 2 µL transfection reagent per µg DNA). Cells were imaged approximately 24 hours after transfection. A full media change into complete DMEM supplemented with 25 mM HEPES was performed before imaging to reduce pH drift, as the TTM instrument's sample incubator maintained temperature at 36°C but did not control $CO_2$. Cells were left on the microscope in that environment for a maximum of 2 hours.

*Sample preparation for in vitro tumbling measurements*

Triplet properties of fluorescent proteins (**Supplementary Fig. 3**) were characterized using purified proteins immobilized in polyacrylamide gels. Gel solution was prepared immediately before use from 30% acrylamide/bis-acrylamide (37.5:1, Sigma-Aldrich) mixed at a ratio of 100 µL acrylamide, 1.2 µL of 10% ammonium persulfate in water (made fresh) and 0.12 µL TEMED. 10 µL of 50 µM fluorescent protein in PBS was mixed with 90 µL of gel mix in a well of an 18 well chambered coverglass (CellVis) and allowed to set overnight before imaging.

Rapamycin-induced binding measurement on purified protein were performed at room temperature in PBS with 20% glycerol (v/v) to approximate cytoplasmic viscosity.[14] Samples were imaged in glass-bottom 384 well plates that had been blocked with 1 mM BSA in 10 mM phosphate for at least 1 hour. After blocking, plates were washed 3x in PBS before addition of proteins. The protein-based components were first mixed (1 µM fluorescent component and 0-2 µM nonfluorescent binding partner), and rapamycin was added to a final concentration of 2 µM in 50 µL total volume. Samples were incubated at room temperature for 5-40 minutes between rapamycin addition and imaging; tumbling curves were stable over this time range.

mStayGold-coated CML latex beads (Thermo Fisher) were imaged on BSA-blocked glass bottom 384-well plates. Samples were prepared by first mixing 10 µL of 100 mM MES pH 6.3 with 1 µL of 1% Tween in 100 mM MES pH 6.3 and x µL of 80 µM mStayGold, where x varied with bead surface area (4.44 µL, 2.22 µL and 1.45 µL for 20, 40, and 60 nm diameter beads, respectively). 10 µL of CML latex beads were then added and mixed thoroughly. Samples were incubated for at least 5 minutes before imaging and were not kept for longer than an hour to avoid aggregation. The ratios above were identified by successively decreasing fluorescent protein concentration until unbound protein was minimized, as previously described.[6]

*$K_d$ estimation for the FRB:rapamycin:FKBP interaction*

After fitting the tumbling curves to a double exponential model (equation 8, Supplementary Note 1), we obtained a fraction bound at each input concentration. We used this information to estimate apparent $K_d$ for our tagged FRB and FKBP components. Because our concentrations of

FRB and FKBP were similar, we compensated for ligand depletion in our calculation of $K_d$ as previously described[15] (equation 4), where $f_b$ is the experimentally determined fraction bound, $[FKBP]_t$ is the total concentration of mStayGold-FKBP (assuming 2 µM of rapamycin with 1 µM mStayGold-FKBP yields complete binding), $[FRB]_t$ is the total concentration of MBP-FRB, and $K_d$ is the dissociation constant.

$$f_b = \frac{[FRB]_t + [FKBP]_t + K_d - \sqrt{([FRB]_t + [FKBP]_t + K_d)^2 - 4[FRB]_t[FKBP]_t}}{2[FKBP]_t} \quad (13)$$

*Fluorescence anisotropy measurements (singlets)*

Polarization measurements from prompt fluorescence were performed on a Leica SP8 inverted point-scanning confocal equipped with a PicoHarp 300 time-correlated single photon counting module (PicoQuant). Cells were maintained at 36°C and 5% $CO_2$ throughout the experiment with a stagetop $CO_2$ chamber and a large temperature enclosure around the microscope (Okolabs). Pulsed excitation light (488 nm, 80 MHz, approximately 0.3 µW point focused at the sample) was generated by a white light laser and delivered to the sample via a 40x air objective (NA 0.85, HC PL APO CS). Emission passed through a confocal pinhole (1 AU) and was directed to the back port of the instrument, where a polarizing beamsplitter directed the light to two SPAD detectors collecting the parallel and perpendicular components of the light. Green fluorescence was selected with a 488 nm notch filter to block the laser and a 531/40 nm bandpass filter (Semrock) on the detectors. Images were generated by accumulating photons for 100 sweeps over a 58 µm x 58 µm field of view (approximately 55 seconds).

Anisotropy, r, was calculated using the standard formula (equation 2, **Supplementary Note 1**) from the parallel ($I_{\parallel}$) and perpendicular ($I_{\perp}$) emission. A g factor ($\frac{I_{\parallel}}{I_{\perp}}$) was measured from a 10 µM solution of fluorescein in 0.1 N NaOH and was applied to all data. Using a custom parsing script, we summed the photons over the entire image and binned 32x in time (approximately 100 bins in the final recording). The parsing script is available on GitHub (https://github.com/AndrewGYork/tools/blob/master/picoquant_tttr.py) and with the source data.

*Western blotting*

HEK293 or U2OS cells were plated into 6 well plastic tissue culture plates 48 hours before lysis at a density of 600,000-1,000,000 cells per well. Approximately 24 hours after plating, cells were transfected with as described above using Mirus LT1 transfection reagent at a total volume 12x larger than what was used for the same plasmids in 8 well chambers. Cells were returned to the incubator for approximately 24 hours before lysis.

For crosslinking experiments, prior to lysis, cells were washed 3x with PBS and crosslinked at room temperature for 10 minutes with PBS containing 2 mM disuccinimidyl glutarate (Thermo Fisher, made fresh as a 25 mM DMSO stock). Crosslinking solutions were removed and immediately replaced with Tris-containing lysis buffer to quench the reaction. Cells were placed on ice and then lysed as described below.

For analysis of total protein levels, cells were placed on ice, washed once with PBS, and lysed in 100-200 µL of either RIPA lysis buffer (crosslinking or total protein levels: 25 mM Tris•HCl pH 7.6, 150 mM NaCl, 1% NP-40, 1% sodium deoxycholate, 0.1% SDS) or Pierce IP lysis buffer (nonreducing blots: 25 mM Tris-HCl pH 7.4, 150 mM NaCl, 1 mM EDTA, 1% NP-40, 5% glycerol) with 1x Halt Protease Inhibitor Cocktail. After 5 minutes of lysis, remaining adherent materials were dislodged by gentle scraping with the back of a P1000 pipette tip. Lysates were transferred to 1.5 mL Eppendorf tubes and clarified by centrifugation at 14,000 x g at 4ºC for 10 minutes. Supernatants were transferred to fresh tubes and either used immediately or flash frozen at -80ºC until analysis.

Total protein content of the lysates was measured with a modified Lowry assay (DC assay, BioRad) and read out on a CLARIOstar plate reader (BMG Labtech). For SDS-PAGE, samples were prepared in Tris-glycine sample loading buffer at 1x final concentration containing 5% beta-mercaptoethanol. 5 to 10 µg of total protein per lane was used. For non-reducing gels, the beta-mercaptoethanol was omitted. Samples were heated at 85ºC for 5 minutes, briefly centrifuged, and resolved on a 4-12% gradient Tris-glycine gel (15-well, Thermo Fisher) run at either 225 V for 35-45 minutes or 125 V for 90 minutes.

Proteins were transferred to PVDF membranes (0.2 µm pore, premade stacks from BioRad) using the TransBlot Turbo Semi-Dry Transfer system (BioRad) with either the Mixed MW or the High MW preset protocols (for a single mini gel, 1.3 A for 7 or 10 minutes respectively). For all subsequent steps, incubations were performed with gentle rocking. Membranes were immediately placed into TBST (20 mM Tris HCl, 150 mM NaCl, 0.1% Tween-20, pH 7.5) and then blocked in 5% dry milk in TBST for 1-4 hours. Primary antibodies were incubated overnight at 4ºC at 1:1000 dilution. Antibodies were washed off with 5 x 5 minute washes in TBST, followed by secondary antibody incubation (1:1000 dilution in 5% dry milk in TBST, room temperature, 1 hour) and another set of 5 x 5 minute washes in TBST. Membranes were developed using SignalFire ECL Reagent (Cell Signaling Technologies), and chemiluminescent signal was imaged on a ChemiDoc imager (BioRad). GAPDH levels were used to normalize loading.

*Western blot analysis of p53 monomer, dimer, trimer, and tetramer distribution*

For quantification of p53 oligomerization after crosslinking, four resolved bands were measured and assigned as putative monomeric, dimeric, trimeric, and tetrameric species relative a non-crosslinked control. The trimer likely arises from incomplete crosslinking. The intensity of each band ($I_m$, $I_d$, $I_{tri}$, $I_{tet}$, $I_{tot}$ for monomer, dimer, trimer, tetramer, and total) was measured in ImageJ,[16] and the fraction that each band represents of the total was determined. The intensity-weight average apparent mass is the weighted average size across all bands (equation 14). The monomer mass (mm) used was 85 kDa (p53 + mStayGold tag).

$$Apparent\ mass = \frac{I_m}{I_{tot}} * mm + \frac{I_d}{I_{tot}} * mm * 2 + \frac{I_{tri}}{I_{tot}} * mm * 3 + \frac{I_{tet}}{I_{tot}} * mm * 4 \qquad (14)$$

*Co-immunoprecipitation*

Cell lysates were prepared and cleared as described for Western blotting above, using Pierce IP lysis buffer (25 mM Tris-HCl pH 7.4, 150 mM NaCl, 1 mM EDTA, 1% NP-40, 5% glycerol, 1x Halt Protease Inhibitor Cocktail) to better preserve protein-protein interactions. Cells were either lysed directly from culture or trypsinized, spun down, washed in PBS, flash frozen in liquid nitrogen as pellets, and stored at -80 ºC before lysis.

Cell lysates were quantified by Pierce 660 nm Protein Assay (Life Technologies). For each immunoprecipitation, 150 µg of total lysate was incubated with 50 µL of anti-FLAG M2 affinity gel slurry (Millipore) in 500 µL of Pierce lysis buffer for 2 hours at 4ºC with rotation. Agarose beads were washed twice with 1 mL wash buffer (25 mM Tris pH 7.5, 150 mM NaCl, 5% glycerol). Bound protein was eluted with 50 µL of 500 ng/µL 3XFLAG peptide in wash buffer, shaking at 1000 rpm for 30 minutes at 25 ºC. Eluate was separated from agarose beads using microspin columns. Input and eluted samples were resolved on 4-20% Criteron TGX gels and analyzed either by Western blotting as described above or by silver staining (Pierce silver stain kit).

*Fluorescence-detection Size Exclusion Chromatography (FSEC)*

Cell lysates were generated as described above with Pierce IP lysis buffer (25 mM Tris-HCl pH 7.4, 150 mM NaCl, 1 mM EDTA, 1% NP-40, 5% glycerol) with 1x Halt Protease Inhibitors. For most samples, 75 µL of cell lysate was injected onto a Superose 6 Increase 10/300 GL column (Cytiva) equilibrated in Pierce IP Lysis buffer at a flow rate of 0.5 mL/min. Lysates were kept in The column and running buffer were kept at room temperature; samples were maintained at 4ºC until injection. For lysates containing E6AP (WT) co-expressed with 16E6 WT, 16E6 C58S, or 16E6 F54R, 300 µL of lysate was injected due to the lower mStayGold fluorescence signal. Separations were performed on a Shimadzu HPLC system equipped with an RF-20Axs fluorescence detector, monitoring mStayGold fluorescence at 480 nm excitation and 512 nm emission, as described previously.[17] Where normalized traces are shown, elution traces were scaled to the highest peak intensity within the comparison group using the Shimadzu LabSolutions analytical software version 5.132.

A molecular weight standard curve (**Supplementary Fig. 12b**) was generated by injecting 30 µL of Gel Filtration Standard (BioRad, cat no. 1511901, diluted to 1:10 in water based on the manufacturer's recommendations). Standards were detected by tryptophan fluorescence (excitation: 280 nm, emission: 312 nm) using the same RF-20Axs fluorescence detector.

*Surface plasmon resonance (SPR)*

SPR was performed on a Biacore 8K Plus (Cytiva). Approximately 450 RU of MBP-FRB was immobilized onto an NTA chip (Catalog # BR100532) using His-NTA oriented lysine coupling of His-tagged MBP-FRB to carboxymethyl dextran. The chip was conditioned with 350 mM EDTA for 60 seconds, followed by a system wash with HBS-EP (10 mM HEPES pH 7.6, 150 mM NaCl, 3 mM EDTA, 0.05% Tween-20). The NTA surface was then activated with a 60 second injection of 500 µM $NiSO_4$. The system was washed with HBS-EP. The chip surface was then activated for 420 seconds with EDC/NHS (Cytiva Amine Coupling Kit). The flow system was then washed with ethanolamine to quench. His-tagged MBP-FRB (20 nM) was then injected for 150 seconds to yield 450 RU of surface density. The surface was then deactivated with a 420-second injection of ethanolamine followed by a 60-second injection of 350 mM EDTA to remove residual nickel from the surface. The flow system was washed with HBS-EP, and the surface was allowed to equilibrate in running buffer (10 mM HEPES pH 7.6, 150 mM NaCl, 3 mM EDTA, 0.05% Tween-20, 1% DMSO) for 15 minutes.

The binding kinetics of mStayGold-FKBP:rapamycin binary complexes to MBP-FRB were measured using single cycle kinetics. mStayGold-FKBP was pre-incubated with four concentrations of rapamycin (1, 3, 5, and 10 µM) and injected across the surface at six concentrations ranging from 3.7 to 900 nM. All injections were performed at a 30 µL/min flow rate. Analytes were injected for 200 seconds and allowed to dissociate for 20 minutes. Two replicates were measured. Sensograms were double referenced by subtracting the signal from the reference channel (no ligand) and the signal from the DMSO blank injection.

The data were analyzed using Biacore Insight Software (Cytiva, version 7.0.9.1992) with both a two-state binding model and a steady state affinity model. A 1:1 Langmuir model gave poor fits with systematic residuals ($\chi^2 \approx 200$–$260$ RU²) and could not describe the biphasic dissociation. A two-state (conformational change) model reduced $\chi^2$ to $\approx 7$–$12$ RU² and removed the structured residuals, so it was selected for global analysis. We also performed steady state analysis to directly compare to the endpoint TTM measurements of apparent $K_d$. The sensograms reached a steady state (flat top on each injection), enabling the use of a steady state model to mimic an endpoint measurement.

*General notes: statistical testing*

Throughout this work, asterisks are used to represent significance levels of statistical tests: ns p>0.05, * p<0.05, ** p<0.01, *** p<0.001, **** p<0.0001.

*Software packages and versions*

Image inspection and analysis was performed in ImageJ.[16] Tumbling analysis was done in python 3.12.9 with the use of the following packages:
-Numpy (version 2.2.3)[18]
-Scipy (version 1.15.2)[2]
-Pandas (version 2.2.3)[19]
-Matplotlib (version 3.10.1)[20]

*Data and Code Availability*

Raw data, python processing scripts, and instrument control code used for each acquisition is available on Zenodo here: https://doi.org/10.5281/zenodo.20041587. Code to run the instrument is also available on GitHub at: https://github.com/calico/triplet_tumbling_microscopy.

**Supplementary Figure 1. Detailed schematic of triplet tumbling instrument**

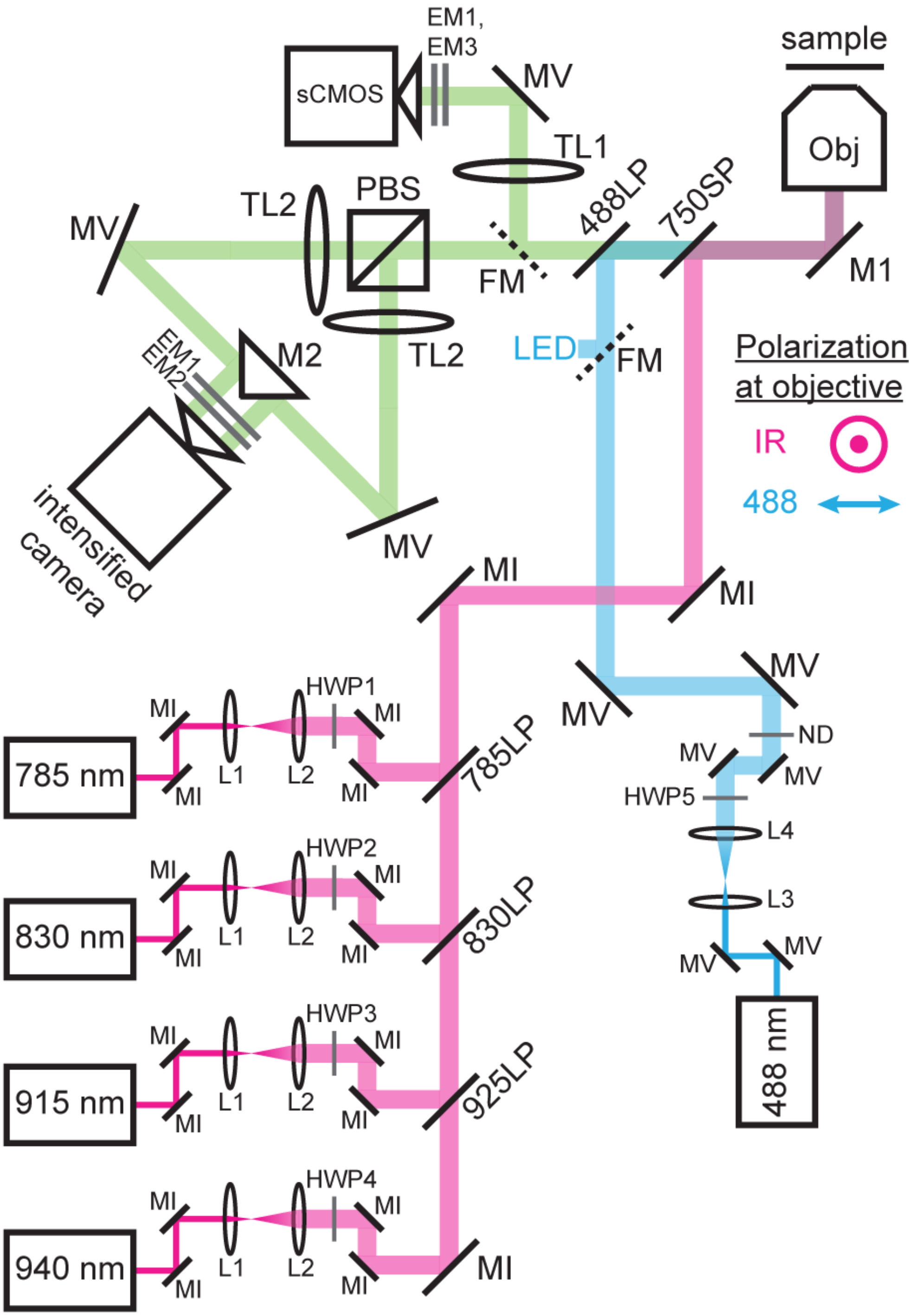


**Supplementary Figure 1. Detailed schematic of triplet tumbling instrument**
A full parts list is available in **Supplementary Table 4**. M: mirror, L: lens, HWP: half-wave plate, ND: neutral density filter (movable), TL: tube lens, Obj: objective lens, PBS: polarizing beamsplitter. The half-wave plates were used to align the output polarizations of the various lasers to a coordinate system defined by the polarizing beam splitter on the emission path.

**Supplementary Figure 2. Triggered triplet signal requires both 488 nm and infrared illumination**

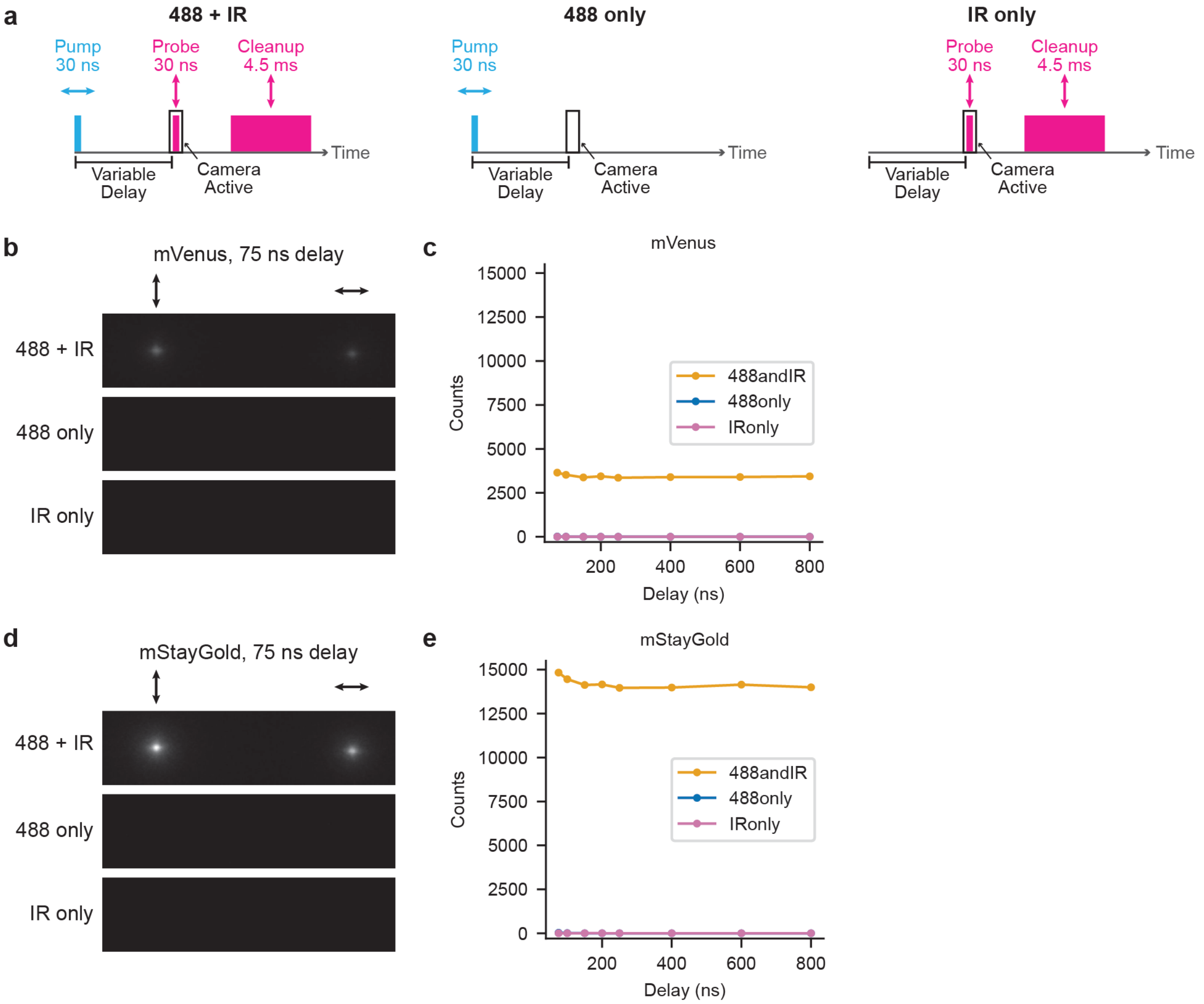


**Supplementary Figure 2. Triggered triplet signal requires both 488 nm and infrared illumination**

(**a**) Schematic of illumination sequences used. Each exposure consisted of 39920 cycles of the indicated pulse sequence (total of 1 second). Delays are measured from the start of the 488 nm pulse to the start of the infrared pulse. (**b**) Fluorescence emission images acquired at a 75 ns delay for 2 µM mVenus in PBS. Emission polarized perpendicular to the 488 beam forms the lefthand spot and emission polarized parallel to the 488 beam forms the righthand spot. (**c**) Summed counts (parallel and perpendicular) for a delay sequence measured on 2 µM mVenus in PBS. Each point indicates the average of two images taken on the same sample. (**d**) Images from a 75 ns delay for 2 µM mStayGold in PBS. (**e**) Summed counts (parallel and perpendicular) for a delay sequence measured on 2 µM mStayGold in PBS.

### Supplementary Figure 3. Comparing triggerable triplet properties across green fluorescent proteins

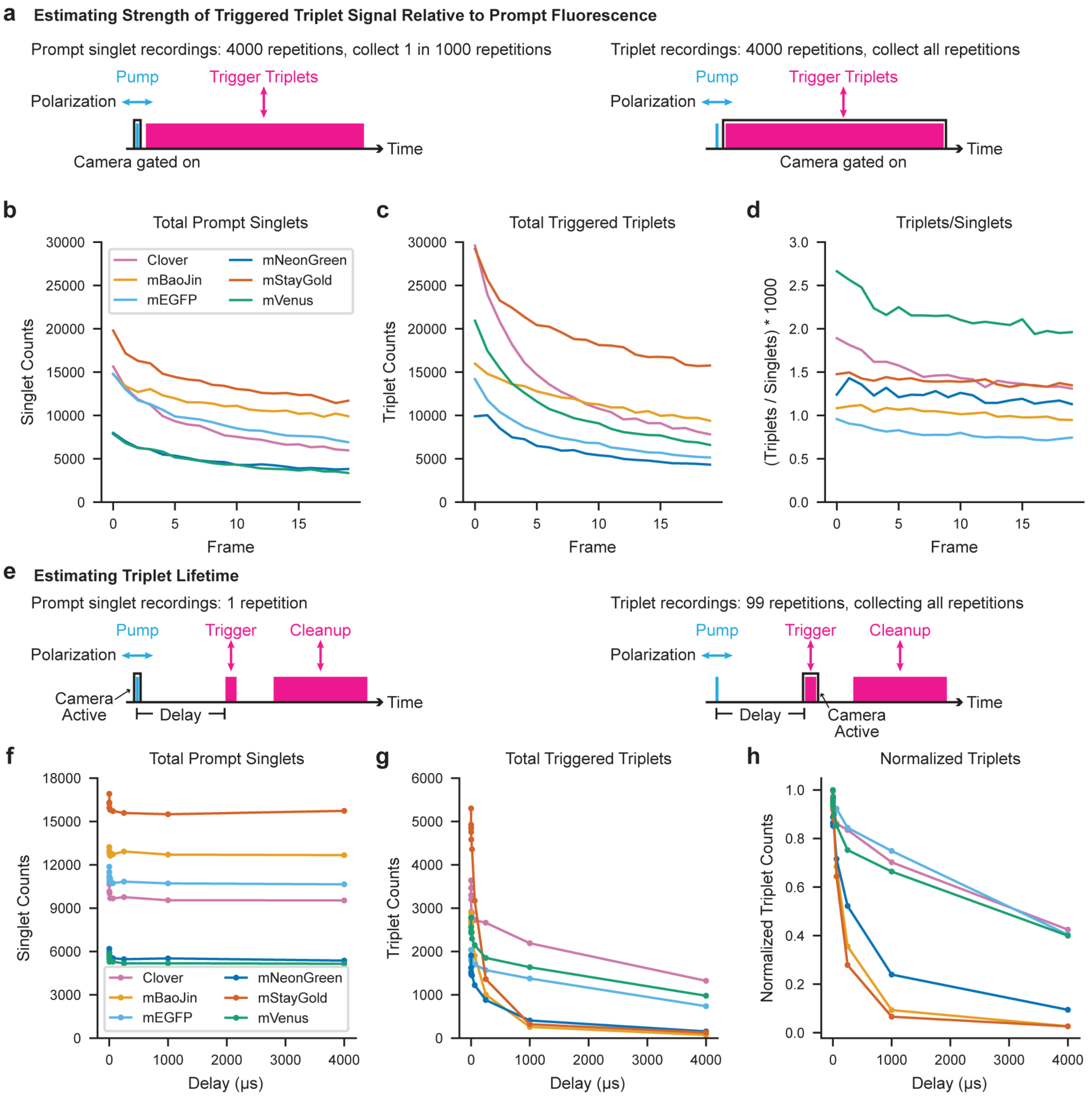


### Supplementary Figure 3. Comparing triggerable triplet properties across green fluorescent proteins.

Purified green fluorescent protein derivatives were immobilized in polyacrylamide gel for estimation of triplet state properties. (**a**) Schematic of pulse sequence used to determine relative levels of prompt singlet (fluorescence) and infrared-triggered triplet signals. A 50 ns pulse of 488 nm light (blue rectangle) was followed by a 90 μs pulse of infrared light (pink rectangle, combination of 785, 830, 915, and 940 nm) beginning 1 μs after the start of the sequence. For collection of prompt fluorescence, the camera was gated on (black outline) during every thousandth 488 nm pulse. For collection of triggered triplets, the camera was gated on during every infrared pulse. (**b**) Prompt singlet and (**c**) triggered triplet signals measured alternating over 20 frames of the scheme in (a). Photobleaching causes the signal decrease over successive frames. (**d**) Ratio of signals in (b) and (c) for each fluorescent protein. (**e**) Schematic of pulse sequence used to determine approximate triplet lifetime. A 100 ns pulse of 488 nm light (blue rectangle) was followed by a variable delay and then a 100 ns pulse of infrared light (pink rectangle, combination of 785, 830, 915, and 940 nm). A 100 μs 'clean-up' infrared pulse was delivered 4.5 ms after the initial 488 nm pump. For triggered triplet recordings, the camera was gated on (black outline) during the 100 ns infrared pulse. For prompt singlet recordings, the camera was gated on during the 100 ns blue pulse. (**f**) Total prompt singlet and (**g**) triggered triplet signal measured with the pulse scheme in (e). (**h**) Triplet signal normalized to the maximum triggered triplet level as well as the singlet level (measured every other frame) during the acquisition. Traces are recordings from individual samples.

**Supplementary Figure 4. Characterization of protein samples by SDS-PAGE and SPR**

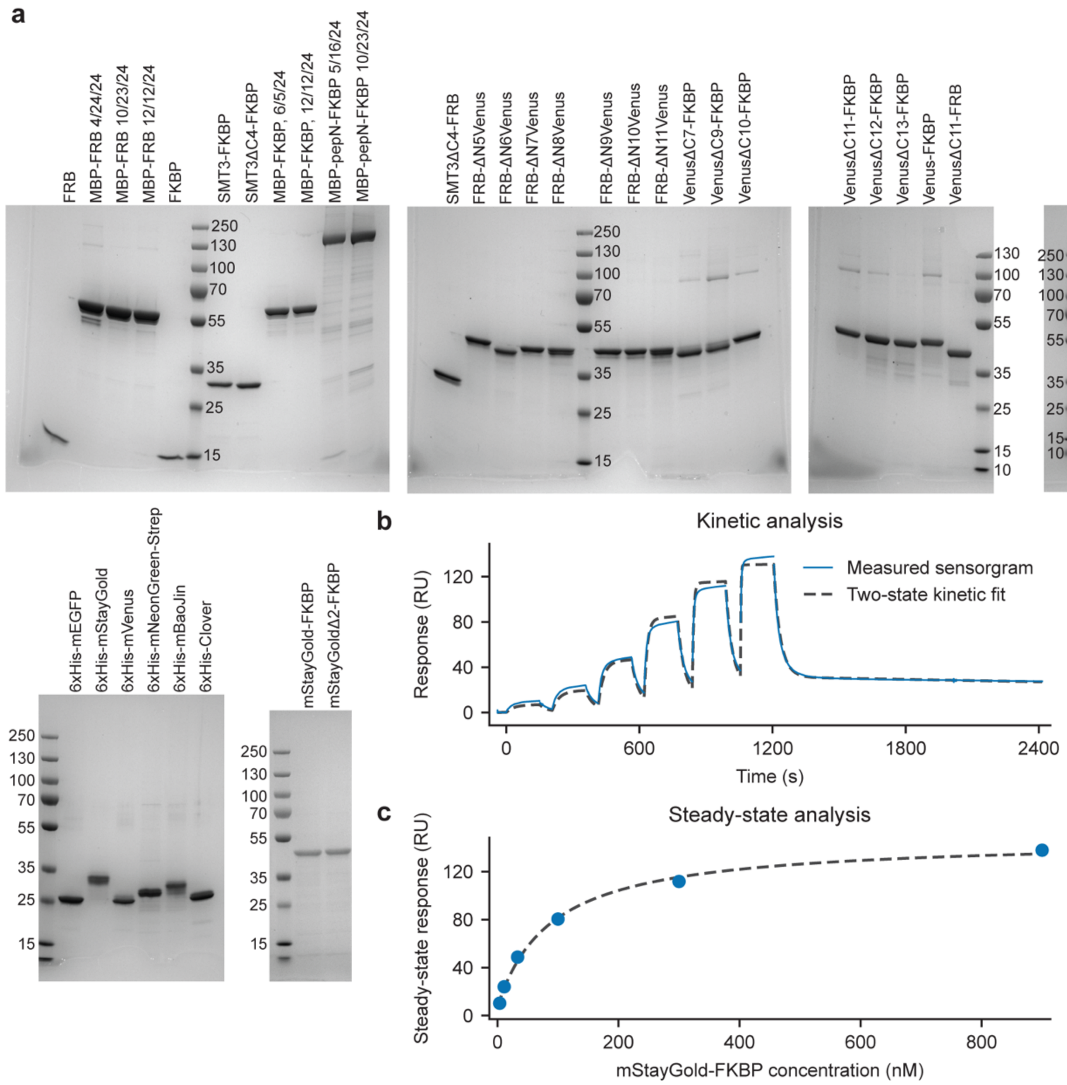


**Supplementary Figure 4. Characterization of protein samples by SDS-PAGE and SPR.** (**a**) SDS-PAGE analysis of purified protein used for tumbling experiments. Approximately 1 μg of protein per lane was loaded onto a 4-20% gradient Tris-glycine gel. Molecular weights were estimated from the PageRuler Plus Prestained Protein Ladder. Throughout the text, SMTΔC4-FRB and SMTΔC4-FKBP are referred to as SMT3-FRB and SMT3-FKBP respectively. (**b**) Measured SPR sensorgram (average of 2 channels) from addition of mStayGold-FKBP:rapamycin complexes (5 μM total rapamycin) to immobilized MBP-FRB (blue line). mStayGold-FKBP concentrations used were 3.7, 11.1, 33.3, 100, 300, and 900 nM. Dashed black line is the fit to a two-state model (kinetics chi$^2$ 8.84 RU$^2$). (**c**) Steady-state analysis of the data in (b) (blue points). Dashed black line is the fit used to determine an apparent $K_d$ of 89 nM (total offset 7.4 RU).

**Supplementary Figure 5. Shortening fluorescent tag termini improves tumbling dynamic range**

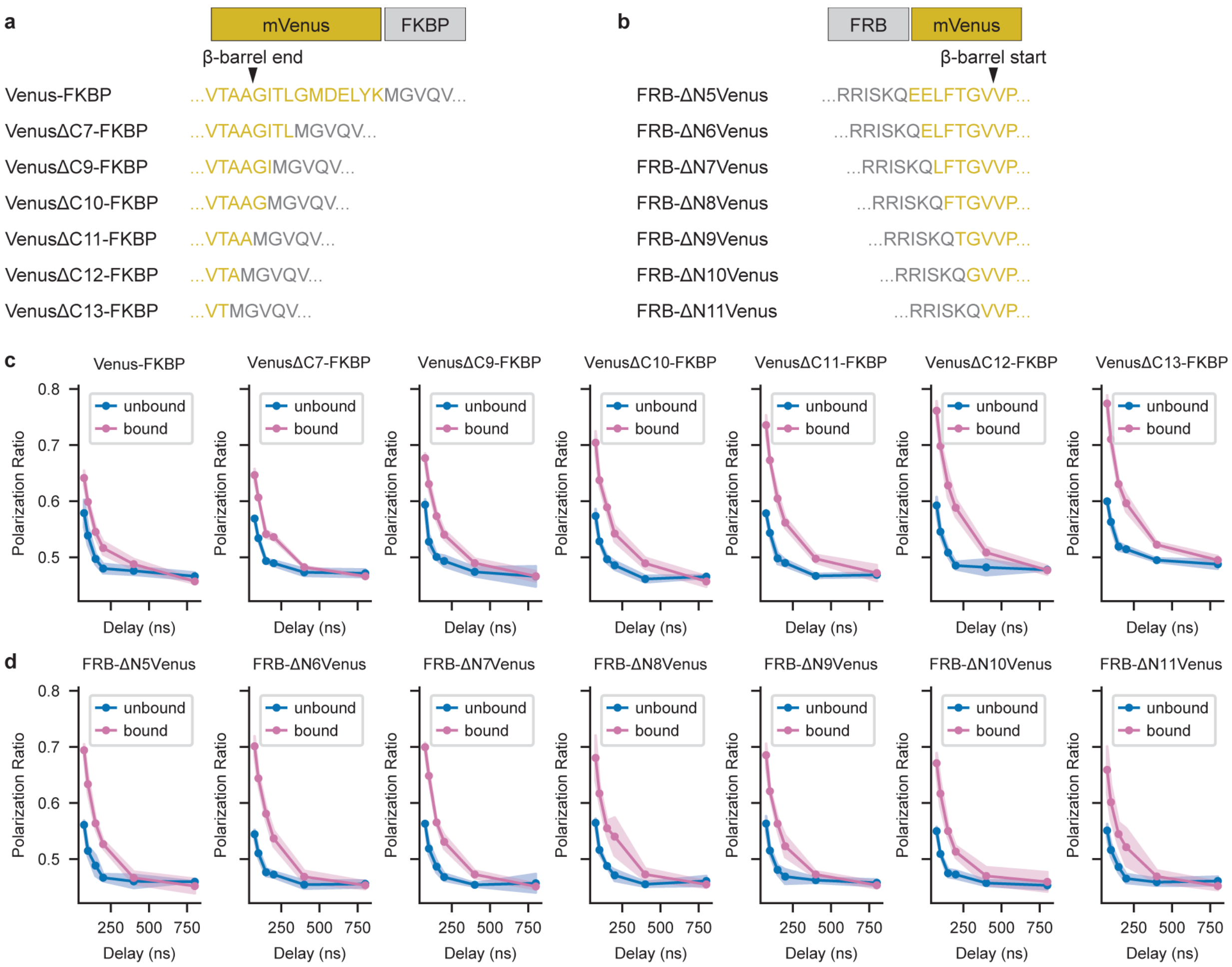


**Supplementary Figure 5. Shortening fluorescent tag termini improves tumbling dynamic range**

Partial sequences of fusion proteins with varying amounts of mVenus truncation at the (**a**) C terminus and (**b**) N terminus. (**c**) Tumbling curves measured on purified protein for the sequences in (a). 'Unbound' corresponds to measurements taken before addition of rapamycin dimerizer, and 'bound' corresponds to measurements taken at least 5 minutes after addition of rapamycin dimerizer in the presence of purified MBP-FRB. (**d**) Tumbling curves measured on purified protein for the sequences in (b), either unbound or bound to MBP-FKBP. Shaded areas indicate mean ± s.d. of n=6 samples.

**Supplementary Figure 6. Tumbling results are consistent across different fusion configurations**

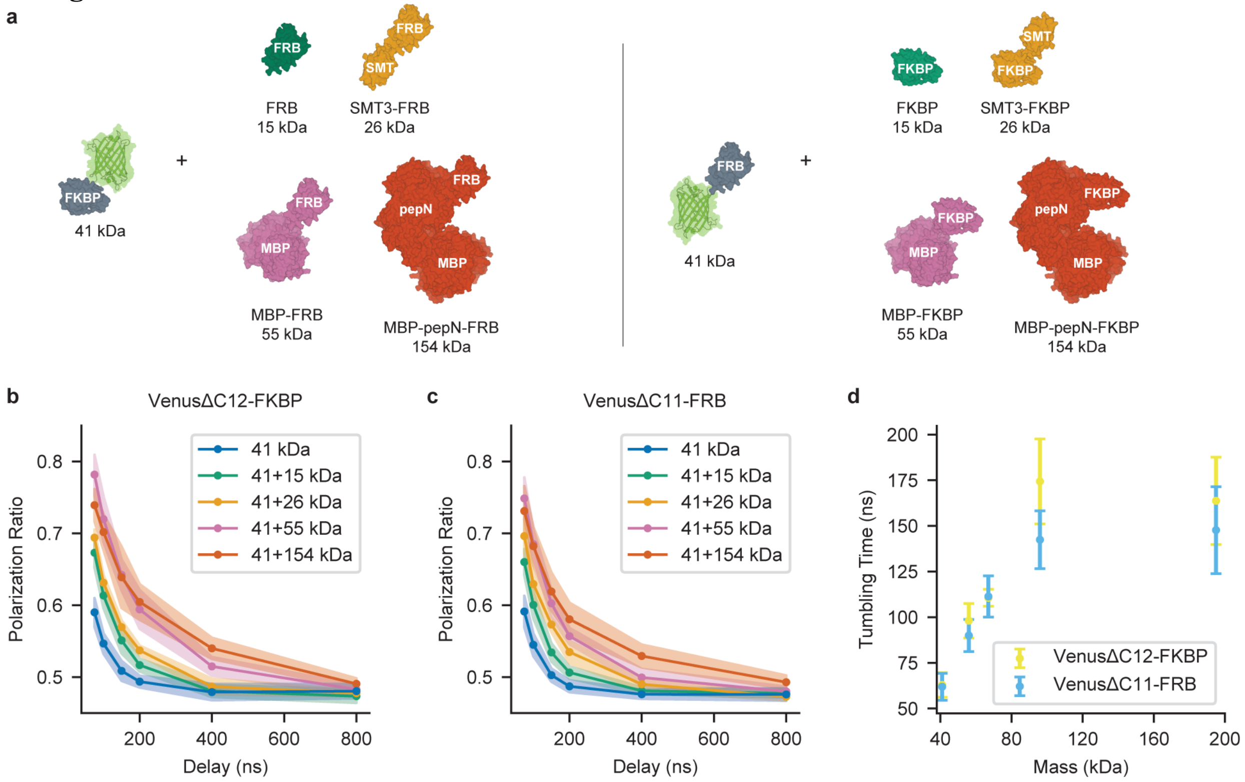


**Supplementary Figure 6. Tumbling results are consistent across different fusion configurations**

(**a**) Cartoon of purified protein components to be dimerized. mVenus was fused to either FKBP or FRB and mixed with the corresponding binding partners of different sizes. (**b**) Averaged tumbling curves for VenusΔC12-FKBP mixed with FRB-containing nonfluorescent components either before (blue, 41 kDa) or after (all other colors) addition of 10 µM rapamycin. Shaded area is mean ± s.d. of n=4-9 replicates for post-rapamycin measurements and n=28 for pre-rapamycin measurements. (**c**) Averaged tumbling curves for VenusΔC11-FRB mixed with FKBP-containing nonfluorescent components. Shaded area is mean ± s.d. of 6 replicates for all post-rapamycin measurements and 24 replicates of pre-rapamycin measurements. (**d**) Tumbling times obtained from fitting the data in (b) and (c) to a single exponential rotational diffusion model. Error bars are mean ± s.d. from fitting each recording individually.

**Supplementary Figure 7. mVenus can be used as a tumbling tag in mammalian cells**

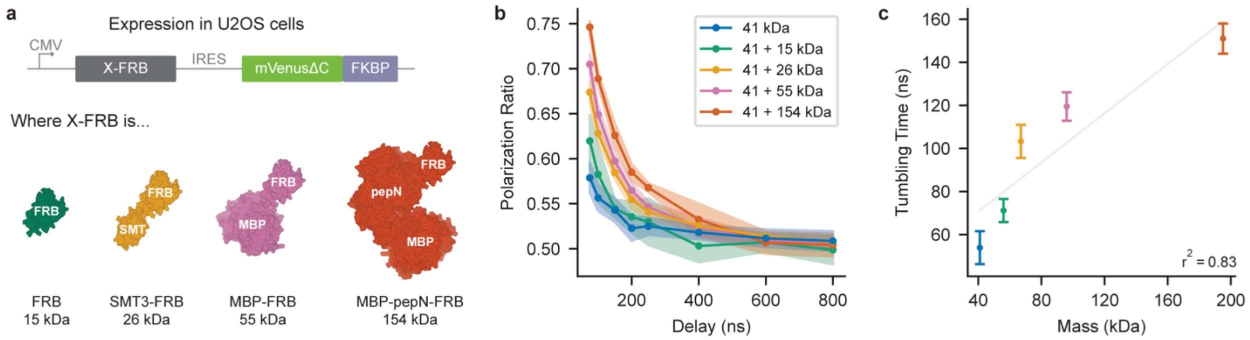


**Supplementary Figure 7. mVenus can be used as a tumbling tag in mammalian cells**
(**a**) Cartoon of expression system for introducing mVenus-tagged FKBP and an untagged FRB binding partner of varying sizes. (**b**) Averaged tumbling curves recorded from cells before (41 kDa, blue) or one hour after addition of 10 µM rapamycin (other colors). Shaded areas are mean ± s.d. of n=18 cells before rapamycin addition or n=3-5 cells for each binding partner following rapamycin addition. (**c**) Tumbling times from the data in (b). Light gray line indicates the best linear fit of tumbling time vs. mass. Points represent mean ± s.d. of n=3-5 cells; untreated cells were pooled across constructs (n=18).

**Supplementary Figure 8. Additional time course recordings of rapamycin addition**

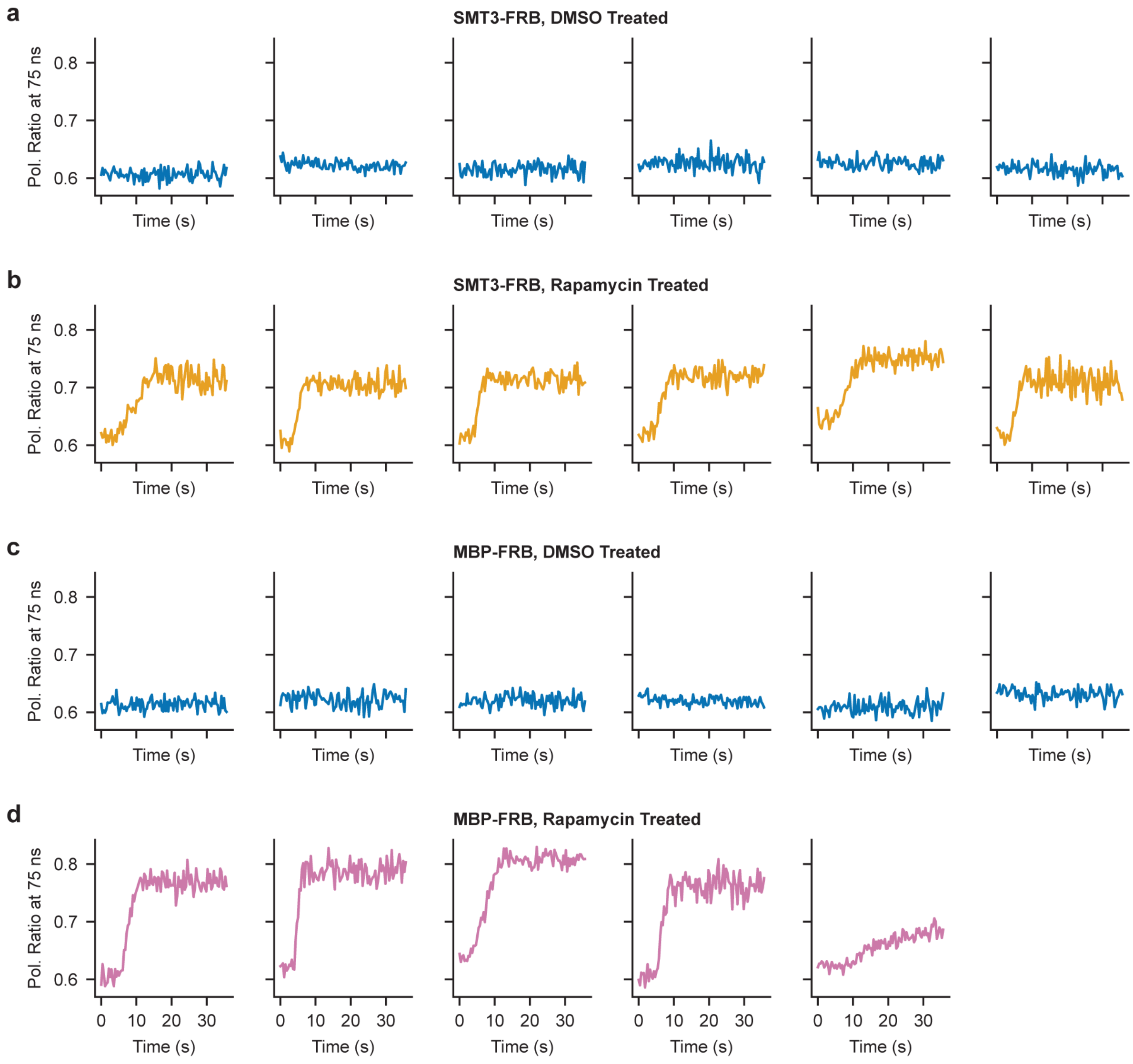


**Supplementary Figure 8. Additional time course recordings of rapamycin addition**
Polarization ratio time courses measured from individual U2OS cells expressing mStayGold-FKBP and a tagged FRB binding partner. In all time courses, media containing either DMSO or 20 µM rapamycin (10 µM final concentration) was added at frame 10 (time 3.5 seconds). (**a**) DMSO control time courses in cells expressing SMT3-FRB. (**b**) Rapamycin time courses in cells expressing SMT3-FRB. (**c**) DMSO control time courses in cells expressing MBP-FRB. (**d**) Rapamycin time courses in cells expressing MBP-FRB. Pol. ratio, polarization ratio.

**Supplementary Figure 9. Photobleaching during TTM time lapse imaging**

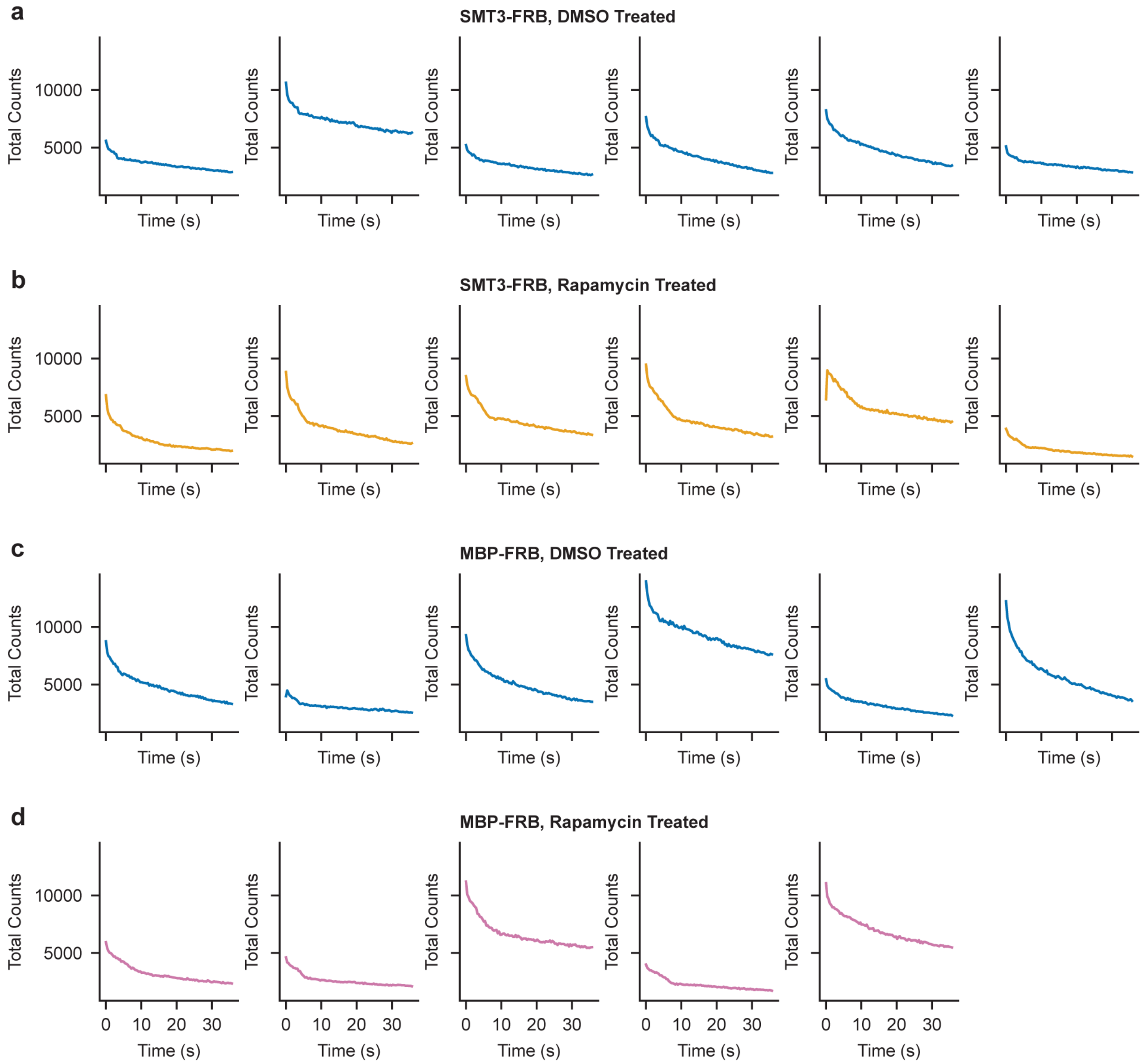


**Supplementary Figure 9. Photobleaching during TTM time lapse imaging**
Single-trial total photon count (sum of parallel and perpendicular spots) recordings of rapamycin addition in live cells (same dataset as **Supplementary Figure 8**). Each trace is from an individual U2OS cells expressing mStayGold-FKBP and a tagged FRB binding partner. In all traces, media containing either DMSO or 20 μM rapamycin (10 μM final concentration) was added to the samples at frame 10 (time 3.5 seconds). (**a**) DMSO control recordings in U2OS cells expressing SMT3-FRB. (**b**) Rapamycin recordings in cells expressing SMT3-FRB. (**c**) DMSO control recordings in cells expressing MBP-FRB. (**d**) Rapamycin recordings in cells expressing MBP-FRB.

**Supplementary Figure 10. Tumbling curves and fit components for p53 tumbling in U2OS cells.**

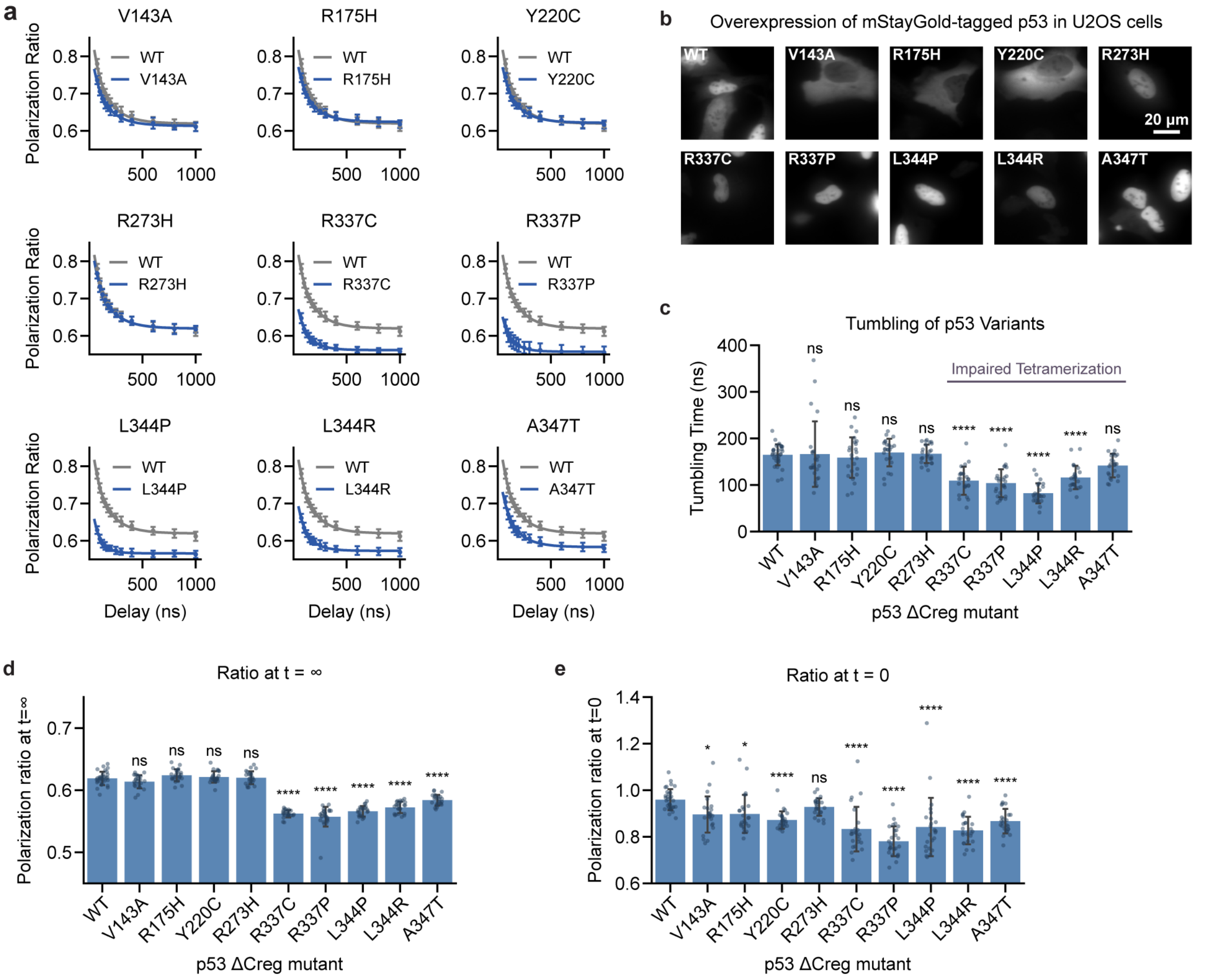


**Supplementary Figure 10. Tumbling curves and fit components for p53 tumbling in U2OS cells.**

(**a**) Averaged tumbling curves for p53 $\Delta$Creg expressed in U2OS cells. Solid line indicates fit to a single exponential rotational diffusion model. Wild type p53 data (gray) is shown on each plot. For WT, R273H, L344P, and A347T, data are reproduced from **Figure 4**. (**b**) Representative widefield fluorescence images of U2OS cells overexpressing mStayGold-tagged p53. Tumbling was measured in the cytoplasm for cytosolically localized variants (V143A, R175H, Y220C); for all other constructs, tumbling was measured in the nucleus. (**c**) Tumbling times from fits to individual cells in (a). Asterisks indicate results of Dunnett's post hoc tests following one-way ANOVA $F(9, 240) = 25.6$, $p<0.0001$. (**d**) Polarization ratio at long times ($\rho_\infty$) for the data in (a). Higher $\rho_\infty$ is consistent with increased homo FRET between mStayGold tags, as FRET scrambling of orientations will lead to less polarization alignment with the IR beam. Asterisks indicate results of Dunnett's post hoc tests following one-way ANOVA $F(9, 240) = 193$, $p<0.0001$. (**e**) Polarization ratio at t=0 ($\rho_0$) for the data in (a). Asterisks indicate Dunnett's post hoc tests following one-way ANOVA $F(9, 240) = 9.5$, $p<0.0001$. Error bars are mean $\pm$ s.d. of n=24-32 cells; points are individual cells.

### Supplementary Figure 11. Crosslinked Western blots reveal extent of p53 homo-oligomerization

**a** Total p53 (No Crosslinking) Western blots. anti-FLAG, anti-GAPDH.

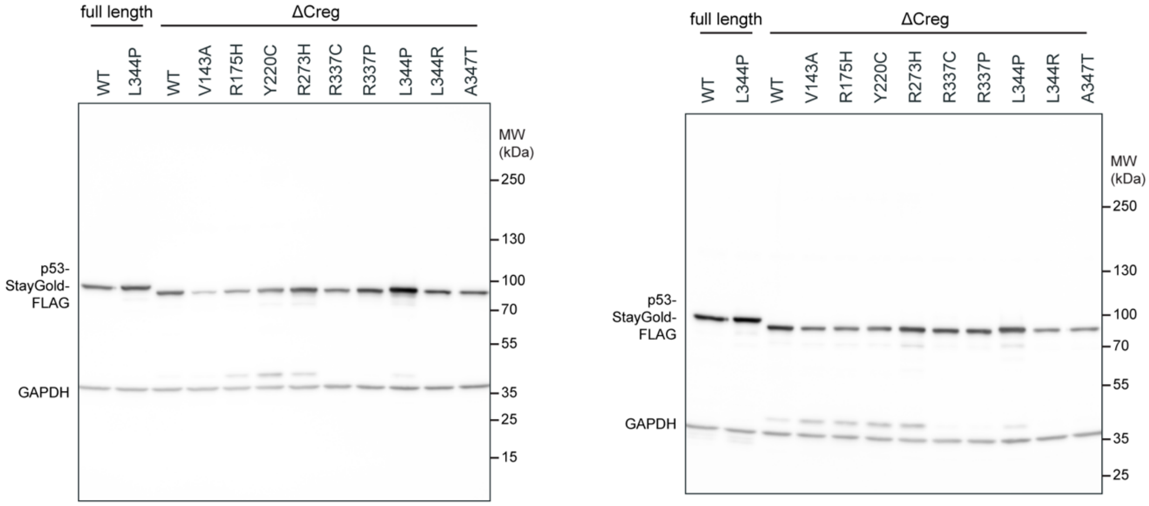


**b** p53 Crosslinked Western blots. anti-FLAG, anti-GAPDH.

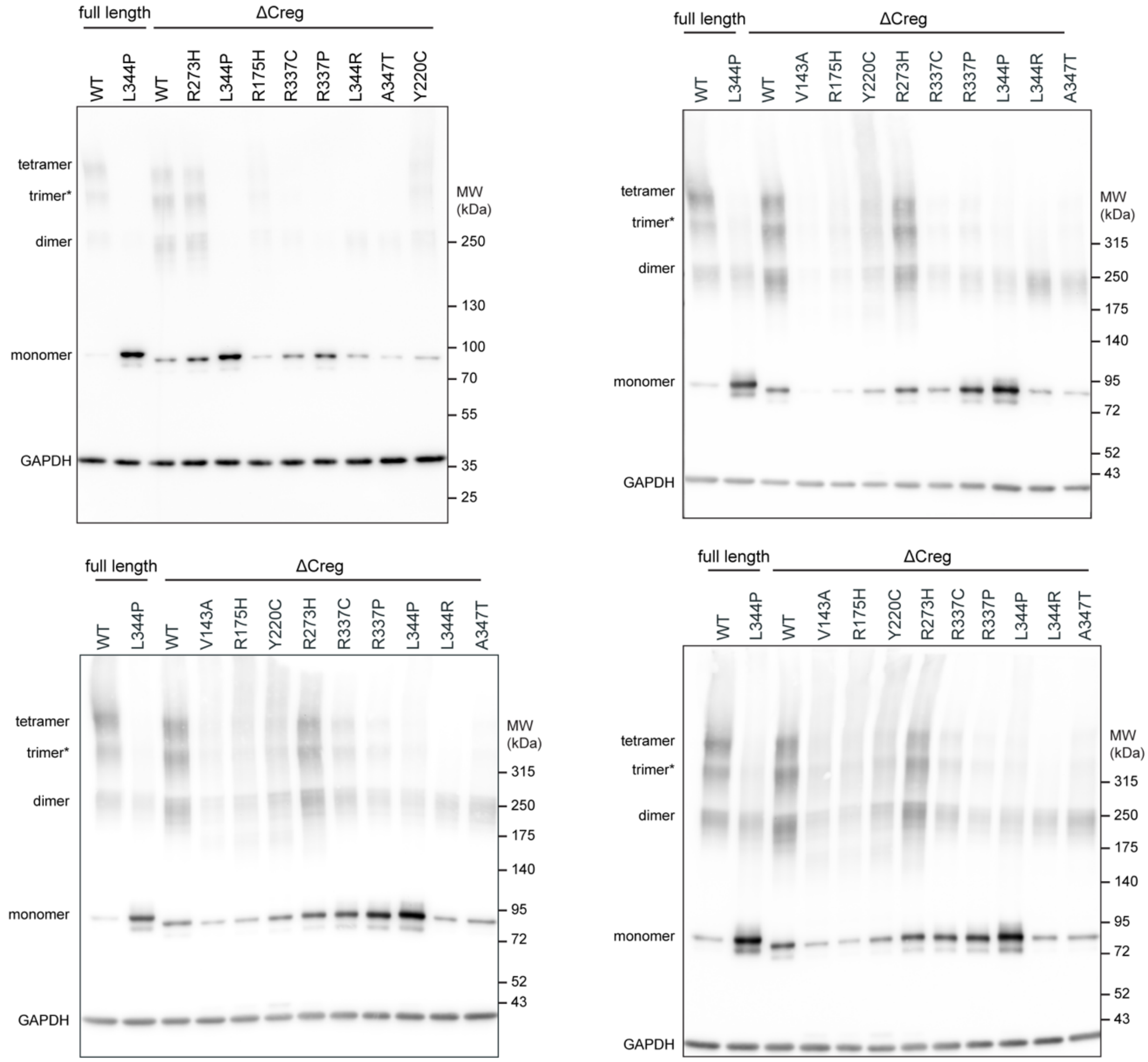

**Supplementary Figure 11. Crosslinked Western blots reveal extent of p53 homo-oligomerization**

(**a**) Western blots of lysate from U2OS cells transiently overexpressing p53 mutants. (**b**) Crosslinked Western blots of U2OS cells transiently overexpressing p53 mutants. Cells were crosslinked with 2 mM disuccinimidyl glutarate at room temperature for 10 minutes before lysis. Note that lane order and ladder differs between blots. The blot in **Figure 4** used the same samples that are shown in lanes 3, 7, 8, 9, 10, 11, and 12 of the lower right blot in (b).

**Supplementary Figure 12. FSEC analysis of p53 homo-oligomerization**

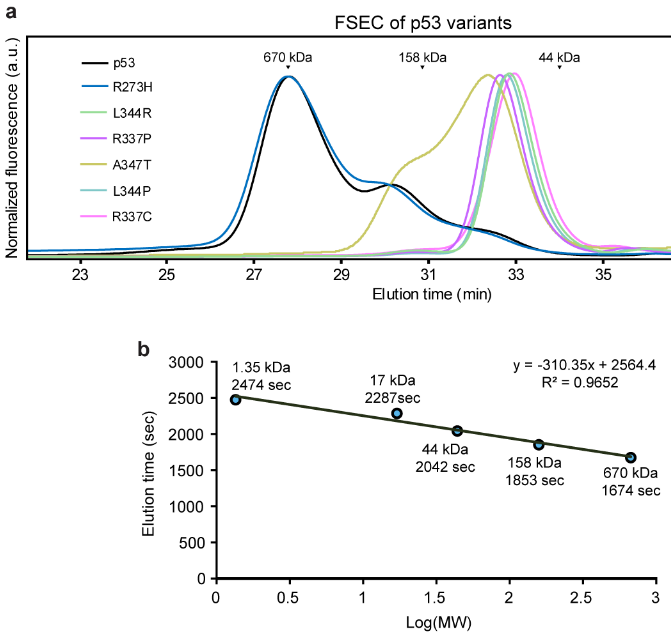


**Supplementary Figure 12. FSEC analysis of p53 homo-oligomerization**

(**a**) FSEC of U2OS lysates expressing mStayGold-tagged p53. The three peaks are tentatively assigned to tetramers, dimers, and monomers (left-to-right). p53 tetramers have previously been reported to run at higher molecular weight than expected in chromatography assays.[21] Arrows indicate elution times of molecular weight references. All traces were normalized to their highest peak. Note that that the NP-40 lysis conditions used here have been reported to leave the nuclear envelope intact,[22] so these traces predominantly reflect the cytosolic p53 fraction. (**b**) Elution times plotted as a function of the logarithm of molecular weight standards (kDa, Bio-Rad). A line of best fit is shown.

**Supplementary Figure 13. Fluorescence anisotropy indicates homo-FRET between p53 tags.**

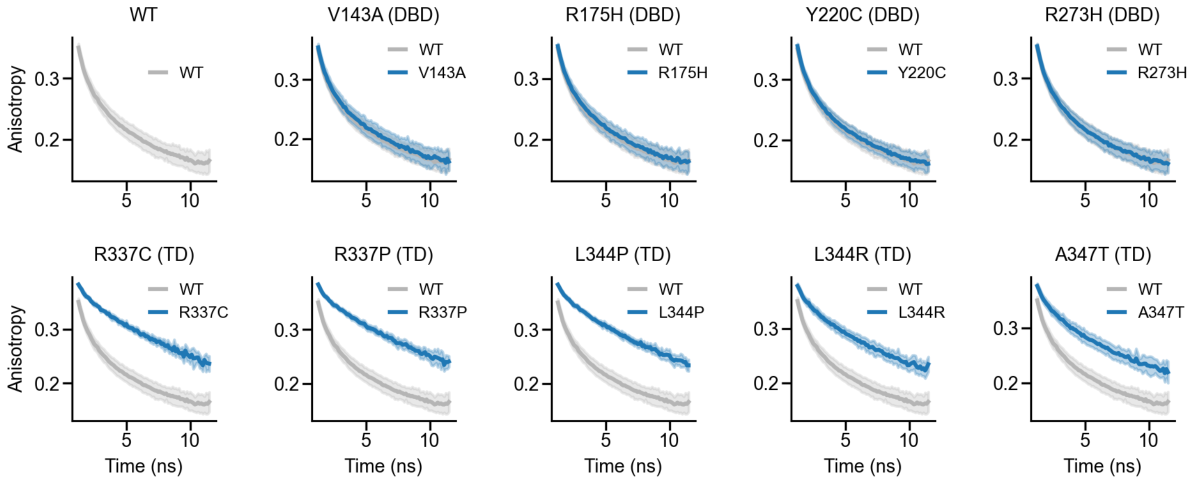


**Supplementary Figure 13. Fluorescence anisotropy indicates homo-FRET between p53 tags.**
Fluorescence anisotropy curves collected for U2OS cells expressing the indicated p53 (1-363, Creg truncation) mutant. The same wild type data is overlaid on each mutant for ease of comparison. Anisotropy traces were summed over the entire image and then averaged across 17-24 images per mutant. Lines and shaded areas indicate mean ± s.d. DBD, DNA binding domain; TD, tetramerization domain.

**Supplementary Figure 14. TTM reveals differences between wild type and L344P in full length p53.**

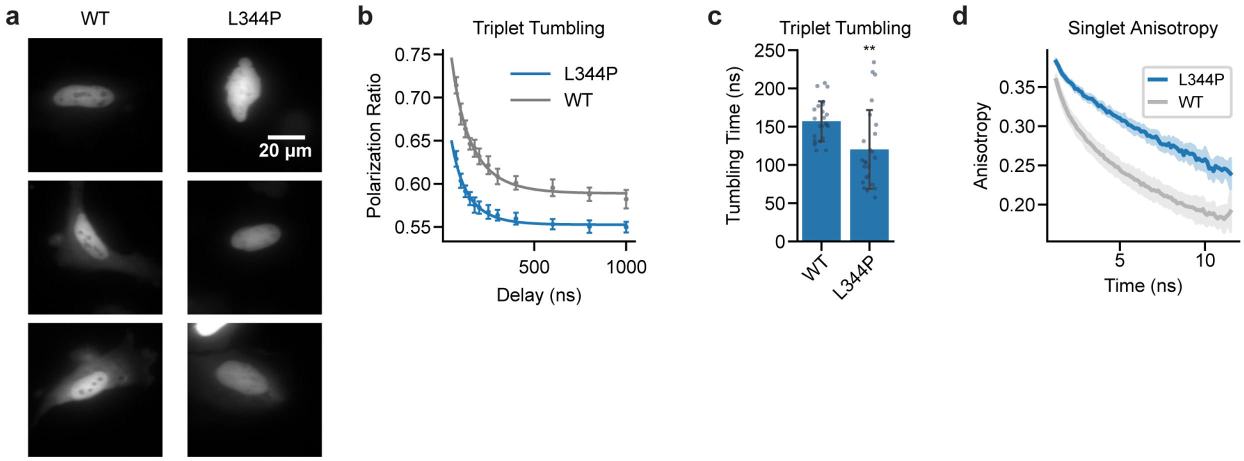


**Supplementary Figure 14. TTM reveals differences between wild type and L344P in full length p53.**
(**a**) Representative widefield fluorescence images of U2OS cells transiently overexpressing full length p53 wild type or L344P mutant. For both constructs, fluorescence was primarily observed in the nucleus. (**b**) Tumbling curves for tagged wild type and L344P mutant p53, measured from the nucleus of U2OS cells. Points indicate mean ± s.d. (n=24 cells per genotype). Solid line indicates the fit to a single exponential rotational diffusion model. (**c**) Tumbling times for per-cell fits to the data in (b). Bar indicates mean ± s.d. (**d**) Averaged fluorescence (singlet) anisotropy. Fluorescence was summed across the entire field of view (containing 1-3 cells) to generate a global anisotropy decay. Lines and shaded areas indicate mean ± s.d. (n=18 anisotropy decays).

**Supplementary Figure 15. Levels of E6AP and E6 expressed in HEK293 cells**

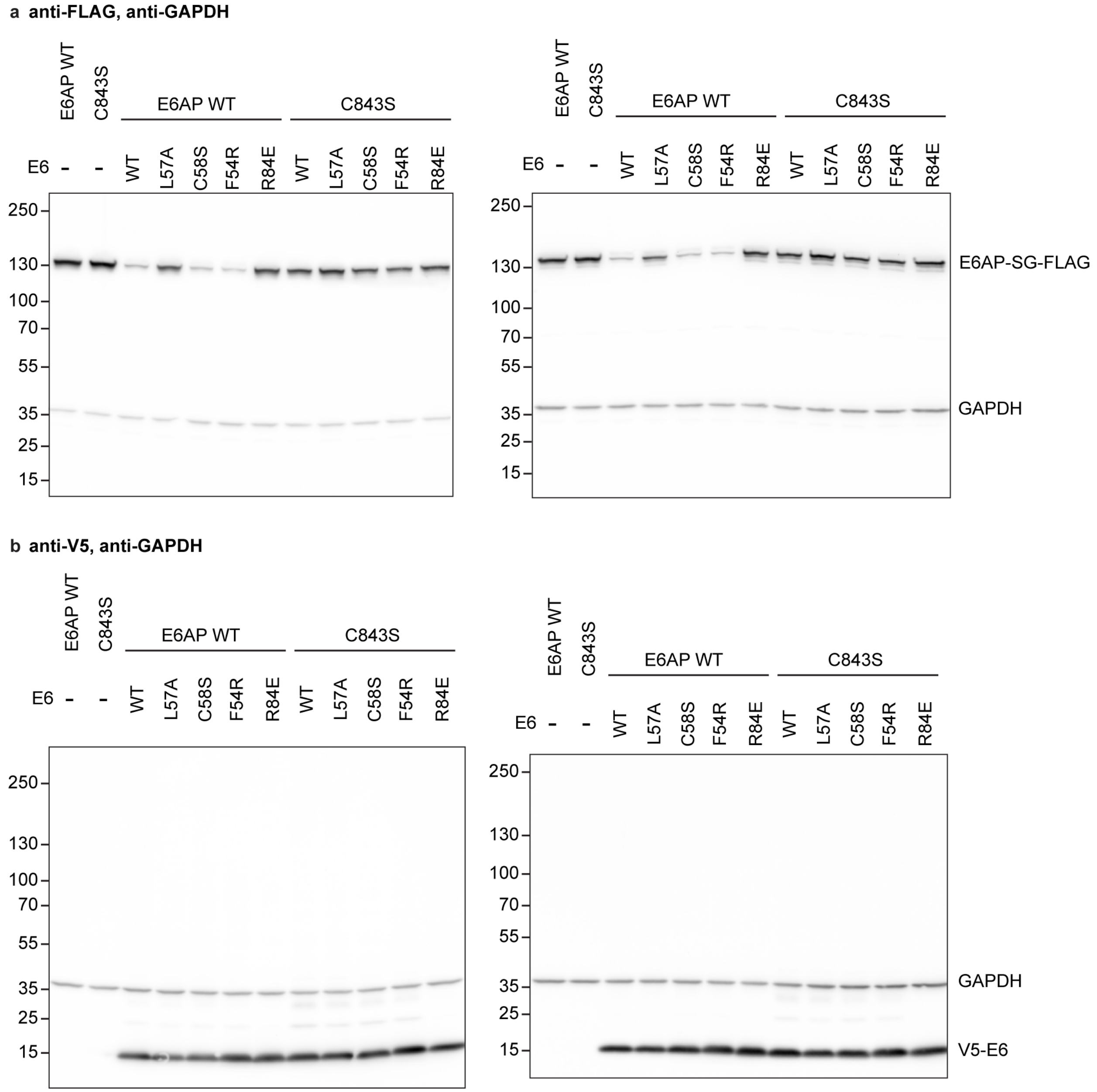


**Supplementary Figure 15. Levels of E6AP and E6 expressed in HEK293 cells**
(**a**) Western blots of HEK293 lysates overexpressing E6AP+E6 constructs and blotted for FLAG (E6AP tag) and GAPDH. (**b**) Western blots of HEK293 lysates overexpressing E6AP+E6 constructs and blotted for V5 (16E6 tag) and GAPDH. In both (a) and (b), left and right blots are biological replicates of each other.

### Supplementary Figure 16. Tumbling of E6AP WT and C843S

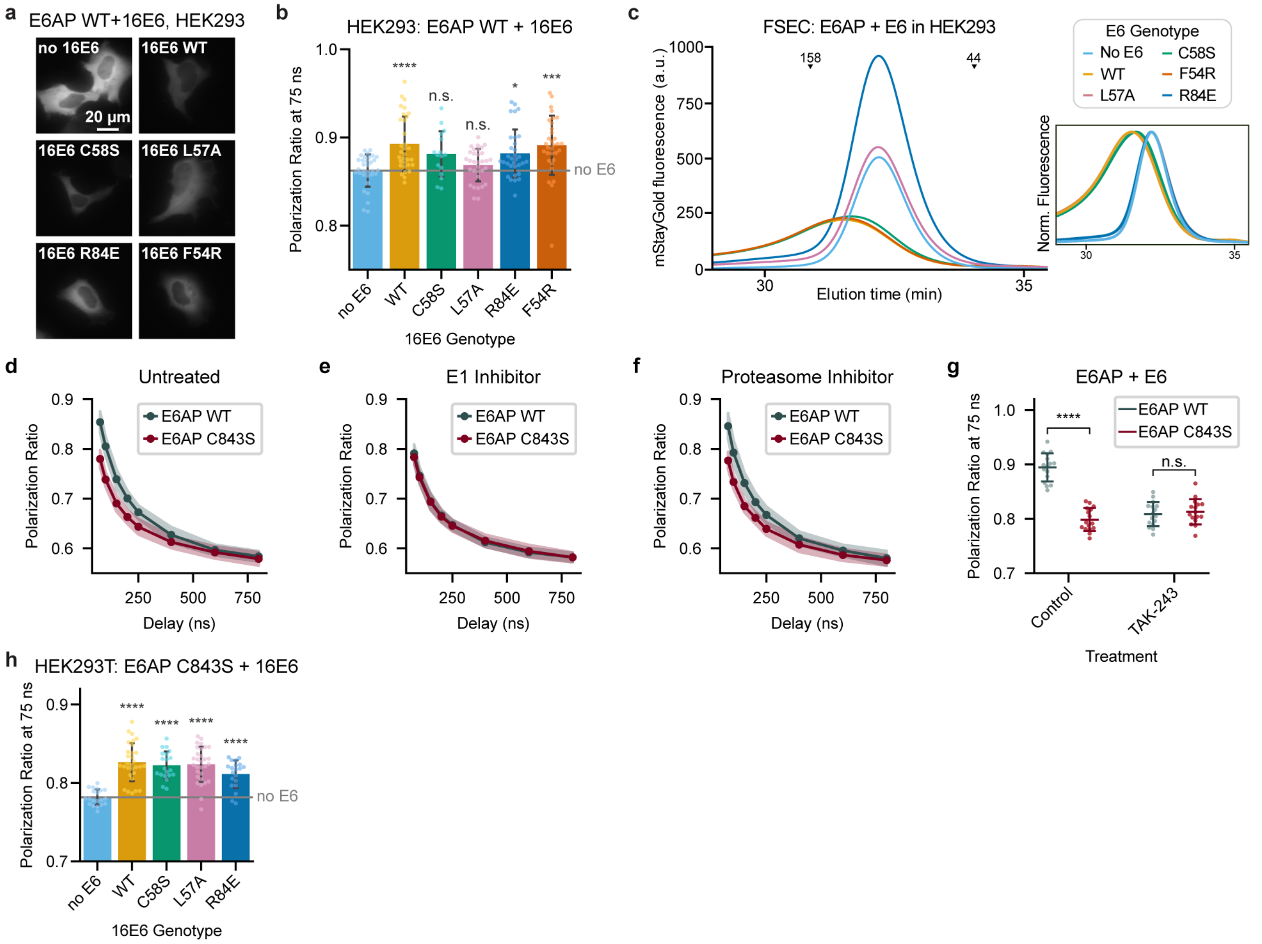


**Supplementary Figure 16. Tumbling of E6AP WT and C843S**
(**a**) Representative widefield fluorescence images of cells HEK293 expressing E6AP WT and 16E6 variants. (**b**) Polarization ratios at 75 ns delay in HEK293 cells expressing wild type E6AP-mStayGold with the indicated 16E6 variant. Bars represent the mean ± s.d. (n=16-32 cells). Asterisks indicate results of Dunnett post-hoc tests relative to the no 16E6 control, following one-way ANOVA ($F(5,170)=6.6$, $p<0.0001$). (**c**) FSEC of E6AP WT and 16E6 constructs. To counteract the reduced fluorescence, four times more lysate was loaded for binding competent 16E6 (WT, C58S, F54R) than reduced affinity 16E6 (L57A, R84E). Inset shows the same data with each trace normalized to its maximum. (**d-f**) Tumbling curves for E6AP WT and C843S without E6 in (d) untreated HEK293, (e) HEK293 treated with 1 µM TAK-243 for 4 hours, or (f) HEK293 treated with 1 µM carfilzomib for 4 hours. Shaded area represents mean ± s.d. (n=24-40 cells). (**g**) HEK293 expressing E6AP WT or C843S and 16E6 were treated with either 0.1% DMSO control or 1 µM TAK-243 for 4 hours. Asterisks indicate results of Student's t-test; bars indicate mean ± s.d (n=16 cells). (**h**) Polarization ratios at 75 ns delay of HEK293T cells expressing catalytically dead E6AP C843S-mStayGold with the indicated 16E6 variant. Bars represent the mean ± s.d. (n=20-30 cells). Asterisks indicate results of Dunnett post-hoc tests relative to the no 16E6 control, following one-way ANOVA ($F(4,115)=19.3$, $p<0.0001$).

**Supplementary Figure 17. E6AP isoforms 1 and 2 have similar tumbling, regardless of localization**

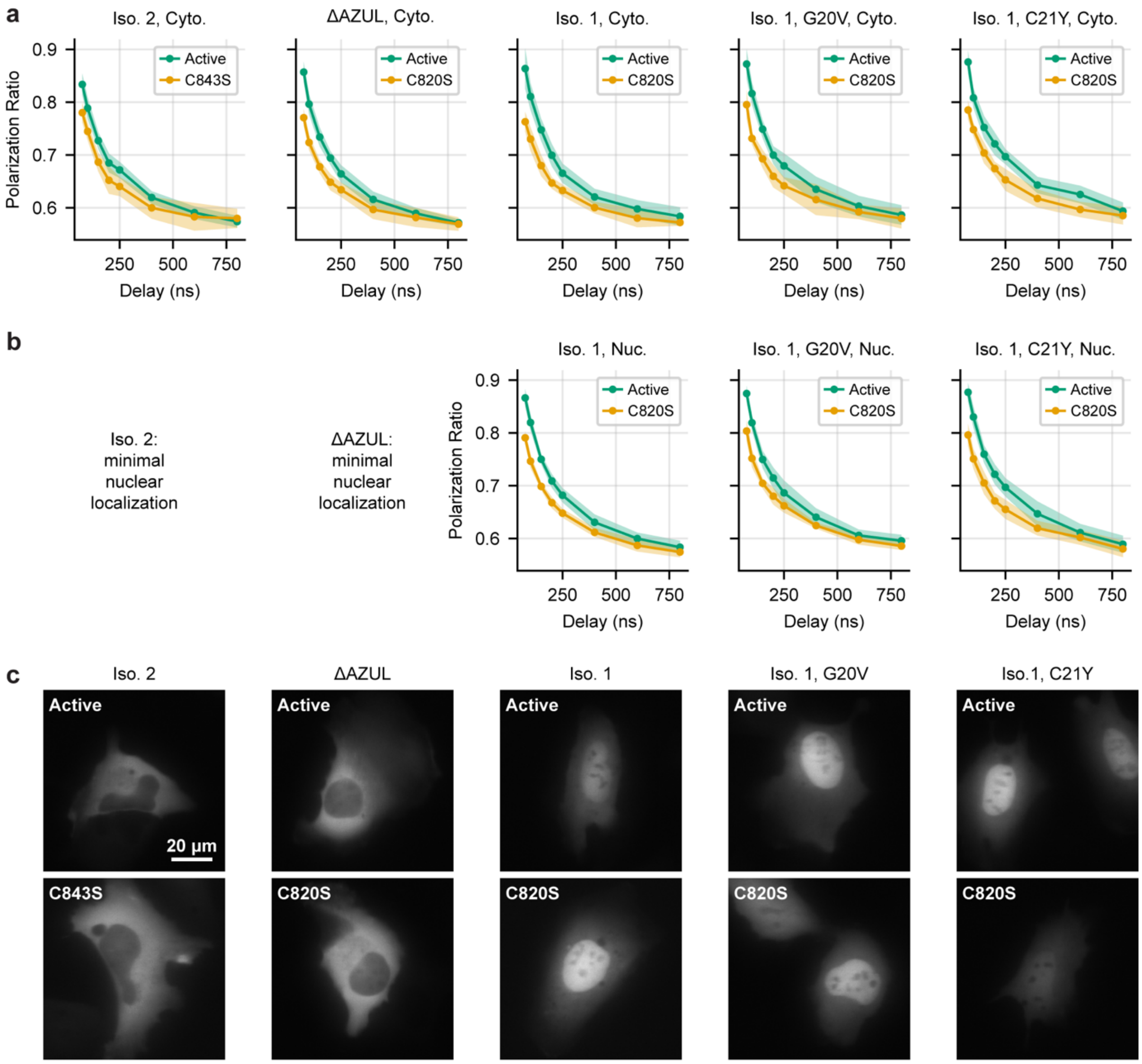


**Supplementary Figure 17. E6AP isoforms 1 and 2 have similar tumbling, regardless of localization.**
(**a**) Tumbling curves measured in the cytoplasm of U2OS cells for active E6AP or E6AP with a mutated active site cysteine (C820S or C843S, catalytically dead). (**b**) Tumbling curves measured in the nucleus for E6AP variants where there was substantial nuclear localization. (**c**) Representative images of E6AP variants. Note that the ratio of nuclear to cytoplasmic fluorescence, as well as the total brightness, varied between cells for a given construct. Iso. 2: isoform 2, iso. 1: isoform 1, cyto.: cytoplasm, nuc.: nucleus. G20V and C21Y mutations in the AZUL domain were reported to affect E6AP:proteasome interactions.[23] Isoform 1 numbering was used for the AZUL domain deletion mutant. Where possible, both nuclear and cytoplasmic measurements were made on the same cell. Points and shaded areas are mean ± s.d. (n=5-13 cells).

**Supplementary Figure 18. C843S mutation does not affect E6AP co-immunoprecipitation profile**

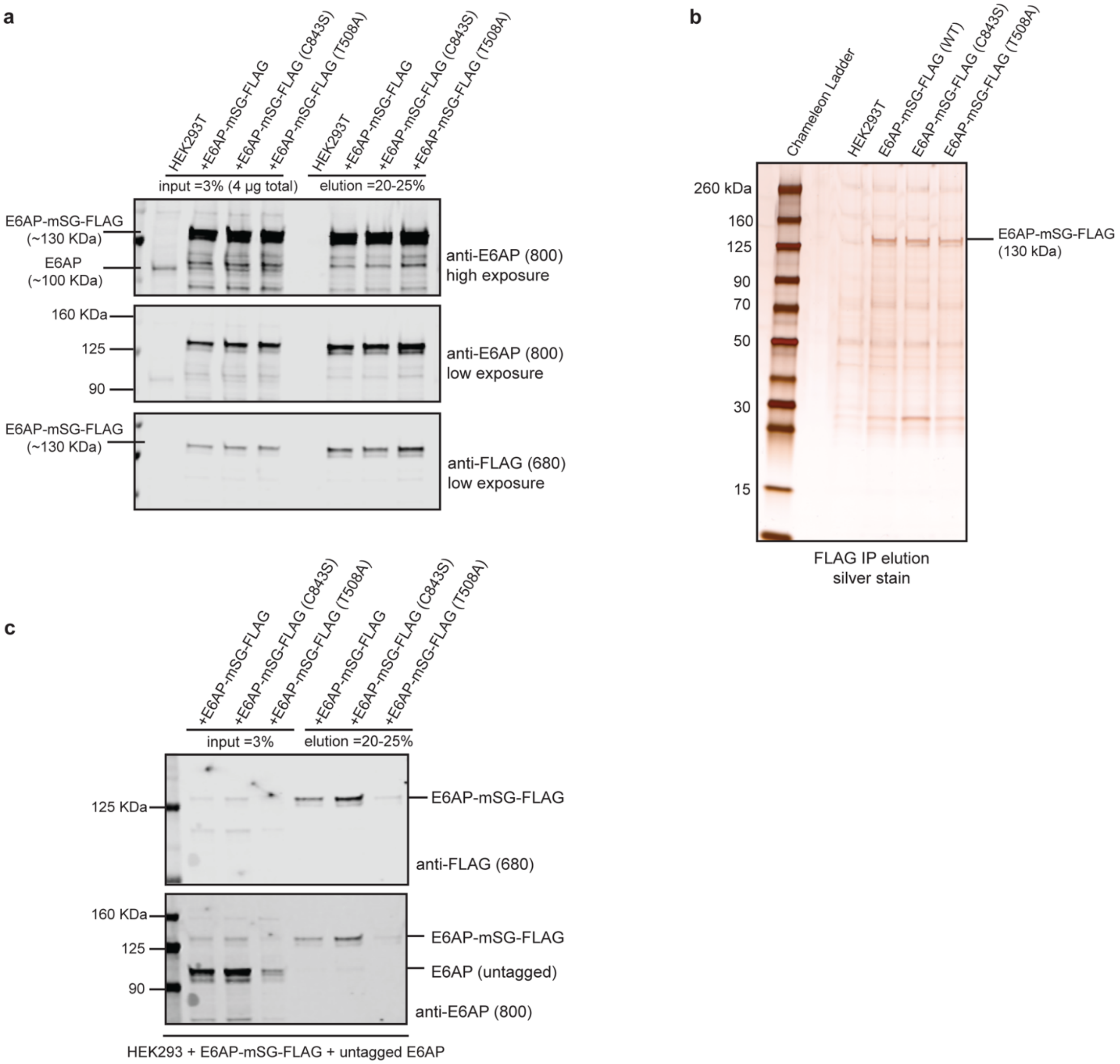


**Supplementary Figure 18. C843S mutation does not affect E6AP co-immunoprecipitation profile**
(**a-b**) FLAG immunoprecipitation was performed on cells expressing mStayGold-FLAG-tagged E6AP WT, C843S, and T508A. FLAG IP inputs and eluates were run on SDS-PAGE gel and a) immunoblotted with E6AP and FLAG antibodies or b) silver stained. a) Endogenous E6AP does not associate with tagged E6AP and b) no protein associations are visibly different in WT versus mutant E6AP proteomes. (**c**) FLAG IP was performed on cells expressing mStayGold-FLAG-tagged E6AP and untagged E6AP. No dimerization is apparent between tagged and untagged proteins. mSG: mStayGold.

**Supplementary Figure 19. Non-reducing Western blots of E6AP wild type and C843S**

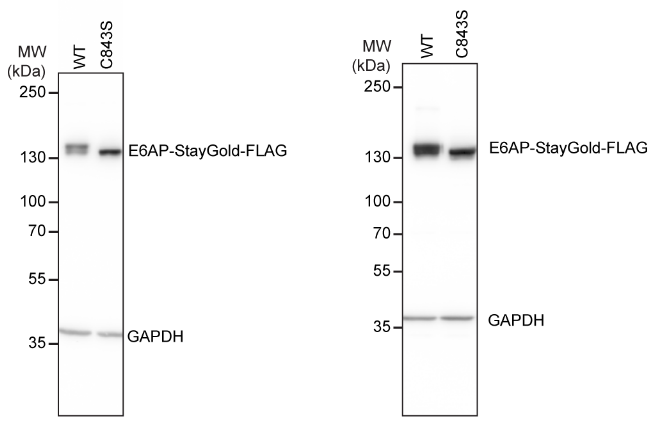


**Supplementary Figure 19. Non-reducing Western blots of E6AP wild type and C843S**
Non-reducing Western blots of E6AP wild type and C843S (catalytically dead). A shift to higher molecular weight is seen in E6AP WT, consistent with ubiquitin loading on the active cysteine or another thiol-dependent modification. Corresponding reducing Western blots can be seen in **Supplementary Figure 15**. Left and right blots are biological replicates. Samples were blotted for FLAG tag and GAPDH.

**Supplementary Table 1. TTM acquisition parameters**

| Data type | Cycles per exposure | Cycle period (μs) | 488 nm width (ns) | IR trigger width (ns) | IR cleanup width (μs) | Exposures at each delay per recording | Intensifier voltage (V) |
|---|---|---|---|---|---|---|---|
| mStayGold coated beads | 4990 | 200 | 50 | 50 | 49 | 6 | 850 |
| Rapamycin dimerization in vitro | 39920 | 25 | 30 | 30 | 20 | 4 | 850-900 |
| Rapamycin in cells | 10000 | 25 | 30 | 30 | 20 | 2 | 800-900 |
| Rapamycin in cells – time course | 10000 | 25 | 30 | 30 | 20 | 1 | 800-900 |
| p53 and E6AP (full tumbling curves) in cells | 10000 | 25 | 30 | 30 | 20 | 2 | 800 |
| E6+E6AP in cells (75 ns delay only) | 10000 | 25 | 30 | 30 | 20 | 10 | 800 |

**Supplementary Table 1. TTM acquisition parameters**

Acquisition parameters used for TTM in various samples using the pulse scheme indicated in **Figure 1**. Cycles per exposure refers to the number of 488-IR pulse pairs within a single camera exposure. Where multiple replicates were acquired, delays were taken first in ascending order and then in descending order, alternating for successive exposures. Total acquisition time for beads and in vitro dimerization measurements was 1 second per delay; for cellular measurements total acquisition time per delay was 250 ms.

**Supplementary Table 2. Comparison of predicted and measured tumbling times**

| Bead diameter (nm) | Experimental θ (μs) | Predicted θ in pure water (μs) | Predicted θ with 1.3x higher viscosity (μs) | Experimental CV of diameter | Reported CV of diameter |
|---|---|---|---|---|---|
| 26 | 3.0 | 2.3 | 3.0 | 40% | 15.5% |
| 37 | 8.5 | 6.6 | 8.5 | 29% | 12.2% |
| 60 | 39.9 | 27.9 | 36.3 | 3% | 13.8% |

**Supplementary Table 2. Comparison of predicted and measured tumbling times**
Experimentally determined tumbling times were modeled as a log normal distribution with median θ and log-width σ as described in **Supplementary Note 1**. Predicted rotational diffusion coefficients (D) for spheres in pure water at 298 K and 1 atm of pressure (viscosity of 0.89 mPa·s) were calculated from the Stokes-Einstein-Debye equation ($D = \frac{k_B T}{8\pi\eta r^3}$, where $k_B$ is Boltzmann's constant, T is temperature, $\eta$ is viscosity, and r is the radius of the particle).[24] Rotational correlation time was defined as 1/6D, as described for spherical rotors by the Perrin equation. The samples measured were latex beads at 4% solids, which is concentrated enough to be milky white; their viscosity is likely higher than that of pure water. The experimentally determined tumbling times are on average 30% higher than of those predicted for pure water; we calculate the Stokes-Einstein-Debye predictions for a solution 30% more viscous than water for comparison. Experimentally determined coefficient of variation (CV) was calculated from the fit σ and compared to the reported CV on the supplier's specification sheet (rightmost column). Some additional aggregation was observed from addition of the fluorescent protein to the beads, which would increase CV above the manufacturer's specifications. Note that bead diameters here are actual diameters from specification sheet and differ slightly from diameters in product name.

**Supplementary Table 3. Reagents and Resources**

| Name | Supplier | Part Number |
|---|---|---|
| Fluorescent Standards | | |
| Upconversion Nanoparticles, NaYF4,Yb,Tm@NaYF4,Yb,Nd, PEG-COOH modified core-shell, fluorescence λex 808 nm, blue light | Millipore Sigma | 926574-500UL |
| CML latex beads, 4% w/v, 20 nm diameter | Thermo Fisher | C37231 |
| CML latex beads, 4% w/v, 40 nm diameter | Thermo Fisher | C37232 |
| CML latex beads, 4% w/v, 60 nm diameter | Thermo Fisher | C37233 |
| Fluorescein | Millipore Sigma | 46955 |
| Molecular Biology | | |
| NEB5alpha competent E. coli (high efficiency) | New England Biolabs | C2987I |
| NEB HiFi Assembly Master Mix | New England Biolabs | E2621L |
| Q5 High Fidelity 2x Master Mix | New England Biolabs | M0492L |
| Restriction enzymes (DpnI, BamHI, EcoRI, SpeI, XhoI) | New England Biolabs | various |
| GeneJet plasmid miniprep kit | Thermo Fisher | K0503 |
| Carbenicillin, disodium salt | Sigma Aldrich | C1389-1G |
| Protein Purification & *in vitro* Measurements | | |
| BL21(DE3) competent E. coli | New England Biolabs | C2527I |
| NEBExpress Ni resin | New England Biolabs | S1428S |
| Strep-Tactin Superflow Plus | Qiagen | 30004 |
| 10x BugBuster protein extraction reagent | Millipore | 70921-3 |
| Pierce Protein Concentrator, PES, 10K MWCO, 0.5 mL | Thermo Fisher | 88513 |
| Novex Tris-glycine 4-20% gradient gel | Thermo Fisher | XP04205BOX |
| GelCode Blue stain reagent | Thermo Fisher | 24590 |
| Ammonium persulfate | Sigma-Aldrich | CAS 7727-54-0 |
| TEMED | Sigma-Aldrich | CAS 110-18-9 |
| Acrylamide/Bis-acrylamide, 30% solution | Sigma-Aldrich | A3699-100ML |
| D-desthiobiotin | Millipore Sigma | CAS 533-48-2 |
| Imidazole | Fisher Scientific | O3196 (CAS 288-32-4) |
| Bovine Serum Albumin, lyophilized powder | Sigma | A2153-100G |
| Glycerol, UltraPure | Invitrogen | 15514-029 |
| Tissue Culture | | |
| DMEM, high glucose, no glutamine, no phenol red | Thermo Fisher (Gibco) | 31053028 |

| | | |
|---|---|---|
| GlutaMAX | Thermo Fisher (Gibco) | 35050061 |
| Sodium pyruvate, 100 mM | Thermo Fisher (Gibco) | 11360070 |
| HEPES, 1 M | Thermo Fisher (Gibco) | 15630080 |
| Fetal bovine serum, qualified | Thermo Fisher (Gibco) | 26140-079 |
| TrypLE express dissociation reagent | Thermo Fisher (Gibco) | 12604-013 |
| Phosphate buffered saline | Corning | 21-040-CV |
| Bovine fibronectin, liquid, suitable for cell culture | Sigma-Aldrich | F1141 |
| Opti-MEM Reduced Serum Media | Thermo Fisher (Gibco) | 31985062 |
| TransIT LT1 Transfection Reagent | Mirus Bio | MIR2300 |
| #1.5 Chambered Coverglass, 8 well | CellVis | C8-1.5H-N |
| #1.5 Chambered Coverglass, 18 well | CellVis | C18-1.5H |
| 384 Well glass bottom plate with high performance #1.5 cover glass | CellVis | P384-1.5H-N |
| Western Blotting & Immunoprecipitation | | |
| Pierce RIPA Lysis and Extraction Buffer | Life Technologies | 89900 |
| Pierce IP Lysis Buffer | Life Technologies | 87787 |
| Halt Protease Inhibitor Cocktail (100X) | Life Technologies | 87786 |
| Pierce Bovine Serum Albumin Standard | Life Technologies | 23208 |
| Novex Tris-glycine 4-12% gradient gel | Thermo Fisher Scientific | XP04125BOX |
| Trans-Blot Turbo 0.2 µm PVDF Transfer Packs | Bio-Rad | 1704156 |
| DC Protein Assay Kit | Bio-Rad | 5000111 |
| Beta-Mercaptoethanol | MP Biomedicals | CAS 60-24-2 |
| 2x Tris-glycine SDS Loading Buffer | Life Technologies | LC2676 |
| 10x Tris-glycine SDS Running Buffer | Life Technologies | LC2675 |
| Anti-DYKDDDDK (FLAG) Monoclonal antibody, FG4R (mouse) | Life Technologies | MA191878 |
| GAPDH Loading Control Monoclonal Antibody, GA1R (mouse) | Life Technologies | MA515738 |
| V5 Tag Monoclonal Antibody (SV5-Pk1, mouse) | Life Technologies | R96025 |
| Anti-mouse IgG, HRP-linked antibody | Cell Signaling Technology | 7076S |
| SignalFire ECL Reagent | Cell Signaling Technology | 6883S |
| 5 M NaCl | Thermo Fisher | AM9759 |
| 1 M Tris pH 7.5 | Thermo Fisher | 15567027 |
| Dry Milk Omniblock | American Bio | AB10109-010000 |
| Tween-20 | Thermo Fisher | J20605-AP |

| | | |
|---|---|---|
| Disuccinimidyl glutarate, no weigh, 10 x 1 mg | Thermo Fisher | A35392 |
| Pierce 660nm Protein Assay Reagent | Life Technologies | 22660 |
| Anti-FLAG M2 Affinity Gel | Thermo Fisher | A2220 |
| Pierce Silver Stain Kit | Life Technologies | 24612 |
| Micro Bio-Spin Chromatography Columns | BioRad | 7326204 |
| 4-20% Criterion TGX Precast Midi Protein Gel | BioRad | 5671025 |
| IRDye 680RD Donkey anti-Mouse IgG Secondary antibody | LI-COR | 926-68072 |
| IRDye 800CW Donkey anti-Rabbit IgG Secondary antibody | LI-COR | 926-32213 |
| E6AP/UBE3A (D10D3) Rabbit Monoclonal Antibody | Cell Signaling Technology | 7526 |
| Compounds and Cell Treatments | | |
| Rapamycin | Fisher Scientific | CAS 53123-88-9 |
| Carfilzomib | MedChemExpress | CAS 868540-17-4 |
| TAK-243 | MedChemExpress | CAS 1450833-55-2 |

**Supplementary Table 3. Reagents and Resources.**

**Supplementary Table 4. Optical components of the triplet tumbling instrument**

| Label in **Supplementary Fig. 1** | Description | Source | Part Number |
|---|---|---|---|
| Dichroic beamsplitters and emission filters | | | |
| 925LP | 925 nm edge BrightLine multiphoton single edge dichroic beamsplitter | Semrock | FF925-Di01 |
| 830LP | 830 nm laser BrightLine single-edge laser-flat dichroic beamsplitter | Semrock | Di02-R830 |
| 785LP | 785 nm laser BrightLine single-edge super-resolution TIRF dichroic beamsplitter | Semrock | Di03-R785 |
| 750SP | 749 nm edge BrightLine single-edge short-pass image-splitting dichroic beamsplitter | Semrock | FF749-SDi01 |
| 488LP | 488 nm Long Pass dichroic mirror, UltraFlat, 2.0 waves/inch, P-V, 24mm x 34mm with Cut Corners For Nikon Eclipse x 1mm | Chroma | ZT488rdc-UF1 |
| EM1 | 500 nm longpass TIRF filter, mounted in 25mm ring | Chroma | ET500lp-25mmR |
| EM2 | IR cleanup filter, 750 nm blocking edge BrightLine multiphoton short pass emission filter | Semrock | FF01-750/SP-25 |
| EM3 | IR cleanup filter, 785 nm RazorEdge ultrasteep short pass edge filter | Semrock | SP01-785RU-25 |
| ND | Automated filter wheel containing neutral density filters ranging from ND 0 to ND 8 to modulate 488 nm excitation intensity | Sutter Instruments, ThorLabs | |
| | First excitation cleanup filter for LED | ThorLabs | FESH500 |
| | Second excitation cleanup filter for LED | Chroma | ET470/40 |
| Polarization | | | |
| HWP1 | Ø1/2" Zero-Order Half-Wave Plate, Ø1" Mount, 780 nm | ThorLabs | WPH05M-780 |
| HWP2 | Ø1/2" Zero-Order Half-Wave Plate, Ø1" Mount, 830 nm | ThorLabs | WPH05M-830 |
| HWP3 | Ø1/2" Zero-Order Half-Wave Plate, Ø1" Mount, 915 nm | ThorLabs | WPH05M-915 |
| HWP4 | 1/2" x 1/2" Mounted Superachromatic Half-Wave Plate, Ø1" Mount, 310 - 1100 nm | ThorLabs | SAHWP05M-700 |
| HWP5 | Ø1/2" Zero-Order Half-Wave Plate, Ø1" Mount, 488 nm | ThorLabs | WPH05M-488 |

| | | | |
|---|---|---|---|
| PBS | Polarizing beam splitter cube, Vis coated | ThorLabs | CCM1-PBS251/M |
| Mirrors and Lenses | | | |
| M1 | Elliptical, protected silver coated mirror | ThorLabs | PFE10-P01 |
| M2 | Knife-edge right angle prism protected silver mirror | ThorLabs | MRAK25-P01 |
| MI | Dielectric coated mirrors (IR coating -E03) | ThorLabs | BB1-E03 or BB2-E03 |
| MV | Dielectric coated mirrors (Vis coating -E02) | ThorLabs | BB1-E02 or BB2-E02 |
| FM | Flipper mirror. Automated flipper holding a -E02 dielectric coated mirror. | Thorlabs | MFF101/M |
| TL1 | f=200 mm, infinity corrected tube lens | ThorLabs | ITL200 |
| TL2 | f=500 mm, ∅2” Achromatic doublet | ThorLabs | ACT508-500-A-ML |
| L1 | Ø1" N-BK7 Plano-Convex Lens, f = 40 mm, Anti-reflective coating 400-1100 nm | ThorLabs | LA1422-AB-ML |
| L2 | Ø1" N-BK7 Plano-Convex Lens, f = 150 mm, Anti-reflective coating 400-1100 nm | ThorLabs | LA1433-AB-ML |
| L3 | Ø1" N-BK7 Plano-Convex Lens, f = 60 mm, Anti-reflective coating 400-1100 nm | ThorLabs | LA1134-AB-ML |
| L4 | Ø1" N-BK7 Plano-Convex Lens, f = 125 mm, Anti-reflective coating 400-1100 nm | ThorLabs | LA1986-AB-ML |
| Obj | 25X Nikon CFI APO LWD Objective, 380-1050 nm, 1.10 NA, 2.0 mm WD | ThorLabs | N25X-APO-MP |
| Light sources | | | |
| LED | M490L3 diode controlled by LEDD1B. Beam shaping with a telescope of AC254-050-A-ML and AC254-200-A-ML | ThorLabs | various |
| 488 nm | Cobolt 06-MLD 488 nm 200 mW laser head | Hubner Photonics | |
| 785 nm | Cobolt 06-MLD 785 nm 250 mW laser head | Hubner Photonics | |
| 830 nm | Cobolt 06-MLD 830 nm 250 mW laser head | Hubner Photonics | |
| 915 nm | Cobolt 06-MLD 915 nm 250 mW laser head | Hubner Photonics | |
| 940 nm | Cobolt 06-MLD 940 nm 250 mW laser head | Hubner Photonics | |
| Detectors | | | |

| | | | |
|---|---|---|---|
| sCMOS | PCO panda 4.2 sCMOS camera | Excelitas Technologies | pco.panda 4.2 bi USB |
| Intensified camera | PCO dicam C1 intensified camera with 18 mm S20 intensifier and P46 phosphor | Excelitas Technologies | pco.dicam C1 UHS 18-S20-P46 sel |

**Supplementary Table 4. Optical components of the triplet tumbling instrument.**

**Supplementary Table 5. Power delivered to the sample.**

| Wavelength (nm) | Nominal power output (mW) | Measured CW power at the sample (mW) | Maximum energy per 30 ns pulse (nJ)* | Average power during an acquisition (mW) |
|---|---|---|---|---|
| 488 | 40 | 39 | 1.2 | 0.04 |
| 785 | 250 | 180 | 5.4 | 157 |
| 830 | 250 | 125 | 3.8 | 133 |
| 915 | 250 | 108 | 3.2 | 109 |
| 940 | 250 | 98 | 2.9 | 100 |

**Supplementary Table 5. Power delivered to the sample.**

For all measurements, the laser was focused to an approximately 8 µm diameter ($1/e^2$) spot. The average power during an acquisition was measured with the 25 µs cycle period used for purified protein and cell measurements. Almost all the infrared intensity is delivered during the "clean up" pulse rather than the trigger pulse (20 µs duration vs. 30 ns duration).

*These calculations assume that the laser comes to full power instantaneously, which is not accurate. Empirically, measured with a fast photodiode (Thorlabs DET10A/M), the lasers take 20 to 30 ns to ramp to full power; this ramp is approximately linear. Therefore, these values provide an upper bound on the energy in a 30 ns pulse.

**Amino acid sequences of constructs used in this work**

*Purified fluorescent proteins*
Constructs were cloned into the pRSET B backbone.

6xHis-mStayGold
MRGSHHHHHHGMASTSMVSTGEELFTGVVPFKFQLKGTINGKSFTVEGEGEGNSHEGSHKGKYV
CTSGKLPMSWAALGTSFGYGMKYYTKYPSGLKNWFHEVMPEGFTYDRHIQYKGDGSIHAKHQHF
MKNGTYHNIVEFTGQDFKENSPVLTGDMDVSLPNEVQHIPIDDGVECTVTLQYPLLSDESKCVE
AYQNTIIKPLHNQPAPDVPFHWIRKQYTQSKDDTEERDHIIQSETLEAHL*

6xHis-mBaoJin
MRGSHHHHHHGMASTSMVSKGEEENMASTPFKFQLKGTINGKSFTVEGEGEGNSHEGSHKGKYV
CTSGKLPMSWAALGTTFGYGMKYYTKYPSGLKNWFREVMPGGFTYDRHIQYKGDGSIHAKHQHF
MKNGTYHNIVEFTGQDFKENSPVLTGDMNVSLPNEVPQIPRDDGVECPVTLLYPLLSDKSKYVE
AHQYTICKPLHNQPAPDVPYHWIRKQYTQSKDDAEERDHICQSETLEAHLKGMDELYK*

6xHis-mVenus
MRGSHHHHHHGMASMVSKGEELFTGVVPILVELDGDVNGHKFSVSGEGEGDATYGKLTLKLICT
TGKLPVPWPTLVTTLGYGLQCFARYPDHMKQHDFFKSAMPEGYVQERTIFFKDDGNYKTRAEVK
FEGDTLVNRIELKGIDFKEDGNILGHKLEYNYNSHNVYITADKQKNGIKANFKIRHNIEDGGVQ
LADHYQQNTPIGDGPVLLPDNHYLSYQSKLSKDPNEKRDHMVLLEFVTAAGITLGMDELYK*

6xHis-mNeonGreen-Strep
MRGSHHHHHHGGSGGSMVSKGEEDNMASLPATHELHIFGSINGVDFDMVGQGTGNPNDGYEELN
LKSTKGDLQFSPWILVPHIGYGFHQYLPYPDGMSPFQAAMVDGSGYQVHRTMQFEDGASLTVNY
RYTYEGSHIKGEAQVKGTGFPADGPVMTNSLTAADWCRSKKTYPNDKTIISTFKWSYTTGNGKR
YRSTARTTYTFAKPMAANYLKNQPMYVFRKTELKHSKTELNFKEWQKAFTDVMGMDELYKGTSW
SHPQFEK*

6xHis-mEGFP
MRGSHHHHHHGMASMVSKGEELFTGVVPILVELDGDVNGHKFSVSGEGEGDATYGKLTLKFICT
TGKLPVPWPTLVTTLTYGVQCFSRYPDHMKQHDFFKSAMPEGYVQERTIFFKDDGNYKTRAEVK
FEGDTLVNRIELKGIDFKEDGNILGHKLEYNYNSHNVYIMADKQKNGIKVNFKIRHNIEDGSVQ
LADHYQQNTPIGDGPVLLPDNHYLSTQSKLSKDPNEKRDHMVLLEFVTAAGITLGMDELYK*

6xHis-Clover
MRGSHHHHHHGMASTSMVSKGEELFTGVVPILVELDGDVNGHKFSVRGEGEGDATNGKLTLKFI
CTTGKLPVPWPTLVTTFGYGVACFSRYPDHMKQHDFFKSAMPEGYVQERTISFKDDGTYKTRAE
VKFEGDTLVNRIELKGIDFKEDGNILGHKLEYNFNSHNVYITADKQKNGIKANFKIRHNVEDGS
VQLADHYQQNTPIGDGPVLLPDNHYLSHQSALSKDPNEKRDHMVLLEFVTAAGITHGMDELYK*

*Purified protein: rapamycin dimerization*
Constructs were cloned into the pRSET B backbone.

6xHis-FKBP-Strep
MRGSHHHHHHGGSGGSMGVQVETISPGDGRTFPKRGQTCVVHYTGMLEDGKKFDSSRDRNKPFK
FMLGKQEVIRGWEEGVAQMSVGQRAKLTISPDYAYGATGHPGIIPPHATLVFDVELLKLEGTSG
ASWSHPQFEK*

6xHis-FRB-Strep
MRGSHHHHHHGGSGGSRVAILWHEMWHEGLEEASRLYFGERNVKGMFEVLEPLHAMMERGPQTL
KETSFNQAYGRDLMEAQEWCRKYMKSGNVKDLTQAWDLYYHVFRRISKQGTSGASWSHPQFEK*

6xHis-SMT3(missing last 4 aas)-FKBP-Strep
MRGSHHHHHHGGSGGSDSEVNQEAKPEVKPEVKPETHINLKVSDGSSEIFFKIKKTTPLRRLME
AFAKRQGKEMDSLRFLYDGIRIQADQAPEDLDMEDNDIIEAHREMGVQVETISPGDGRTFPKRG
QTCVVHYTGMLEDGKKFDSSRDRNKPFKFMLGKQEVIRGWEEGVAQMSVGQRAKLTISPDYAYG
ATGHPGIIPPHATLVFDVELLKLEGTSGASWSHPQFEK*

6xHis-SMT3(missing last 4 aas)-FRB-Strep
MRGSHHHHHHGGSGGSDSEVNQEAKPEVKPEVKPETHINLKVSDGSSEIFFKIKKTTPLRRLME
AFAKRQGKEMDSLRFLYDGIRIQADQAPEDLDMEDNDIIEAHRERVAILWHEMWHEGLEEASRL
YFGERNVKGMFEVLEPLHAMMERGPQTLKETSFNQAYGRDLMEAQEWCRKYMKSGNVKDLTQAW
DLYYHVFRRISKQGTSGASWSHPQFEK*

6xHis-MBP-FKBP-Strep
MRGSHHHHHHGGSGGSKTEEGKLVIWINGDKGYNGLAEVGKKFEKDTGIKVTVEHPDKLEEKFP
QVAATGDGPDIIFWAHDRFGGYAQSGLLAEITPDKAFQDKLYPFTWDAVRYNGKLIAYPIAVEA
LSLIYNKDLLPNPPKTWEEIPALDKELKAKGKSALMFNLQEPYFTWPLIAADGGYAFKYENGKY
DIKDVGVDNAGAKAGLTFLVDLIKNKHMNADTDYSIAEHAFNHGETAMTINGPWAWSNIDTSKV
NYGVTVLPTFKGQPSKPFVGVLSAGINAASPNKELAKEFLENYLLTDEGLEAVNKDKPLGAVAL
KSYEEELVKDPRVAATMENAQKGEIMPNIPQMSAFWYAVRTAVINAASGRQTVDEALKDAQTMG
VQVETISPGDGRTFPKRGQTCVVHYTGMLEDGKKFDSSRDRNKPFKFMLGKQEVIRGWEEGVAQ
MSVGQRAKLTISPDYAYGATGHPGIIPPHATLVFDVELLKLEGTSWSHPQFEK*

6xHis-MBP-FRB-Strep
MRGSHHHHHHGGSGGSKTEEGKLVIWINGDKGYNGLAEVGKKFEKDTGIKVTVEHPDKLEEKFP
QVAATGDGPDIIFWAHDRFGGYAQSGLLAEITPDKAFQDKLYPFTWDAVRYNGKLIAYPIAVEA
LSLIYNKDLLPNPPKTWEEIPALDKELKAKGKSALMFNLQEPYFTWPLIAADGGYAFKYENGKY
DIKDVGVDNAGAKAGLTFLVDLIKNKHMNADTDYSIAEHAFNHGETAMTINGPWAWSNIDTSKV
NYGVTVLPTFKGQPSKPFVGVLSAGINAASPNKELAKEFLENYLLTDEGLEAVNKDKPLGAVAL
KSYEEELVKDPRVAATMENAQKGEIMPNIPQMSAFWYAVRTAVINAASGRQTVDEALKDAQTRV
AILWHEMWHEGLEEASRLYFGERNVKGMFEVLEPLHAMMERGPQTLKETSFNQAYGRDLMEAQE
WCRKYMKSGNVKDLTQAWDLYYHVFRRISKQGTSWSHPQFEK*

6xHis-MBP-pepN-FKBP-Strep
MRGSHHHHHHGGSGGSKTEEGKLVIWINGDKGYNGLAEVGKKFEKDTGIKVTVEHPDKLEEKFP
QVAATGDGPDIIFWAHDRFGGYAQSGLLAEITPDKAFQDKLYPFTWDAVRYNGKLIAYPIAVEA

LSLIYNKDLLPNPPKTWEEIPALDKELKAKGKSALMFNLQEPYFTWPLIAADGGYAFKYENGKY
DIKDVGVDNAGAKAGLTFLVDLIKNKHMNADTDYSIAEHAFNHGETAMTINGPWAWSNIDTSKV
NYGVTVLPTFKGQPSKPFVGVLSAGINAASPNKELAKEFLENYLLTDEGLEAVNKDKPLGAVAL
KSYEEELVKDPRVAATMENAQKGEIMPNIPQMSAFWYAVRTAVINAASGRQTVDEALKDAQTTQ
QPQAKYRHDYRAPDYQITDIDLTFDLDAQKTVVTAVSQAVRHGASDAPLRLNGEDLKLVSVHIN
DEPWTAWKEEEGALVISNLPERFTLKIINEISPAANTALEGLYQSGDALCTQCEAEGFRHITYY
LDRPDVLARFTTKIIADKIKYPFLLSNGNRVAQGELENGRHWVQWQDPFPKPCYLFALVAGDFD
VLRDTFTTRSGREVALELYVDRGNLDRAPWAMTSLKNSMKWDEERFGLEYDLDIYMIVAVDFFN
MGAMENKGLNIFNSKYVLARTDTATDKDYLDIERVIGHEYFHNWTGNRVTCRDWFQLSLKEGLT
VFRDQEFSSDLGSRAVNRINNVRTMRGLQFAEDASPMAHPIRPDMVIEMNNFYTLTVYEKGAEV
IRMIHTLLGEENFQKGMQLYFERHDGSAATCDDFVQAMEDASNVDLSHFRRWYSQSGTPIVTVK
DDYNPETEQYTLTISQRTPATPDQAEKQPLHIPFAIELYDNEGKVIPLQKGGHPVNSVLNVTQA
EQTFVFDNVYFQPVPALLCEFSAPVKLEYKWSDQQLTFLMRHARNDFSRWDAAQSLLATYIKLN
VARHQQGQPLSLPVHVADAFRAVLLDEKIDPALAAEILTLPSVNEMAELFDIIDPIAIAEVREA
LTRTLATELADELLAIYNANYQSEYRVEHEDIAKRTLRNACLRFLAFGETHLADVLVSKQFHEA
NNMTDALAALSAAVAAQLPCRDALMQEYDDKWHQNGLVMDKWFILQATSPAANVLETVRGLLQH
RSFTMSNPNRIRSLIGAFAGSNPAAFHAEDGSGYLFLVEMLTDLNSRNPQVASRLIEPLIRLKR
YDAKRQEKMRAALEQLKGLENLSGDLYEKITKALAMGVQVETISPGDGRTFPKRGQTCVVHYTG
MLEDGKKFDSSRDRNKPFKFMLGKQEVIRGWEEGVAQMSVGQRAKLTISPDYAYGATGHPGIIP
PHATLVFDVELLKLEGTSWSHPQFEK*

6xHis-MBP-pepN-FRB-Strep

MRGSHHHHHHGGSGGSKTEEGKLVIWINGDKGYNGLAEVGKKFEKDTGIKVTVEHPDKLEEKFP
QVAATGDGPDIIFWAHDRFGGYAQSGLLAEITPDKAFQDKLYPFTWDAVRYNGKLIAYPIAVEA
LSLIYNKDLLPNPPKTWEEIPALDKELKAKGKSALMFNLQEPYFTWPLIAADGGYAFKYENGKY
DIKDVGVDNAGAKAGLTFLVDLIKNKHMNADTDYSIAEHAFNHGETAMTINGPWAWSNIDTSKV
NYGVTVLPTFKGQPSKPFVGVLSAGINAASPNKELAKEFLENYLLTDEGLEAVNKDKPLGAVAL
KSYEEELVKDPRVAATMENAQKGEIMPNIPQMSAFWYAVRTAVINAASGRQTVDEALKDAQTTQ
QPQAKYRHDYRAPDYQITDIDLTFDLDAQKTVVTAVSQAVRHGASDAPLRLNGEDLKLVSVHIN
DEPWTAWKEEEGALVISNLPERFTLKIINEISPAANTALEGLYQSGDALCTQCEAEGFRHITYY
LDRPDVLARFTTKIIADKIKYPFLLSNGNRVAQGELENGRHWVQWQDPFPKPCYLFALVAGDFD
VLRDTFTTRSGREVALELYVDRGNLDRAPWAMTSLKNSMKWDEERFGLEYDLDIYMIVAVDFFN
MGAMENKGLNIFNSKYVLARTDTATDKDYLDIERVIGHEYFHNWTGNRVTCRDWFQLSLKEGLT
VFRDQEFSSDLGSRAVNRINNVRTMRGLQFAEDASPMAHPIRPDMVIEMNNFYTLTVYEKGAEV
IRMIHTLLGEENFQKGMQLYFERHDGSAATCDDFVQAMEDASNVDLSHFRRWYSQSGTPIVTVK
DDYNPETEQYTLTISQRTPATPDQAEKQPLHIPFAIELYDNEGKVIPLQKGGHPVNSVLNVTQA
EQTFVFDNVYFQPVPALLCEFSAPVKLEYKWSDQQLTFLMRHARNDFSRWDAAQSLLATYIKLN
VARHQQGQPLSLPVHVADAFRAVLLDEKIDPALAAEILTLPSVNEMAELFDIIDPIAIAEVREA
LTRTLATELADELLAIYNANYQSEYRVEHEDIAKRTLRNACLRFLAFGETHLADVLVSKQFHEA
NNMTDALAALSAAVAAQLPCRDALMQEYDDKWHQNGLVMDKWFILQATSPAANVLETVRGLLQH
RSFTMSNPNRIRSLIGAFAGSNPAAFHAEDGSGYLFLVEMLTDLNSRNPQVASRLIEPLIRLKR
YDAKRQEKMRAALEQLKGLENLSGDLYEKITKALARVAILWHEMWHEGLEEASRLYFGERNVKG
MFEVLEPLHAMMERGPQTLKETSFNQAYGRDLMEAQEWCRKYMKSGNVKDLTQAWDLYYHVFRR
ISKQGTSGASWSHPQFEK*

6xHis-mVenus-FKBP-Strep

MRGSHHHHHHGGSGGSMVSKGEELFTGVVPILVELDGDVNGHKFSVSGEGEGDATYGKLTLKLI
CTTGKLPVPWPTLVTTLGYGLQCFARYPDHMKQHDFFKSAMPEGYVQERTIFFKDDGNYKTRAE
VKFEGDTLVNRIELKGIDFKEDGNILGHKLEYNYNSHNVYITADKQKNGIKANFKIRHNIEDGG
VQLADHYQQNTPIGDGPVLLPDNHYLSYQSKLSKDPNEKRDHMVLLEFVTAAGITLGMDELYKM
GVQVETISPGDGRTFPKRGQTCVVHYTGMLEDGKKFDSSRDRNKPFKFMLGKQEVIRGWEEGVA
QMSVGQRAKLTISPDYAYGATGHPGIIPPHATLVFDVELLKLEGTSWSHPQFEK*

6xHis-mVenusΔC7-FKBP-Strep

MRGSHHHHHHGGSGGSMVSKGEELFTGVVPILVELDGDVNGHKFSVSGEGEGDATYGKLTLKLI
CTTGKLPVPWPTLVTTLGYGLQCFARYPDHMKQHDFFKSAMPEGYVQERTIFFKDDGNYKTRAE
VKFEGDTLVNRIELKGIDFKEDGNILGHKLEYNYNSHNVYITADKQKNGIKANFKIRHNIEDGG
VQLADHYQQNTPIGDGPVLLPDNHYLSYQSKLSKDPNEKRDHMVLLEFVTAAGITLMGVQVETI
SPGDGRTFPKRGQTCVVHYTGMLEDGKKFDSSRDRNKPFKFMLGKQEVIRGWEEGVAQMSVGQR
AKLTISPDYAYGATGHPGIIPPHATLVFDVELLKLEGTSWSHPQFEK*

6xHis-mVenusΔC9-FKBP-Strep

MRGSHHHHHHGGSGGSMVSKGEELFTGVVPILVELDGDVNGHKFSVSGEGEGDATYGKLTLKLI
CTTGKLPVPWPTLVTTLGYGLQCFARYPDHMKQHDFFKSAMPEGYVQERTIFFKDDGNYKTRAE
VKFEGDTLVNRIELKGIDFKEDGNILGHKLEYNYNSHNVYITADKQKNGIKANFKIRHNIEDGG
VQLADHYQQNTPIGDGPVLLPDNHYLSYQSKLSKDPNEKRDHMVLLEFVTAAGIMGVQVETISP
GDGRTFPKRGQTCVVHYTGMLEDGKKFDSSRDRNKPFKFMLGKQEVIRGWEEGVAQMSVGQRAK
LTISPDYAYGATGHPGIIPPHATLVFDVELLKLEGTSWSHPQFEK*

6xHis-mVenusΔC10-FKBP-Strep

MRGSHHHHHHGGSGGSMVSKGEELFTGVVPILVELDGDVNGHKFSVSGEGEGDATYGKLTLKLI
CTTGKLPVPWPTLVTTLGYGLQCFARYPDHMKQHDFFKSAMPEGYVQERTIFFKDDGNYKTRAE
VKFEGDTLVNRIELKGIDFKEDGNILGHKLEYNYNSHNVYITADKQKNGIKANFKIRHNIEDGG
VQLADHYQQNTPIGDGPVLLPDNHYLSYQSKLSKDPNEKRDHMVLLEFVTAAGMGVQVETISPG
DGRTFPKRGQTCVVHYTGMLEDGKKFDSSRDRNKPFKFMLGKQEVIRGWEEGVAQMSVGQRAKL
TISPDYAYGATGHPGIIPPHATLVFDVELLKLEGTSGASWSHPQFEK*

6xHis-mVenusΔC11-FKBP-Strep

MRGSHHHHHHGGSGGSMVSKGEELFTGVVPILVELDGDVNGHKFSVSGEGEGDATYGKLTLKLI
CTTGKLPVPWPTLVTTLGYGLQCFARYPDHMKQHDFFKSAMPEGYVQERTIFFKDDGNYKTRAE
VKFEGDTLVNRIELKGIDFKEDGNILGHKLEYNYNSHNVYITADKQKNGIKANFKIRHNIEDGG
VQLADHYQQNTPIGDGPVLLPDNHYLSYQSKLSKDPNEKRDHMVLLEFVTAAMGVQVETISPGD
GRTFPKRGQTCVVHYTGMLEDGKKFDSSRDRNKPFKFMLGKQEVIRGWEEGVAQMSVGQRAKLT
ISPDYAYGATGHPGIIPPHATLVFDVELLKLEGTSGASWSHPQFEK*

6xHis-mVenusΔC12-FKBP-Strep

MRGSHHHHHHGGSGGSMVSKGEELFTGVVPILVELDGDVNGHKFSVSGEGEGDATYGKLTLKLI
CTTGKLPVPWPTLVTTLGYGLQCFARYPDHMKQHDFFKSAMPEGYVQERTIFFKDDGNYKTRAE
VKFEGDTLVNRIELKGIDFKEDGNILGHKLEYNYNSHNVYITADKQKNGIKANFKIRHNIEDGG
VQLADHYQQNTPIGDGPVLLPDNHYLSYQSKLSKDPNEKRDHMVLLEFVTAMGVQVETISPGDG
RTFPKRGQTCVVHYTGMLEDGKKFDSSRDRNKPFKFMLGKQEVIRGWEEGVAQMSVGQRAKLTI
SPDYAYGATGHPGIIPPHATLVFDVELLKLEGTSGASWSHPQFEK*

6xHis-mVenusΔC13-FKBP-Strep

MRGSHHHHHHGGSGGSMVSKGEELFTGVVPILVELDGDVNGHKFSVSGEGEGDATYGKLTLKLI
CTTGKLPVPWPTLVTTLGYGLQCFARYPDHMKQHDFFKSAMPEGYVQERTIFFKDDGNYKTRAE
VKFEGDTLVNRIELKGIDFKEDGNILGHKLEYNYNSHNVYITADKQKNGIKANFKIRHNIEDGG
VQLADHYQQNTPIGDGPVLLPDNHYLSYQSKLSKDPNEKRDHMVLLEFVTMGVQVETISPGDGR
TFPKRGQTCVVHYTGMLEDGKKFDSSRDRNKPFKFMLGKQEVIRGWEEGVAQMSVGQRAKLTIS
PDYAYGATGHPGIIPPHATLVFDVELLKLEGTSGASWSHPQFEK*

6xHis-mVenusΔC11-FRB-Strep

MRGSHHHHHHGGSGGSMVSKGEELFTGVVPILVELDGDVNGHKFSVSGEGEGDATYGKLTLKLI
CTTGKLPVPWPTLVTTLGYGLQCFARYPDHMKQHDFFKSAMPEGYVQERTIFFKDDGNYKTRAE
VKFEGDTLVNRIELKGIDFKEDGNILGHKLEYNYNSHNVYITADKQKNGIKANFKIRHNIEDGG
VQLADHYQQNTPIGDGPVLLPDNHYLSYQSKLSKDPNEKRDHMVLLEFVTAARVAILWHEMWHE
GLEEASRLYFGERNVKGMFEVLEPLHAMMERGPQTLKETSFNQAYGRDLMEAQEWCRKYMKSGN
VKDLTQAWDLYYHVFRRISKQGTSGASWSHPQFEK*

6xHis-FRB-ΔN5mVenus-Strep

MRGSHHHHHHGGSGGSRVAILWHEMWHEGLEEASRLYFGERNVKGMFEVLEPLHAMMERGPQTL
KETSFNQAYGRDLMEAQEWCRKYMKSGNVKDLTQAWDLYYHVFRRISKQEELFTGVVPILVELD
GDVNGHKFSVSGEGEGDATYGKLTLKLICTTGKLPVPWPTLVTTLGYGLQCFARYPDHMKQHDF
FKSAMPEGYVQERTIFFKDDGNYKTRAEVKFEGDTLVNRIELKGIDFKEDGNILGHKLEYNYNS
HNVYITADKQKNGIKANFKIRHNIEDGGVQLADHYQQNTPIGDGPVLLPDNHYLSYQSKLSKDP
NEKRDHMVLLEFVTAAGITLGMDELYKGTSGASWSHPQFEK*

6xHis-FRB-ΔN6mVenus-Strep

MRGSHHHHHHGGSGGSRVAILWHEMWHEGLEEASRLYFGERNVKGMFEVLEPLHAMMERGPQTL
KETSFNQAYGRDLMEAQEWCRKYMKSGNVKDLTQAWDLYYHVFRRISKQELFTGVVPILVELDG
DVNGHKFSVSGEGEGDATYGKLTLKLICTTGKLPVPWPTLVTTLGYGLQCFARYPDHMKQHDFF
KSAMPEGYVQERTIFFKDDGNYKTRAEVKFEGDTLVNRIELKGIDFKEDGNILGHKLEYNYNSH
NVYITADKQKNGIKANFKIRHNIEDGGVQLADHYQQNTPIGDGPVLLPDNHYLSYQSKLSKDPN
EKRDHMVLLEFVTAAGITLGMDELYKGTSWSHPQFEK*

6xHis-FRB-ΔN7mVenus-Strep

MRGSHHHHHHGGSGGSRVAILWHEMWHEGLEEASRLYFGERNVKGMFEVLEPLHAMMERGPQTL
KETSFNQAYGRDLMEAQEWCRKYMKSGNVKDLTQAWDLYYHVFRRISKQLFTGVVPILVELDGD
VNGHKFSVSGEGEGDATYGKLTLKLICTTGKLPVPWPTLVTTLGYGLQCFARYPDHMKQHDFFK
SAMPEGYVQERTIFFKDDGNYKTRAEVKFEGDTLVNRIELKGIDFKEDGNILGHKLEYNYNSHN
VYITADKQKNGIKANFKIRHNIEDGGVQLADHYQQNTPIGDGPVLLPDNHYLSYQSKLSKDPNE
KRDHMVLLEFVTAAGITLGMDELYKGTSGASWSHPQFEK*

6xHis-FRB-ΔN8mVenus-Strep

MRGSHHHHHHGGSGGSRVAILWHEMWHEGLEEASRLYFGERNVKGMFEVLEPLHAMMERGPQTL
KETSFNQAYGRDLMEAQEWCRKYMKSGNVKDLTQAWDLYYHVFRRISKQFTGVVPILVELDGDV
NGHKFSVSGEGEGDATYGKLTLKLICTTGKLPVPWPTLVTTLGYGLQCFARYPDHMKQHDFFKS
AMPEGYVQERTIFFKDDGNYKTRAEVKFEGDTLVNRIELKGIDFKEDGNILGHKLEYNYNSHNV

YITADKQKNGIKANFKIRHNIEDGGVQLADHYQQNTPIGDGPVLLPDNHYLSYQSKLSKDPNEK
RDHMVLLEFVTAAGITLGMDELYKGTSGASWSHPQFEK*

6xHis-FRB-ΔN9mVenus-Strep
MRGSHHHHHHGGSGGSRVAILWHEMWHEGLEEASRLYFGERNVKGMFEVLEPLHAMMERGPQTL
KETSFNQAYGRDLMEAQEWCRKYMKSGNVKDLTQAWDLYYHVFRRISKQTGVVPILVELDGDVN
GHKFSVSGEGEGDATYGKLTLKLICTTGKLPVPWPTLVTTLGYGLQCFARYPDHMKQHDFFKSA
MPEGYVQERTIFFKDDGNYKTRAEVKFEGDTLVNRIELKGIDFKEDGNILGHKLEYNYNSHNVY
ITADKQKNGIKANFKIRHNIEDGGVQLADHYQQNTPIGDGPVLLPDNHYLSYQSKLSKDPNEKR
DHMVLLEFVTAAGITLGMDELYKGTSGASWSHPQFEK*

6xHis-FRB-ΔN10mVenus-Strep
MRGSHHHHHHGGSGGSRVAILWHEMWHEGLEEASRLYFGERNVKGMFEVLEPLHAMMERGPQTL
KETSFNQAYGRDLMEAQEWCRKYMKSGNVKDLTQAWDLYYHVFRRISKQGVVPILVELDGDVNG
HKFSVSGEGEGDATYGKLTLKLICTTGKLPVPWPTLVTTLGYGLQCFARYPDHMKQHDFFKSAM
PEGYVQERTIFFKDDGNYKTRAEVKFEGDTLVNRIELKGIDFKEDGNILGHKLEYNYNSHNVYI
TADKQKNGIKANFKIRHNIEDGGVQLADHYQQNTPIGDGPVLLPDNHYLSYQSKLSKDPNEKRD
HMVLLEFVTAAGITLGMDELYKGTSGASWSHPQFEK*

6xHis-FRB-ΔN11mVenus-Strep
MRGSHHHHHHGGSGGSRVAILWHEMWHEGLEEASRLYFGERNVKGMFEVLEPLHAMMERGPQTL
KETSFNQAYGRDLMEAQEWCRKYMKSGNVKDLTQAWDLYYHVFRRISKQVVPILVELDGDVNGH
KFSVSGEGEGDATYGKLTLKLICTTGKLPVPWPTLVTTLGYGLQCFARYPDHMKQHDFFKSAMP
EGYVQERTIFFKDDGNYKTRAEVKFEGDTLVNRIELKGIDFKEDGNILGHKLEYNYNSHNVYIT
ADKQKNGIKANFKIRHNIEDGGVQLADHYQQNTPIGDGPVLLPDNHYLSYQSKLSKDPNEKRDH
MVLLEFVTAAGITLGMDELYKGTSGASWSHPQFEK*

6xHis-mStayGoldΔC2-FKBP-Strep
MRGSHHHHHHGGSGGSVSTGEELFTGVVPFKFQLKGTINGKSFTVEGEGEGNSHEGSHKGKYVC
TSGKLPMSWAALGTSFGYGMKYYTKYPSGLKNWFHEVMPEGFTYDRHIQYKGDGSIHAKHQHFM
KNGTYHNIVEFTGQDFKENSPVLTGDMDVSLPNEVQHIPIDDGVECTVTLQYPLLSDESKCVEA
YQNTIIKPLHNQPAPDVPFHWIRKQYTQSKDDTEERDHIIQSETLEAMGVQVETISPGDGRTFP
KRGQTCVVHYTGMLEDGKKFDSSRDRNKPFKFMLGKQEVIRGWEEGVAQMSVGQRAKLTISPDY
AYGATGHPGIIPPHATLVFDVELLKLEGTSGASWSHPQFEK*

*Mammalian expression: rapamycin dimerization*
Constructs were cloned into the pcDNA3.1(+) backbone.

FLAG-FRB-IRES-mVenusΔC12-FKBP
ORF1:
MDYKDDDDKRVAILWHEMWHEGLEEASRLYFGERNVKGMFEVLEPLHAMMERGPQTLKETSFNQ
AYGRDLMEAQEWCRKYMKSGNVKDLTQAWDLYYHVFRRISKQ*
ORF2:
MVSKGEELFTGVVPILVELDGDVNGHKFSVSGEGEGDATYGKLTLKLICTTGKLPVPWPTLVTT
LGYGLQCFARYPDHMKQHDFFKSAMPEGYVQERTIFFKDDGNYKTRAEVKFEGDTLVNRIELKG
IDFKEDGNILGHKLEYNYNSHNVYITADKQKNGIKANFKIRHNIEDGGVQLADHYQQNTPIGDG
PVLLPDNHYLSYQSKLSKDPNEKRDHMVLLEFVTAMGVQVETISPGDGRTFPKRGQTCVVHYTG
MLEDGKKFDSSRDRNKPFKFMLGKQEVIRGWEEGVAQMSVGQRAKLTISPDYAYGATGHPGIIP
PHATLVFDVELLKLE*

FLAG-SMT3(missing last 4 aas)-FRB-IRES-mVenusΔC12-FKBP
ORF1:
MDYKDDDDKDSEVNQEAKPEVKPEVKPETHINLKVSDGSSEIFFKIKKTTPLRRLMEAFAKRQG
KEMDSLRFLYDGIRIQADQAPEDLDMEDNDIIEAHRERVAILWHEMWHEGLEEASRLYFGERNV
KGMFEVLEPLHAMMERGPQTLKETSFNQAYGRDLMEAQEWCRKYMKSGNVKDLTQAWDLYYHVF
RRISKQ*
ORF2:
MVSKGEELFTGVVPILVELDGDVNGHKFSVSGEGEGDATYGKLTLKLICTTGKLPVPWPTLVTT
LGYGLQCFARYPDHMKQHDFFKSAMPEGYVQERTIFFKDDGNYKTRAEVKFEGDTLVNRIELKG
IDFKEDGNILGHKLEYNYNSHNVYITADKQKNGIKANFKIRHNIEDGGVQLADHYQQNTPIGDG
PVLLPDNHYLSYQSKLSKDPNEKRDHMVLLEFVTAMGVQVETISPGDGRTFPKRGQTCVVHYTG
MLEDGKKFDSSRDRNKPFKFMLGKQEVIRGWEEGVAQMSVGQRAKLTISPDYAYGATGHPGIIP
PHATLVFDVELLKLE*

FLAG-MBP-FRB-IRES-mVenusΔC12-FKBP
ORF1:
MDYKDDDDKKTEEGKLVIWINGDKGYNGLAEVGKKFEKDTGIKVTVEHPDKLEEKFPQVAATGD
GPDIIFWAHDRFGGYAQSGLLAEITPDKAFQDKLYPFTWDAVRYNGKLIAYPIAVEALSLIYNK
DLLPNPPKTWEEIPALDKELKAKGKSALMFNLQEPYFTWPLIAADGGYAFKYENGKYDIKDVGV
DNAGAKAGLTFLVDLIKNKHMNADTDYSIAEHAFNHGETAMTINGPWAWSNIDTSKVNYGVTVL
PTFKGQPSKPFVGVLSAGINAASPNKELAKEFLENYLLTDEGLEAVNKDKPLGAVALKSYEEEL
VKDPRVAATMENAQKGEIMPNIPQMSAFWYAVRTAVINAASGRQTVDEALKDAQTRVAILWHEM
WHEGLEEASRLYFGERNVKGMFEVLEPLHAMMERGPQTLKETSFNQAYGRDLMEAQEWCRKYMK
SGNVKDLTQAWDLYYHVFRRISKQ*
ORF2:
MVSKGEELFTGVVPILVELDGDVNGHKFSVSGEGEGDATYGKLTLKLICTTGKLPVPWPTLVTT
LGYGLQCFARYPDHMKQHDFFKSAMPEGYVQERTIFFKDDGNYKTRAEVKFEGDTLVNRIELKG
IDFKEDGNILGHKLEYNYNSHNVYITADKQKNGIKANFKIRHNIEDGGVQLADHYQQNTPIGDG
PVLLPDNHYLSYQSKLSKDPNEKRDHMVLLEFVTAMGVQVETISPGDGRTFPKRGQTCVVHYTG
MLEDGKKFDSSRDRNKPFKFMLGKQEVIRGWEEGVAQMSVGQRAKLTISPDYAYGATGHPGIIP
PHATLVFDVELLKLE*

FLAG-MBP-pepN-FRB-IRES-mVenusΔC12-FKBP
ORF1:
MDYKDDDDKKTEEGKLVIWINGDKGYNGLAEVGKKFEKDTGIKVTVEHPDKLEEKFPQVAATGD
GPDIIFWAHDRFGGYAQSGLLAEITPDKAFQDKLYPFTWDAVRYNGKLIAYPIAVEALSLIYNK
DLLPNPPKTWEEIPALDKELKAKGKSALMFNLQEPYFTWPLIAADGGYAFKYENGKYDIKDVGV
DNAGAKAGLTFLVDLIKNKHMNADTDYSIAEHAFNHGETAMTINGPWAWSNIDTSKVNYGVTVL
PTFKGQPSKPFVGVLSAGINAASPNKELAKEFLENYLLTDEGLEAVNKDKPLGAVALKSYEEEL
VKDPRVAATMENAQKGEIMPNIPQMSAFWYAVRTAVINAASGRQTVDEALKDAQTTQQPQAKYR
HDYRAPDYQITDIDLTFDLDAQKTVVTAVSQAVRHGASDAPLRLNGEDLKLVSVHINDEPWTAW
KEEEGALVISNLPERFTLKIINEISPAANTALEGLYQSGDALCTQCEAEGFRHITYYLDRPDVL
ARFTTKIIADKIKYPFLLSNGNRVAQGELENGRHWVQWQDPFPKPCYLFALVAGDFDVLRDTFT
TRSGREVALELYVDRGNLDRAPWAMTSLKNSMKWDEERFGLEYDLDIYMIVAVDFFNMGAMENK
GLNIFNSKYVLARTDTATDKDYLDIERVIGHEYFHNWTGNRVTCRDWFQLSLKEGLTVFRDQEF
SSDLGSRAVNRINNVRTMRGLQFAEDASPMAHPIRPDMVIEMNNFYTLTVYEKGAEVIRMIHTL
LGEENFQKGMQLYFERHDGSAATCDDFVQAMEDASNVDLSHFRRWYSQSGTPIVTVKDDYNPET
EQYTLTISQRTPATPDQAEKQPLHIPFAIELYDNEGKVIPLQKGGHPVNSVLNVTQAEQTFVFD
NVYFQPVPALLCEFSAPVKLEYKWSDQQLTFLMRHARNDFSRWDAAQSLLATYIKLNVARHQQG
QPLSLPVHVADAFRAVLLDEKIDPALAAEILTLPSVNEMAELFDIIDPIAIAEVREALTRTLAT
ELADELLAIYNANYQSEYRVEHEDIAKRTLRNACLRFLAFGETHLADVLVSKQFHEANNMTDAL
AALSAAVAAQLPCRDALMQEYDDKWHQNGLVMDKWFILQATSPAANVLETVRGLLQHRSFTMSN
PNRIRSLIGAFAGSNPAAFHAEDGSGYLFLVEMLTDLNSRNPQVASRLIEPLIRLKRYDAKRQE
KMRAALEQLKGLENLSGDLYEKITKALARVAILWHEMWHEGLEEASRLYFGERNVKGMFEVLEP
LHAMMERGPQTLKETSFNQAYGRDLMEAQEWCRKYMKSGNVKDLTQAWDLYYHVFRRISKQ*
ORF2:
MVSKGEELFTGVVPILVELDGDVNGHKFSVSGEGEGDATYGKLTLKLICTTGKLPVPWPTLVTT
LGYGLQCFARYPDHMKQHDFFKSAMPEGYVQERTIFFKDDGNYKTRAEVKFEGDTLVNRIELKG
IDFKEDGNILGHKLEYNYNSHNVYITADKQKNGIKANFKIRHNIEDGGVQLADHYQQNTPIGDG
PVLLPDNHYLSYQSKLSKDPNEKRDHMVLLEFVTAMGVQVETISPGDGRTFPKRGQTCVVHYTG
MLEDGKKFDSSRDRNKPFKFMLGKQEVIRGWEEGVAQMSVGQRAKLTISPDYAYGATGHPGIIP
PHATLVFDVELLKLE*

FLAG-FRB-IRES-mStayGoldΔC2-FKBP
ORF1:
MDYKDDDDKRVAILWHEMWHEGLEEASRLYFGERNVKGMFEVLEPLHAMMERGPQTLKETSFNQ
AYGRDLMEAQEWCRKYMKSGNVKDLTQAWDLYYHVFRRISKQ*
ORF2:
MVSTGEELFTGVVPFKFQLKGTINGKSFTVEGEGEGNSHEGSHKGKYVCTSGKLPMSWAALGTS
FGYGMKYYTKYPSGLKNWFHEVMPEGFTYDRHIQYKGDGSIHAKHQHFMKNGTYHNIVEFTGQD
FKENSPVLTGDMDVSLPNEVQHIPIDDGVECTVTLQYPLLSDESKCVEAYQNTIIKPLHNQPAP
DVPFHWIRKQYTQSKDDTEERDHIIQSETLEAMGVQVETISPGDGRTFPKRGQTCVVHYTGMLE
DGKKFDSSRDRNKPFKFMLGKQEVIRGWEEGVAQMSVGQRAKLTISPDYAYGATGHPGIIPPHA
TLVFDVELLKLE*

FLAG-SMT3(missing last 4 aas)-FRB-IRES-mStayGoldΔC2-FKBP
ORF1:
MDYKDDDDKDSEVNQEAKPEVKPEVKPETHINLKVSDGSSEIFFKIKKTTPLRRLMEAFAKRQG
KEMDSLRFLYDGIRIQADQAPEDLDMEDNDIIEAHRERVAILWHEMWHEGLEEASRLYFGERNV

KGMFEVLEPLHAMMERGPQTLKETSFNQAYGRDLMEAQEWCRKYMKSGNVKDLTQAWDLYYHVF
RRISKQ*

ORF2:

MVSTGEELFTGVVPFKFQLKGTINGKSFTVEGEGEGNSHEGSHKGKYVCTSGKLPMSWAALGTS
FGYGMKYYTKYPSGLKNWFHEVMPEGFTYDRHIQYKGDGSIHAKHQHFMKNGTYHNIVEFTGQD
FKENSPVLTGDMDVSLPNEVQHIPIDDGVECTVTLQYPLLSDESKCVEAYQNTIIKPLHNQPAP
DVPFHWIRKQYTQSKDDTEERDHIIQSETLEAMGVQVETISPGDGRTFPKRGQTCVVHYTGMLE
DGKKFDSSRDRNKPFKFMLGKQEVIRGWEEGVAQMSVGQRAKLTISPDYAYGATGHPGIIPPHA
TLVFDVELLKLE*

FLAG-MBP-FRB-IRES-mStayGoldΔC2-FKBP

ORF1:

MDYKDDDDKKTEEGKLVIWINGDKGYNGLAEVGKKFEKDTGIKVTVEHPDKLEEKFPQVAATGD
GPDIIFWAHDRFGGYAQSGLLAEITPDKAFQDKLYPFTWDAVRYNGKLIAYPIAVEALSLIYNK
DLLPNPPKTWEEIPALDKELKAKGKSALMFNLQEPYFTWPLIAADGGYAFKYENGKYDIKDVGV
DNAGAKAGLTFLVDLIKNKHMNADTDYSIAEHAFNHGETAMTINGPWAWSNIDTSKVNYGVTVL
PTFKGQPSKPFVGVLSAGINAASPNKELAKEFLENYLLTDEGLEAVNKDKPLGAVALKSYEEEL
VKDPRVAATMENAQKGEIMPNIPQMSAFWYAVRTAVINAASGRQTVDEALKDAQTRVAILWHEM
WHEGLEEASRLYFGERNVKGMFEVLEPLHAMMERGPQTLKETSFNQAYGRDLMEAQEWCRKYMK
SGNVKDLTQAWDLYYHVFRRISKQ*

ORF2:

MVSTGEELFTGVVPFKFQLKGTINGKSFTVEGEGEGNSHEGSHKGKYVCTSGKLPMSWAALGTS
FGYGMKYYTKYPSGLKNWFHEVMPEGFTYDRHIQYKGDGSIHAKHQHFMKNGTYHNIVEFTGQD
FKENSPVLTGDMDVSLPNEVQHIPIDDGVECTVTLQYPLLSDESKCVEAYQNTIIKPLHNQPAP
DVPFHWIRKQYTQSKDDTEERDHIIQSETLEAMGVQVETISPGDGRTFPKRGQTCVVHYTGMLE
DGKKFDSSRDRNKPFKFMLGKQEVIRGWEEGVAQMSVGQRAKLTISPDYAYGATGHPGIIPPHA
TLVFDVELLKLE*

FLAG-MBP-pepN-FRB-IRES-mStayGoldΔC2-FKBP

ORF1:

MDYKDDDDKKTEEGKLVIWINGDKGYNGLAEVGKKFEKDTGIKVTVEHPDKLEEKFPQVAATGD
GPDIIFWAHDRFGGYAQSGLLAEITPDKAFQDKLYPFTWDAVRYNGKLIAYPIAVEALSLIYNK
DLLPNPPKTWEEIPALDKELKAKGKSALMFNLQEPYFTWPLIAADGGYAFKYENGKYDIKDVGV
DNAGAKAGLTFLVDLIKNKHMNADTDYSIAEHAFNHGETAMTINGPWAWSNIDTSKVNYGVTVL
PTFKGQPSKPFVGVLSAGINAASPNKELAKEFLENYLLTDEGLEAVNKDKPLGAVALKSYEEEL
VKDPRVAATMENAQKGEIMPNIPQMSAFWYAVRTAVINAASGRQTVDEALKDAQTTQQPQAKYR
HDYRAPDYQITDIDLTFDLDAQKTVVTAVSQAVRHGASDAPLRLNGEDLKLVSVHINDEPWTAW
KEEEGALVISNLPERFTLKIINEISPAANTALEGLYQSGDALCTQCEAEGFRHITYYLDRPDVL
ARFTTKIIADKIKYPFLLSNGNRVAQGELENGRHWVQWQDPFPKPCYLFALVAGDFDVLRDTFT
TRSGREVALELYVDRGNLDRAPWAMTSLKNSMKWDEERFGLEYDLDIYMIVAVDFFNMGAMENK
GLNIFNSKYVLARTDTATDKDYLDIERVIGHEYFHNWTGNRVTCRDWFQLSLKEGLTVFRDQEF
SSDLGSRAVNRINNVRTMRGLQFAEDASPMAHPIRPDMVIEMNNFYTLTVYEKGAEVIRMIHTL
LGEENFQKGMQLYFERHDGSAATCDDFVQAMEDASNVDLSHFRRWYSQSGTPIVTVKDDYNPET
EQYTLTISQRTPATPDQAEKQPLHIPFAIELYDNEGKVIPLQKGGHPVNSVLNVTQAEQTFVFD
NVYFQPVPALLCEFSAPVKLEYKWSDQQLTFLMRHARNDFSRWDAAQSLLATYIKLNVARHQQG
QPLSLPVHVADAFRAVLLDEKIDPALAAEILTLPSVNEMAELFDIIDPIAIAEVREALTRTLAT
ELADELLAIYNANYQSEYRVEHEDIAKRTLRNACLRFLAFGETHLADVLVSKQFHEANNMTDAL

AALSAAVAAQLPCRDALMQEYDDKWHQNGLVMDKWFILQATSPAANVLETVRGLLQHRSFTMSN
PNRIRSLIGAFAGSNPAAFHAEDGSGYLFLVEMLTDLNSRNPQVASRLIEPLIRLKRYDAKRQE
KMRAALEQLKGLENLSGDLYEKITKALARVAILWHEMWHEGLEEASRLYFGERNVKGMFEVLEP
LHAMMERGPQTLKETSFNQAYGRDLMEAQEWCRKYMKSGNVKDLTQAWDLYYHVFRRISKQ*

ORF2:
MVSTGEELFTGVVPFKFQLKGTINGKSFTVEGEGEGNSHEGSHKGKYVCTSGKLPMSWAALGTS
FGYGMKYYTKYPSGLKNWFHEVMPEGFTYDRHIQYKGDGSIHAKHQHFMKNGTYHNIVEFTGQD
FKENSPVLTGDMDVSLPNEVQHIPIDDGVECTVTLQYPLLSDESKCVEAYQNTIIKPLHNQPAP
DVPFHWIRKQYTQSKDDTEERDHIIQSETLEAMGVQVETISPGDGRTFPKRGQTCVVHYTGMLE
DGKKFDSSRDRNKPFKFMLGKQEVIRGWEEGVAQMSVGQRAKLTISPDYAYGATGHPGIIPPHA
TLVFDVELLKLE*

*Mammalian expression: p53*

Note that the n1 linker nomenclature refers to the linkers originally tested to improve mStayGold expression.[25] Constructs were cloned into the pcDNA3.1(+) backbone.

p53-Δ6n1-mStayGold-FLAG (full length p53)

MEEPQSDPSVEPPLSQETFSDLWKLLPENNVLSPLPSQAMDDLMLSPDDIEQWFTEDPGPDEAP
RMPEAAPRVAPAPAAPTPAAPAPAPSWPLSSSVPSQKTYQGSYGFRLGFLHSGTAKSVTCTYSP
ALNKMFCQLAKTCPVQLWVDSTPPPGTRVRAMAIYKQSQHMTEVVRRCPHHERCSDSDGLAPPQ
HLIRVEGNLRVEYLDDRNTFRHSVVVPYEPPEVGSDCTTIHYNYMCNSSCMGGMNRRPILTIIT
LEDSSGNLLGRNSFEVRVCACPGRDRRTEEENLRKKGEPHHELPPGSTKRALPNNTSSSPQPKK
KPLDGEYFTLQIRGRERFEMFRELNEALELKDAQAGKEPGGSRAHSSHLKSKKGQSTSRHKKLM
FKTEGPDSDELFTGVVPFKFQLKGTINGKSFTVEGEGEGNSHEGSHKGKYVCTSGKLPMSWAAL
GTSFGYGMKYYTKYPSGLKNWFHEVMPEGFTYDRHIQYKGDGSIHAKHQHFMKNGTYHNIVEFT
GQDFKENSPVLTGDMDVSLPNEVQHIPIDDGVECTVTLQYPLLSDESKCVEAYQNTIIKPLHNQ
PAPDVPFHWIRKQYTQSKDDTEERDHIIQSETLEAHLDYKDDDDK*

p53ΔCreg-Δ6n1-mStayGold-FLAG

MEEPQSDPSVEPPLSQETFSDLWKLLPENNVLSPLPSQAMDDLMLSPDDIEQWFTEDPGPDEAP
RMPEAAPRVAPAPAAPTPAAPAPAPSWPLSSSVPSQKTYQGSYGFRLGFLHSGTAKSVTCTYSP
ALNKMFCQLAKTCPVQLWVDSTPPPGTRVRAMAIYKQSQHMTEVVRRCPHHERCSDSDGLAPPQ
HLIRVEGNLRVEYLDDRNTFRHSVVVPYEPPEVGSDCTTIHYNYMCNSSCMGGMNRRPILTIIT
LEDSSGNLLGRNSFEVRVCACPGRDRRTEEENLRKKGEPHHELPPGSTKRALPNNTSSSPQPKK
KPLDGEYFTLQIRGRERFEMFRELNEALELKDAQAGKEPGGSRELFTGVVPFKFQLKGTINGKS
FTVEGEGEGNSHEGSHKGKYVCTSGKLPMSWAALGTSFGYGMKYYTKYPSGLKNWFHEVMPEGF
TYDRHIQYKGDGSIHAKHQHFMKNGTYHNIVEFTGQDFKENSPVLTGDMDVSLPNEVQHIPIDD
GVECTVTLQYPLLSDESKCVEAYQNTIIKPLHNQPAPDVPFHWIRKQYTQSKDDTEERDHIIQS
ETLEAHLDYKDDDDK*

*Mammalian expression: E6AP + HPV16 E6*

Constructs were expressed either from pcDNA3.1(+) or a sleeping beauty transposase backbone from VectorBuilder. In both cases, E6AP expression was driven by the CMV promoter and E6 expression was driven by the EF1a promoter. The polyA sequence used for E6AP was bGH. The E6 polyA sequence was SV40 in the transposase backbone and rBG in the pcDNA constructs.

E6AP- Δ6n1-mStayGold-FLAG

```
MEKLHQCYWKSGEPQSDDIEASRMKRAAAKHLIERYYHQLTEGCGNEACTNEFCASCPTFLRMD
NNAAAIKALELYKINAKLCDPHPSKKGASSAYLENSKGAPNNSCSEIKMNKKGARIDFKDVTYL
TEEKVYEILELCREREDYSPLIRVIGRVFSSAEALVQSFRKVKQHTKEELKSLQAKDEDKDEDE
KEKAACSAAAMEEDSEASSSRIGDSSQGDNNLQKLGPDDVSVDIDAIRRVYTRLLSNEKIETAF
LNALVYLSPNVECDLTYHNVYSRDPNYLNLFIIVMENRNLHSPEYLEMALPLFCKAMSKLPLAA
QGKLIRLWSKYNADQIRRMMETFQQLITYKVISNEFNSRNLVNDDDAIVAASKCLKMVYYANVV
GGEVDTNHNEEDDEEPIPESSELTLQELLGEERRNKKGPRVDPLETELGVKTLDCRKPLIPFEE
FINEPLNEVLEMDKDYTFFKVETENKFSFMTCPFILNAVTKNLGLYYDNRIRMYSERRITVLYS
LVQGQQLNPYLRLKVRRDHIIDDALVRLEMIAMENPADLKKQLYVEFEGEQGVDEGGVSKEFFQ
LVVEEIFNPDIGMFTYDESTKLFWFNPSSFETEGQFTLIGIVLGLAIYNNCILDVHFPMVVYRK
LMGKKGTFRDLGDSHPVLYQSLKDLLEYEGNVEDDMMITFQISQTDLFGNPMMYDLKENGDKIP
ITNENRKEFVNLYSDYILNKSVEKQFKAFRRGFHMVTNESPLKYLFRPEEIELLICGSRNLDFQ
ALEETTEYDGGYTRDSVLIREFWEIVHSFTDEQKRLFLQFTTGTDRAPVGGLGKLKMIIAKNGP
DTERLPTSHTCFNVLLLPEYSSKEKLKERLLKAITYAKGFGMLELFTGVVPFKFQLKGTINGKS
FTVEGEGEGNSHEGSHKGKYVCTSGKLPMSWAALGTSFGYGMKYYTKYPSGLKNWFHEVMPEGF
TYDRHIQYKGDGSIHAKHQHFMKNGTYHNIVEFTGQDFKENSPVLTGDMDVSLPNEVQHIPIDD
GVECTVTLQYPLLSDESKCVEAYQNTIIKPLHNQPAPDVPFHWIRKQYTQSKDDTEERDHIIQS
ETLEAHLDYKDDDDK*
```

V5-E6

```
MGKPIPNPLLGLDSTGGHQKRTAMFQDPQERPRKLPQLCTELQTTIHDIILECVYCKQQLLRRE
VYDFAFRDLCIVYRDGNPYAVCDKCLKFYSKISEYRHYCYSVYGTTLEQQYNKPLCDLLIRCIN
CQKPLCPEEKQRHLDKKQRFHNIRGRWTGRCMSCCRSSRTRRETQL*
```

**Supplementary References**


1. Lakowicz, J. R. *Principles of Fluorescence Spectroscopy, Third Edition*. (Springer, 2006).

2. Virtanen, P. *et al.* SciPy 1.0: fundamental algorithms for scientific computing in Python. *Nat. Methods* **17**, 261–272 (2020).

3. Hurvich, C. M. & Tsai, C.-L. Regression and time series model selection in small samples. *Biometrika* **76**, 297–307 (1989).

4. Schwarz, G. Estimating the Dimension of a Model. *The Annals of Statistics* **6**, 461 (1978).

5. Lu, Y.-H. *et al.* Sequential Two-Photon Delayed Fluorescence Anisotropy for Macromolecular Size Determination. *J. Phys. Chem. B* **127**, 3861–3869 (2023).

6. Lazzari-Dean, J. R. *et al.* From cameras to confocal to cytometry: measuring tumbling rates is a general way to reveal protein binding. *Zenodo* https://doi.org/10.5281/zenodo.10028432 (2023).

7. Kremers, G.-J., Goedhart, J., Munster, E. B. van & Gadella, T. W. J. Cyan and Yellow Super Fluorescent Proteins with Improved Brightness, Protein Folding, and FRET Förster Radius † , ‡. *Biochemistry* **45**, 6570–6580 (2006).

8. Ando, R. *et al.* StayGold variants for molecular fusion and membrane-targeting applications. *Nat. Methods* **21**, 648–656 (2024).

9. Zacharias, D. A., Violin, J. D., Newton, A. C. & Tsien, R. Y. Partitioning of Lipid-Modified Monomeric GFPs into Membrane Microdomains of Live Cells. *Science* **296**, 913–916 (2002).

10. Zhang, H. *et al.* Bright and stable monomeric green fluorescent protein derived from StayGold. *Nat. Methods* **21**, 657–665 (2024).

11. Shaner, N. C. *et al.* A bright monomeric green fluorescent protein derived from Branchiostoma lanceolatum. *Nat. Methods* **10**, 407–409 (2013).

12. Lam, A. J. *et al.* Improving FRET dynamic range with bright green and red fluorescent proteins. *Nat. Methods* **9**, 1005–1012 (2012).

13. Gasteiger, E. *et al. The Proteomics Protocols Handbook*. (Humana Press, 2005).

14. Swaminathan, R., Hoang, C. P. & Verkman, A. S. Photobleaching recovery and anisotropy decay of green fluorescent protein GFP-S65T in solution and cells: cytoplasmic viscosity probed by green fluorescent protein translational and rotational diffusion. *Biophys. J.* **72**, 1900–1907 (1997).

15. Williams, J. W. & Morrison, J. F. The kinetics of reversible tight-binding inhibition. *Methods Enzymol* **63**, 437–67 (1979).

16. Schneider, C. A., Rasband, W. S. & Eliceiri, K. W. NIH Image to ImageJ: 25 years of image analysis. *Nat. Methods* **9**, 671–675 (2012).

17. Kawate, T. & Gouaux, E. Fluorescence-Detection Size-Exclusion Chromatography for Precrystallization Screening of Integral Membrane Proteins. *Structure* **14**, 673–681 (2006).

18. Harris, C. R. *et al.* Array programming with NumPy. *Nature* **585**, 357–362 (2020).

19. The pandas development team. pandas-dev/pandas: Pandas (v3.0.1). *Zenodo* https://doi.org/10.5281/zenodo.3509134 (2026) doi:10.5281/zenodo.3509134.

20. Hunter, J. D. Matplotlib: A 2D Graphics Environment. *Comput. Sci. Eng.* **9**, 90–95 (2007).

21. Friedman, P. N., Chen, X., Bargonetti, J. & Prives, C. The p53 protein is an unusually shaped tetramer that binds directly to DNA. *Proc. Natl. Acad. Sci.* **90**, 3319–3323 (1993).

22. Galvis, A. E., Fisher, H. E. & Camerini, D. NP-40 Fractionation and Nucleic Acid Extraction in Mammalian Cells. *Bio-protocol* **7**, e2584 (2017).

23. Kühnle, S. *et al.* Angelman syndrome–associated point mutations in the Zn2+-binding N-terminal (AZUL) domain of UBE3A ubiquitin ligase inhibit binding to the proteasome. *J. Biol. Chem.* **293**, 18387–18399 (2018).

24. Landau, L. D. & Lifshitz, E. M. *Fluid Mechanics*. (1987).

25. Hirano, M. *et al.* A highly photostable and bright green fluorescent protein. *Nat. Biotechnol.* **40**, 1132–1142 (2022).
